# RNA-coupled CRISPR Screens Reveal ZNF207 as a Regulator of LMNA Aberrant Splicing in Progeria

**DOI:** 10.1101/2025.04.25.648738

**Authors:** Amit K. Behera, Jeongjin J. Kim, Shreya Kordale, Arun Prasath Damodaran, Bandana Kumari, Sandra Vidak, Ethan Dickson, Mei-Sheng Xiao, Gerard Duncan, Thorkell Andresson, Tom Misteli, Thomas Gonatopoulos-Pournatzis

**Affiliations:** RNA Biology Laboratory, Center for Cancer Research (CCR), National Cancer Institute (NCI), National Institutes of Health (NIH), Frederick, MD 21702, USA; Laboratory of Receptor Biology and Gene Expression, Center for Cancer Research (CCR), National Cancer Institute (NCI), National Institutes of Health (NIH), Bethesda, MD 20892, USA; Protein Characterization Laboratory, Frederick National Laboratory for Cancer Research (FNLCR), Frederick, MD 21701, USA

**Keywords:** Alternative pre-mRNA splicing, pre-mRNA processing, Hutchinson-Gilford Progeria Syndrome (HGPS), ZNF207, RNA-coupled CRISPR screen, Nonsense-mediated mRNA decay (NMD), SRSF7, PKM, BuGZ, CRASP-Seq

## Abstract

Despite progress in understanding pre-mRNA splicing, the regulatory mechanisms controlling most alternative splicing events remain unclear. We developed CRASP-Seq, a method that integrates pooled CRISPR-based genetic perturbations with deep sequencing of splicing reporters, to quantitively assess the impact of all human genes on alternative splicing from a single RNA sample. CRASP-Seq identifies both known and novel regulators, enriched for proteins involved in RNA splicing and metabolism. As proof-of-concept, CRASP-Seq analysis of an LMNA cryptic splicing event linked to progeria uncovered Z*NF*207, primarily known for mitotic spindle assembly, as a regulator of progerin splicing. ZNF207 depletion enhances canonical LMNA splicing and decreases progerin levels in patient-derived cells. High-throughput mutagenesis further showed that ZNF207’s zinc finger domain broadly impacts alternative splicing through interactions with U1 snRNP factors. These findings position ZNF207 as a U1 snRNP auxiliary factor and demonstrate the power of CRASP-Seq to uncover key regulators and domains of alternative splicing.

**Main Points:**

- CRASP-Seq: RNA-coupled CRISPR screen quantifying gene and domain impact on splicing
- Profiling of five events identified 370 genes influencing alternative splicing
- ZNF207 regulates splicing by interacting with U1 snRNP via its zinc-finger domains
- ZNF207 depletion corrects LMNA aberrant splicing causing progeria

## INTRODUCTION

Gene expression fidelity and regulatory flexibility depend on the precise coordination of pre-mRNA processing events, including capping, splicing, and RNA cleavage followed by polyadenylation. At the core of these processes lies the spliceosome, an intricate molecular machine composed of five small nuclear RNAs (snRNAs) and approximately 150 proteins. These components are organized into small nuclear ribonucleoproteins (snRNPs). The snRNP particles dynamically assemble to form the spliceosome, which catalyzes the removal of introns in a stepwise manner. During the early stages of spliceosome assembly, the U1 and U2 snRNPs recognize the 5′ and 3′ splice sites, respectively. This is followed by the recruitment of the U4/U6.U5 tri-snRNP complex, which triggers extensive rearrangements in protein composition and interactions involving snRNAs with pre-mRNA or other snRNAs. These rearrangements culminate in two sequential transesterification reactions that excise introns and produce mature transcripts.^1–4^

A key aspect of pre-mRNA processing is alternative splicing, which enables a single gene to generate multiple mRNA isoforms through the selection of different splice sites. This process can result in diverse protein-coding or regulatory outcomes, highlighting its broad biological significance.^5–8^ Alternative splicing is pervasive, affecting nearly all protein-coding genes^9,10^, and a significant fraction of alternative exons have been shown to influence measurable cellular phenotypes.^11^ Alternative splicing can occur in several forms, such as the inclusion or skipping of “cassette” exons, mutually exclusive selection of adjacent exons, usage of alternative 5′ and 3′ splice sites, and differential intron retention.

The assembly of the spliceosome at alternate splice sites is influenced by multiple factors, including RNA secondary structure, transcription elongation rates, and chromatin modifications. Arguably, RNA-binding proteins, which recognize specific cis-regulatory elements on pre-mRNA, play the most prominent role in regulating splicing.^7,12^ Additionally, several core spliceosome proteins can selectively influence subsets of alternative splicing events broadening the range of regulatory components and stages involved in alternative splicing.^13–17^

Aberrant splicing choices caused by pathogenic mutations are implicated in numerous diseases, highlighting the clinical significance of RNA processing dysregulation.^18,19^ A notable example is Hutchinson-Gilford Progeria Syndrome (HGPS), a rare genetic disorder caused by a *de novo* heterozygous 1824C>T point mutation in the *LMNA* gene, which encodes the nuclear structural proteins lamin A and C.^20,21^ The disease-causing mutation activates a cryptic splice donor site within LMNA, leading to the production of progerin, a defective variant of lamin A lacking 50 crucial amino acids. Progerin exerts dominant-negative effects, causing a range of cellular defects, including disrupted nuclear architecture, alterations in epigenetic modifications and changes to various nuclear proteins, ultimately leading to patient death typically in early adolescence due to cardiovascular complications.^20,21^ It is thus important to determine the splicing regulatory factors that affect progerin aberrant splicing.

Dysregulated expression of spliced isoforms is also causally linked to numerous other diseases and disorders, including cancer.^22,23^ For example, cancer cells exhibit widespread splicing alterations^24–27^, such as those affecting the mutually exclusive exons in *PKM*^28,29^ and the cassette exons in *FAS*^30^, *EZH2*^31^, and *SRSF7*^32^, among many others. While these splicing events have been studied to varying degrees, the regulatory factors and pathways controlling most disease-associated splicing events remain largely unknown. This lack of knowledge is largely due to the absence of easy-to-implement and cost-effective experimental tools capable of providing highly quantitative insights into the pathways and factors that govern biologically relevant exons and introns.

To address this challenge, we developed CRASP-Seq, an innovative methodology that integrates deep sequencing of splicing reporters with genome-wide CRISPR-based genetic perturbations. This approach directly captures splice junctions, enabling quantitative assessment of how the knockout of each human gene influences a specific splicing event. Unlike arrayed screens, CRASP-Seq requires only a single RNA extraction from a pooled, transduced cell population, providing a cost-effective and accessible methodology. Applying this high-throughput tool to the study of factors influencing *LMNA* aberrant splicing identified ZNF207 as a positive regulator of progerin expression. Depletion of ZNF207 in HGPS-derived fibroblasts corrects *LMNA* aberrant splicing and reduces progerin protein expression. ZNF207 regulates a broad spectrum of alternative splicing events and directly binds its splicing targets. By combining CRASP-Seq with base editor-mediated high-throughput mutagenesis, we identified a point mutation within the β-sheet of its second zinc finger domain that disrupts ZNF207’s ability to regulate splicing and abolishes its interaction with U1 snRNP components.

## RESULTS

### Establishing a robust screening platform for quantifying splicing regulation

We sought to develop a scalable and quantitative platform to comprehensively study alternative splicing regulation. To do so, we integrated the expression of minigene reporters within genome-wide lentiviral CRISPR (Clustered Regularly Interspaced Short Palindromic Repeats) gene knockout libraries. This approach allows us to link alternative splicing changes of an inducible minigene reporter to the expression of specific guide (g)RNAs, thereby connecting individual gene knockouts with quantitative splicing inclusion levels (**Figure 1A**).

**Figure 1:**
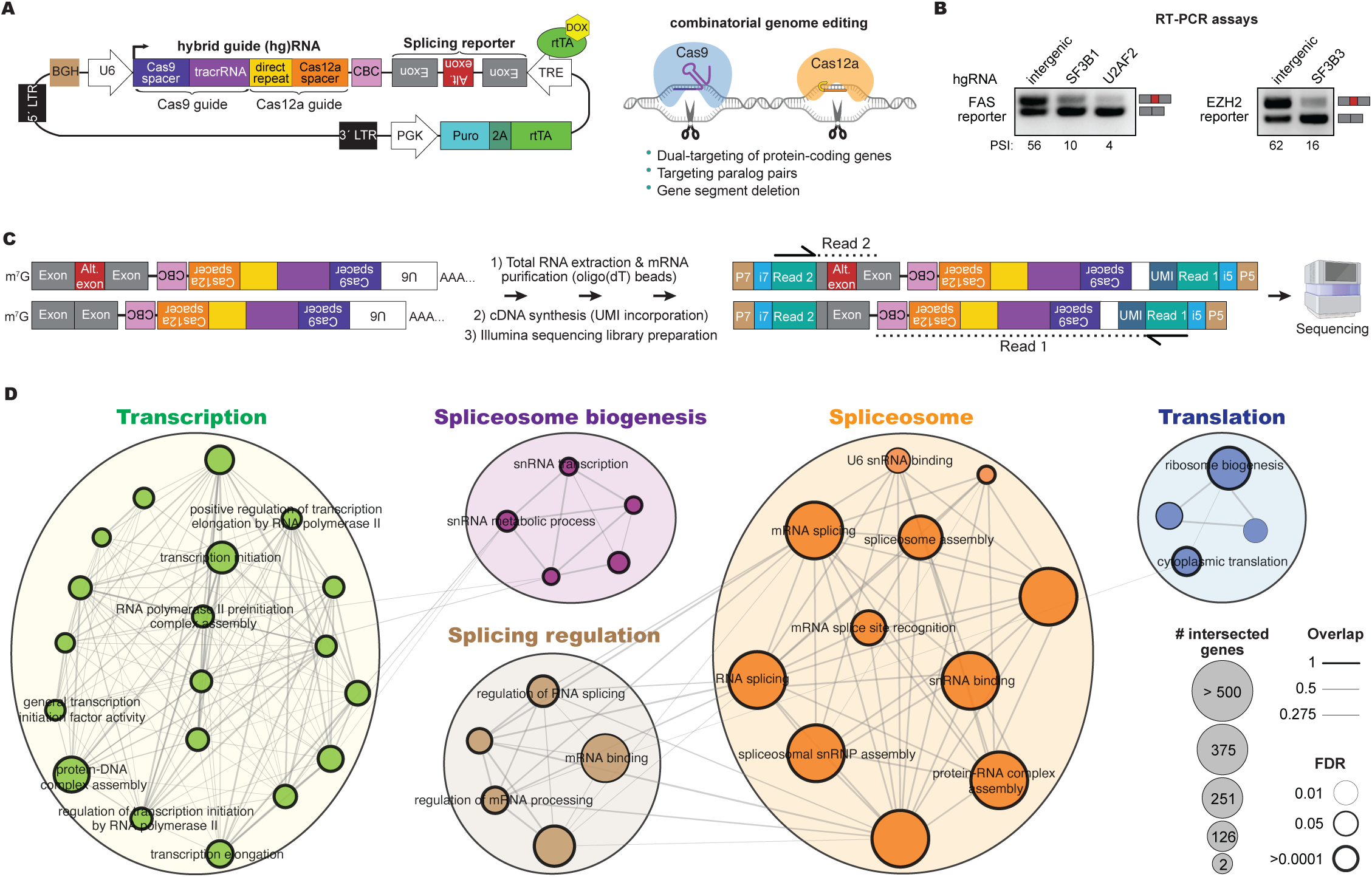
Overview of the CRASP-Seq platform for identifying alternative splicing regulators. **(A)** Schematic of the CRASP-Seq vector, which integrates constitutive Cas9 and Cas12a hgRNA expression under a U6 promoter for genome editing. A doxycycline (DOX)-inducible TRE (Tetracycline Response Element) promoter controls the expression of a splicing minigene reporter. The reverse tetracycline transactivator (rtTA) is fused to a puromycin resistance gene via a T2A cleavage sequence, allowing selectable, inducible reporter expression. An RNA polymerase II termination signal ensures efficient expression. BGH: bovine growth hormone polyadenylation signal. **(B)** RT-PCR validation of alternative splicing outcomes for *FAS* exon-6 and *EZH2* exon-14 using CRASP-Seq vectors in HAP1 cells co-expressing the indicated hgRNAs. **(C)** Workflow of the CRASP-Seq platform, illustrating the generation of splicing-derived mRNA products and the subsequent preparation of Illumina libraries for paired-end sequencing. This approach enables the precise association of each hgRNA within the library with quantified splicing outcomes. **(D)** Gene Ontology (GO) enrichment analysis of regulators identified by CRASP-Seq for the *FAS* and *EZH2* splicing events. Only significantly enriched biological processes are shown. All the terms are listed in Figure S1G.

To achieve efficient gene inactivation, we utilized our previously developed combinatorial CRISPR screening tool, CHyMErA.^11,33–35^ CHyMErA co-expresses Cas9 and Cas12a nucleases alongside libraries of hybrid guide (hg)RNAs, which are engineered by fusing Cas9 and Cas12a gRNAs transcribed from a single constitutive U6 promoter (**Figure 1A**).^33,34^ This platform enables combinatorial genetic perturbations using both Cas9 and Cas12a guides. We previously demonstrated that CHyMErA can achieve ultra-efficient gene knockout in human cells through dual targeting of individual genes with Cas9 and Cas12a gRNAs, surpassing the performance of conventional Cas9 screening platforms.^11,33^ Additionally, CHyMErA allows us to target paralogous gene pairs and other genetic elements of interest.^11,33,34^ To this end, we computationally designed a genome-wide library of 95,893 hgRNAs, targeting 18,888 protein-coding genes with four distinct hgRNAs per gene. The hgRNAs were designed to broadly target major transcript isoforms expressed across diverse cell and tissue types (Methods). Additionally, 430 intergenic and non-targeting control hgRNAs were included. The library also encompasses 2,357 paralogous gene pairs, each targeted with four unique hgRNAs (**Table S1 and Methods**). The oligo library pool was cloned into a lentiviral vector, which includes a unique cell barcode (CBC) to track splicing levels within single-cell lineages and to enable the creation of multiple replicates (**Figure 1A**; Methods).

To enable expression of the minigene reporters following CRISPR perturbations, we engineered our lentiviral vectors to include a minimal doxycycline-inducible tetracycline response element (TRE), an RNA polymerase II transcription termination sequence (bovine growth hormone [BGH] polyadenylation signal), and the open reading frame of rtTA (reverse tetracycline-controlled transactivator) fused to the puromycin resistance gene, separated by a 2A peptide cleavage sequence. These modifications enable controlled expression of the splicing minigene reporters upon doxycycline induction (**Figures 1A and S1A**). Notably, the expression cassette of minigene reporters of interest was cloned into the minus strand of the lentiviral vector, ensuring that the viral genome remains unaffected by splicing and polyadenylation during virus production and packaging in HEK293T cells. In contrast, the hgRNA expression cassette is oriented on the plus strand to prevent the Cas12a nuclease from recognizing and processing the Cas12a direct repeat (DR) within the minigene transcript, which would otherwise lead to endonucleolytic cleavage and degradation of the transcript.

To validate the system, we designed minigene reporters to monitor the splicing of *FAS* exon-6 and *EZH2* exon-14, representing well-studied (*FAS* exon-6) and less characterized (*EZH2* exon-14) alternative cassette exons in terms of their regulatory mechanisms. Misregulation of both exons has been linked to cancer.^30,31,36^ These minigene reporters, like all others used in this study, contain the full native intronic sequences flanked by adjacent constitutive exons (**Table S2**).

We first cloned the *FAS* and *EZH2* minigene splicing reporters into lentiviral vectors expressing either intergenic control hgRNAs or hgRNAs targeting established regulators of these splicing events, specifically *SF3B1* and *U2AF2* for FAS^15^, and *SF3B3* for EZH2.^31^ We then performed RT-PCR assays to assess splicing outcomes from these constructs. As anticipated, CRISPR-mediated knockout of the respective splicing factors induces significant exon skipping in both reporters (**Figure 1B**), confirming the robustness of our perturbation-readout methodology.

Next, we cloned the *FAS* exon-6 and *EZH2* exon-14 minigene reporters into our genome-wide knockout libraries (Methods). The minigene reporter libraries were used to produce lentivirus and transduce human HAP1 and RPE1 cells expressing *Streptococcus pyogenes* (*Sp*)Cas9 and optimized *Acidaminococcus sp.* (op)Cas12a nucleases^11,37^ at a low multiplicity of infection (MOI), ensuring at least 200-fold coverage. After puromycin selection and programmable editing guided by constitutively expressed hgRNAs, we induced reporter expression by treating with doxycycline for 24 hours at two different time points (harvesting at four or seven days after transduction). The resulting reporter transcripts are spliced according to the cellular genotypes, which are influenced by the expressed hgRNAs.

Our workflow quantitatively assesses splicing levels by amplifying and sequencing reporter mRNAs linked to hgRNAs. To achieve this, the hgRNA expression cassette is strategically positioned at the 3′ end of the reporter in the reverse complement orientation (**Figure 1C**). Total RNA extracted from cells transduced with the hgRNA genome-wide knockout lentiviral library is used to isolate mature poly(A)-tailed mRNA using oligodT. Subsequently, cDNA is synthesized using customized primers with unique molecular identifiers (UMI), enabling the removal of PCR duplicates during downstream analysis, and is then used to generate Illumina sequencing libraries. Paired-end sequencing enables precise quantification of splicing reporter phenotypes from read 2, while read 1 captures the corresponding hgRNA sequences to identify specific genetic perturbations (**Figure 1C**). This approach thus allows a genome-wide investigation of splicing regulators, by directly linking functional RNA splicing outcomes—quantified using the percent spliced-in (PSI) index—with specific genetic perturbations associated with each hgRNA in the library.

We initially compared the early (i.e., T1 or four days after transduction) and late (i.e., T4 or seven days after transduction) data points. Despite a strong positive correlation between the two time points, common essential genes exhibit a notably lower correlation coefficient (**Figure S1B**). Additionally, we observed a marked reduction in the number of identified gene hits at the later time point (**Figure S1C**). This reduction is accompanied by a significant depletion of core-essential gene hits (p = 0.0092; Fisher’s exact test; **Figure S1C**). Gene ontology analysis of hits specifically lost at the late time point (i.e., T1-specific hits) reveals a pronounced enrichment of spliceosome-related genes, along with a lower-level detection of other essential cellular machineries (**Figure S1D**). Based on these findings, we identified four days post-transduction (i.e., T1) as the optimal time point for inducing the minigene reporter and performing our experiments, as it ensures efficient detection of the regulatory roles of essential genes.

*FAS* exon-6 has been extensively studied, which guided its selection during the optimization of our experimental pipeline.^15^ Our screen shows a significant overlap of identified regulators with a previous genome-wide siRNA screen monitoring FAS exon-6 inclusion (p-value <0.00001; odds ratio = 3.34; Fisher’s exact test; **Figure S1E**), and an overall positive correlation between these two orthogonal screening methodologies (**Figure S1F**), despite differences in gene perturbation methods and cell lines. Furthermore, gene ontology analysis of the identified hits reveals a strong enrichment for terms related to RNA splicing and spliceosome biogenesis, further validating the effectiveness of our approach (**Figures 1D and S1G**).

We were initially surprised to observe that one of the enriched terms in the gene ontology analysis involves translation-related factors. This finding suggests that the reporter transcripts might be potential targets of the nonsense-mediated decay (NMD) pathway, which is translation-dependent. Closer examination of the reporter transcripts revealed that the minus strand of the U6 promoter contains canonical splice sites, resulting in the excision of a 139-nucleotide cryptic intron (**Figure S2A**), as previously described.^38^ Since this intron is located downstream of the reporter’s stop codon, we hypothesized that it may trigger NMD. Consistent with this hypothesis, treatment of cells with cycloheximide—a translation inhibitor that also suppresses NMD—results in increased detection of spliced transcripts, indicating that the reporter is indeed subject to NMD (**Figure S2B**). To address this issue, we mutated the 3′ splice site in the reporter, successfully preventing splicing of the U6 promoter (**Figures S2A-B**) without compromising the functionality of our CRISPR vectors (**Figure S2C**). We then re-cloned the CRISPR genome-wide knockout library, CBCs, as well as the FAS and EZH2 minigene reporters into this modified vector and repeated the assay. While the correlation between the two screens remained strongly positive (**Figure S2D**), the previously observed strong enrichment of translation-related factors was largely eliminated (**Figure S2E**). Consequently, for the remaining screens described in this manuscript, we utilized the CRISPR library backbone that is not a natural subject to NMD.

Taken together, these data demonstrate that we have developed and optimized a cost-effective and user-friendly technology for sequencing-based, genome-wide identification of regulators for splicing events of interest. We have named this tool CRASP-Seq, which stands for <u>CR</u>ISPR-based identification of <u>A</u>lternative <u>S</u>plicing regulators with <u>P</u>henotypic <u>Seq</u>uencing. CRASP-Seq seamlessly integrates deep sequencing of splicing reporters with genome-wide genetic perturbations, enabling a comprehensive and quantitative assessment of the impact of every protein-coding human gene on alternative splicing events of interest—all from a single RNA extraction sample.

### Application of CRASP-Seq to disease-relevant splicing events

To assess the versatility of CRASP-Seq across diverse alternative splicing events, we cloned three additional splicing reporters into genome-wide lentiviral vectors designed for CRASP-Seq. These reporters enable the monitoring of various alternative splicing patterns, including (1) the mutually exclusive inclusion of exon-9 or exon-10 in the *PKM* gene, which directs cells towards distinct metabolic pathways^28,39,40^; (2) a poison cassette exon in *SRSF7*, which introduces a premature stop codon, triggering NMD and playing a role in autoregulation^41,42^; and (3) *LMNA* exon-11, cloned both as the wild-type sequence and with a synonymous point mutation that activates an aberrant 5′ splice site, leading to progerin production—a protein causing Hutchinson-Gilford Progeria Syndrome (HGPS)^20,21^ (**Figure 2A; Table S2**).

**Figure 2:**
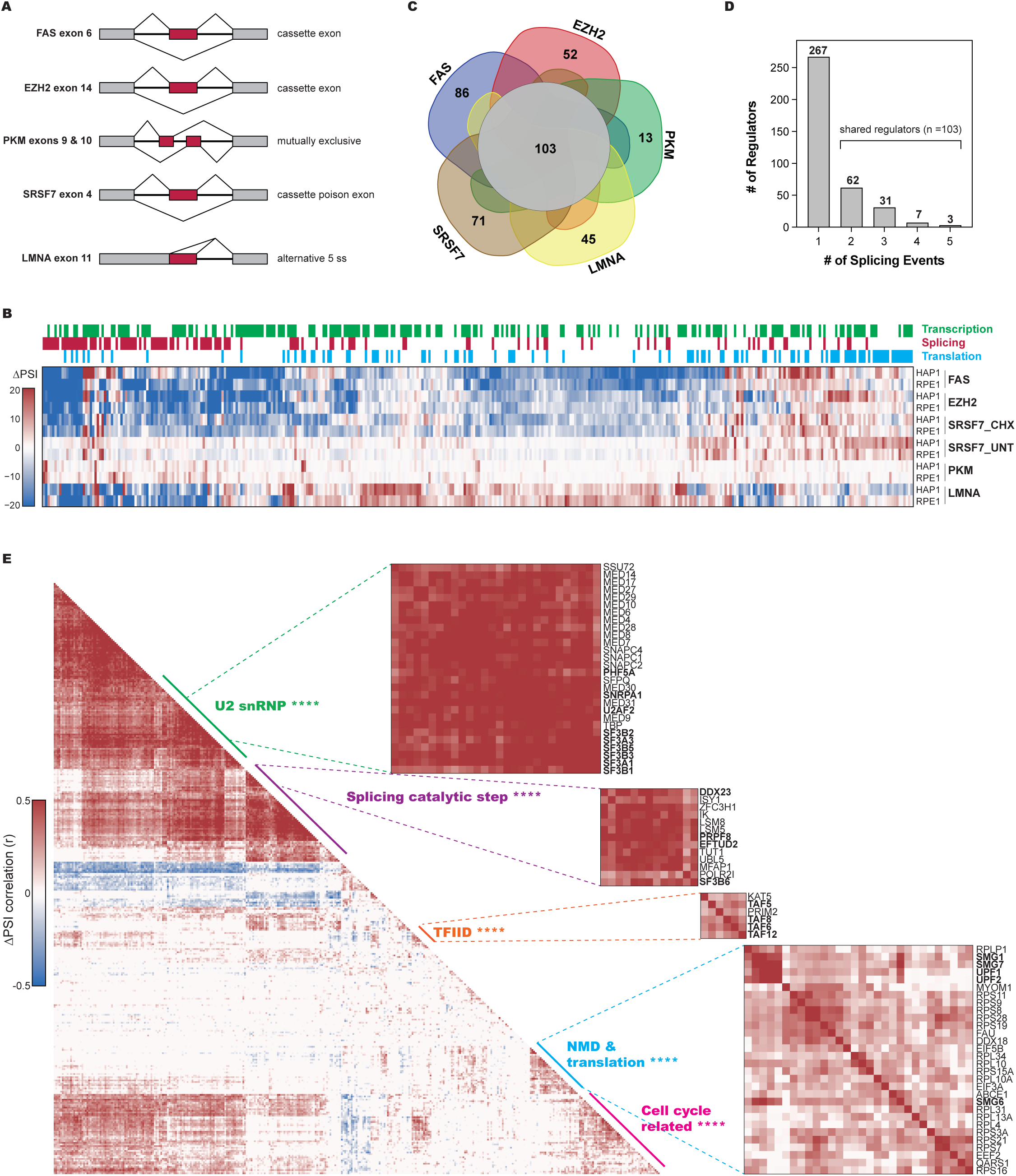
Genome-wide identification of splicing regulatory factors associated with disease-related events. **(A)** Schematic representation of the five alternative splicing events profiled using the CRASP-Seq platform in this study. **(B)** Heatmap of ΔPSI values for CRASP-Seq screen hits across the five alternative splicing events profiled in HAP1 and RPE1 cell lines. Functional annotations of genes are shown above the heatmap. **(C)** Flower plot depicting the number of splicing regulators identified for each alternative splicing event. Regulators shared by at least two events are displayed at the center of the plot. **(D)** Distribution of splicing regulatory hits across the five profiled events. Shared regulators, defined as hits observed in two or more events, are highlighted (n = 103). **(E)** Heatmap of pairwise correlations of ΔPSI values for the five splicing reporter events in HAP1, RPE1, and HepG2 cells. Sub-clusters are annotated with molecular signatures database (MSigDB) terms, with adjusted p-values for enrichment shown. ****p-value < 0.0001; Individual genes within sub-clusters are shown on the right.

All three reporters were successfully cloned into the CRASP-Seq vectors, maintaining a tight distribution of the hgRNA library (**Figure S3A**). Reporter libraries were then used to generate lentivirus, which was used to transduce HAP1, RPE1, or HepG2 cells expressing Cas9 and Cas12a nucleases. CRASP-Seq assays were performed and analyzed following the same experimental and analytical pipeline described above. For the SRSF7 poison cassette exon, the screen was conducted both in the absence or presence of cycloheximide treatment, to allow or inhibit NMD, respectively.

As expected, and consistent with the findings from the FAS screen (**Figures S1E-F**), all our screens effectively identified well-established regulators of the profiled alternative splicing events, as corroborated by previous studies. Specifically, SRSF3^40^, HNRNPA1 and PTBP1^28^ were confirmed as activators of PKM exon-10 (**Figure S3B**), while *SF3B3* knockout was identified to reduce EZH2 exon-14 inclusion (**Figure S3C**), consistent with previous studies^31^ and our focused assays (**Figure 1B**). Additionally, SRSF7 was confirmed as the strongest regulator^41,43^ of its own poison exon (**Figure S3D**). Notably, there is a strong enrichment of translation-related genes among those that promote SRSF7 poison exon inclusion, specifically in untreated cells (**Figures S3D-E**). This observation aligns with the well-established principle that disruptions in translation strongly impact NMD.^44–46^ RT-PCR assays further confirmed this regulation, as cells transduced with *SRSF7* minigene reporters showed increased exon inclusion following cycloheximide treatment, as expected given that the SRSF7 poison exon is an NMD target^41,42^ (**Figure S3F**). In contrast, expression of *SRSF7*-targeting hgRNAs reduced exon inclusion, consistent with the autoregulatory role of the SRSF7 poison exon (**Figure S3F**). Together, these results validate our perturbation-readout approach for identifying cellular factors and pathways modulating minigene splicing.

Interestingly, the profiled reporters identified several regulators, which to our knowledge have not been previously associated with the corresponding events. For instance, RBM39, a U2 snRNP auxiliary factor^47–49^, was found to activate PKM exon-10, while ILF3 and hnRNPK were found to repress PKM exon-10. These findings were further validated through RT-PCR assays that monitored the endogenous PKM mutually exclusive exons in HEK293T cells (**Figure S3G**). Since proliferative cells predominantly use PKM exon-9, with PSI values exceeding 95%, the PSI differences following ILF3 or hnRNPK depletion were minimal. However, when these genes were co-depleted with SRSF3—a factor that promotes exon-10 inclusion—a pronounced repressive effect of ILF3 and hnRNPK on exon-10 was observed (**Figure S3G**). In contrast, knockdown of the splicing regulator RBFOX2—which was not identified as a hit in the CRASP-Seq screen and was used as a negative control—does not result in substantial changes in splicing of the mutually exclusive PKM exons (**Figure S3G**). These data validate ILF2 and ILF3 as negative regulators of PKM2 (i.e., the exon-10 containing isoform) and align with recent studies showing that ILF2 and ILF3 selectively regulate the splicing of mutually exclusive exons.^50^ Moreover, they underscore the high sensitivity of the CRASP-Seq methodology.

Additionally, RBM39, XAB2, and SRSF7 were among several novel identified regulators of *EZH2* exon-14 (**Figure S3C**). The involvement of these genes was confirmed by monitoring endogenous EZH2 exon-14 in both HEK293T cells and patient-derived renal cell carcinoma lines, where aberrant activation of EZH2 exon-14 has been linked to poor prognosis^31^ (**Figures S3H**). Consistent with our screening data (**Figure S3C**), RBM39 exhibits a stronger effect in promoting EZH2 exon-14 inclusion compared to SF3B3 (**Figure S3H**), the only previously identified regulator of this event.^31^

Finally, we identified RBM39, SF3B1, SF3B3 and TNPO3, a nuclear import receptor for serine/arginine-rich (SR) proteins^51,52^, as potential regulators of SRSF7 poison exon (**Figure S3D**). Indeed, RT-PCR assays monitoring the endogenous SRSF7 poison exon confirmed that siRNA-mediated knockdown of these factors reduces poison exon inclusion in an NMD-independent manner (**Figure S3I**). Collectively, these validations underscore the efficacy of CRASP-Seq in uncovering novel regulatory factors involved in alternative splicing events.

Taken together, our five screens identified in total 370 factors that regulate alternative splicing, with significant overlap observed across the different cell lines (**Figures 2B and S4A-B; Table S3**). Principal component analysis suggests that the specific splicing event is the primary determinant of splicing changes across different genetic perturbations (**Figure S4C**). While a substantial number of regulators are shared among the various splicing events, each event also exhibits unique regulators (**Figures 2C-D**). A focused analysis comparing shared regulators with event-specific ones reveals a stronger enrichment of core spliceosome proteins within the shared regulator group (**Figure S4D**), consistent with studies showing that core spliceosome machinery can modulate alternative splicing events.^13–17^

Functional gene ontology analysis across the hits identified by all 5 reporters shows a strong enrichment in RNA splicing and processing terms, along with additional categories such as RNA metabolism and gene regulation, consistent with results from the *FAS* and *EZH2* screens alone (**Figures S4E-G**). The observed enrichment for DNA transcription-related genes is likely driven by factors involved in spliceosome biogenesis through their regulation of snRNA expression, such as components of the mediator complex. However, transcription factors may also influence splicing indirectly—by controlling the expression of key splicing regulators—or even directly by binding to RNA, as previously demonstrated.^53–57^ Consequently, this diverse gene set likely encompasses examples of all these mechanisms. The domains enriched among the factors identified by our screens include several RNA-binding domains such as the RNA recognition motif (RRM) and the C3H zinc finger domain, indicating an enrichment of screen hits to regulate splicing by directly associating with RNA **(Figure S4H)**.

To explore the functional interplay among the factors identified as regulators of alternative splicing by our screens, we assessed the overall correlations in PSI values across the five examined splicing events, conducting pairwise knockout comparisons of all identified splicing regulators (**Figure 2E**). Our screens reveal correlated perturbation responses among factors associated with well-characterized complexes. For example, we identified a cluster of components associated with U2 snRNP, including core U2 snRNP subunits (such as SF3B and SF3A subcomplexes, *SNRPA1*, and *PHF5A*) and auxiliary factors like *U2AF2*. Interestingly, the U2 snRNP cluster also included the mediator complex (**Figure 2E**), aligning with previous findings that indicate a requirement of mediator for U2 snRNA expression.^58,59^ Furthermore, in line with recent studies connecting SF3B6 to PRPF8 and later spliceosomal complexes^17^, we observed a closer association of *SF3B6* with U5 snRNP components (*PRPF8*, *EFTUD*, and *DDX23*) rather than with other U2 snRNP complexes (**Figure 2E**). Finally, we observed correlated perturbations among several core components of the NMD machinery (*UPF1*, *UPF2*, *SMG1*, *SMG7*, and *SMG6*) alongside ribosomal proteins and other translation-related factors (**Figure 2E**). Collectively, these observations underscore the effectiveness of our screening strategy and its potential to provide insights into the physical and functional interactions among trans-acting factors.

### ZNF207 regulates LMNA aberrant splicing in progeria

A *de novo*, synonymous, point mutation (1824C>T) in the *LMNA* gene activates a cryptic 5′ splice site, leading to Hutchinson-Gilford Progeria Syndrome (HGPS), a rare genetic disorder marked by premature aging^20,21^. To identify regulators of LMNA aberrant splicing, we performed a CRASP-Seq assay using a minigene reporter containing this mutation. This approach uncovered multiple genes that regulate LMNA splicing, with several capable of restoring splicing to wild-type levels despite the mutation (**Figure 3A; Table S3**). As expected, the impact of these genes is contingent on the presence of the mutation, and when the screen is performed with a wild-type reporter, only the canonical LMNA isoform is generated (**Figure S5A**). These genes are significantly enriched for RNA splicing and gene regulation-related functions (**Figure S5B**). Among them, we identified ZNF207, also known as BuGZ, which was initially characterized as a mitotic spindle assembly protein due to its stable association with BUB3 via a GLEBS motif^60,61^ (**Figure 3B**). ZNF207 is a zinc finger (ZnF) protein, which has been shown to also function as a transcription factor.^62,63^ Additionally, ZNF207 has been found to bind RNA directly, co-purify with splicing-related proteins and influence RNA metabolism, although its direct role in splicing regulation remains unclear.^63–69^ Given ZNF207’s unclear role in splicing regulation yet its high score in our CRASP-Seq screen (**Figure 3A**), we chose to investigate its impact on LMNA aberrant splicing and the broader regulation of alternative pre-mRNA splicing.

**Figure 3:**
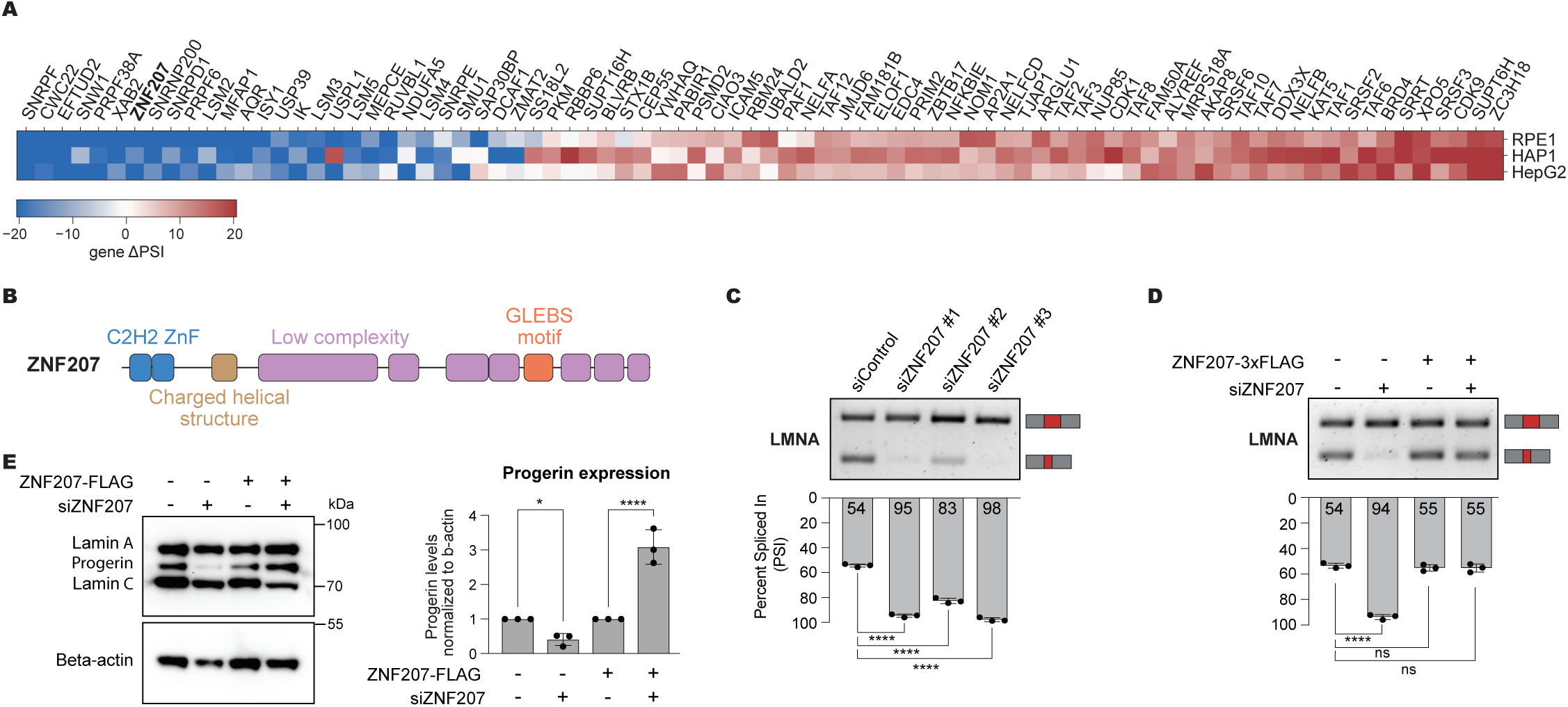
CRASP-Seq identifies ZNF207 as a regulator of progerin aberrant splicing. **(A)** Heatmap showing ΔPSI values of mutant LMNA CRASP-Seq screen hits in HAP1, RPE1, and HepG2 cells. **(B)** Schematic representation of the ZNF207 protein structure, highlighting the C2H2 zinc finger (ZnF) domain, the charged helical structure, low-complexity regions, and the Gle2-binding-sequence (GLEBS) motif. **(C)** RT-PCR analysis of aberrant LMNA splicing in RNA from HGPS patient-derived immortalized fibroblasts treated with three independent siRNAs targeting ZNF207. Quantifications of PSI values from three independent experiments are shown below the gel. Data are presented as mean ± standard deviation (SD). ****p-value < 0.0001; Dunnett’s multiple comparisons test following one-way ANOVA. **(D)** RT-PCR analysis of LMNA splicing in HGPS patient fibroblasts treated with ZNF207-targeting siRNA and/or ectopically expressing a ZNF207 ORF resistant to siRNA for nine days. Quantification of PSI values from three independent experiments are shown below the gel. Data are presented as mean ± SD. ****p-value < 0.0001; Dunnett’s multiple comparisons test following one-way ANOVA. **(E)** Western blot analysis of lamins in HGPS patient-derived immortalized fibroblasts transduced with either a siRNA-resistant 3×FLAG C-terminal-tagged ZNF207 ORF or an empty vector. Cells were treated with either non-targeting control siRNA or siRNA targeting endogenous ZNF207 (siZNF207) for 16 days. Blots were probed with antibodies against lamins, ZNF207, and GAPDH (loading control). Relative progerin expression was quantified and normalized to β-actin, with values further normalized to the corresponding siControl-treated sample. Quantifications from three independent experiments are presented on the right. Data are presented as mean ± SD. *p-value < 0.05, ****p-value < 0.0001; Uncorrected Fisher’s LSD following one-way ANOVA.

To confirm ZNF207’s role in modulating LMNA aberrant splicing, we employed an orthogonal RNAi approach. Knocking down ZNF207 with three independent siRNAs in HEK293T cells expressing the LMNA splicing reporter, results in significant correction of LMNA splicing with all three siRNAs tested (**Figures S5C-D**). To rule out off-target effects, we cloned and expressed a ZNF207 open reading frame (ORF) resistant to the most effective siRNA targeting ZNF207 (**Figure S5E**). Restoring ZNF207 expression partially rescues the aberrant LMNA splicing of the minigene reporter (**Figure S5F**).

Next, we used immortalized fibroblasts from HGPS patients to assess the role of ZNF207 in promoting progerin isoform expression. All three siRNAs tested effectively restore endogenous LMNA splicing (**Figures 3C**), an effect that is reversed by ectopic stable expression of the ZNF207 ORF (**Figures 3D and S5G**). Importantly, knockdown of ZNF207 in HGPS patient-derived fibroblasts leads to a significant reduction in progerin protein levels (**Figure 3E**), despite its well-documented stability.^70,71^ Notably, this effect, akin to the changes in LMNA splicing levels, is reversed upon reintroducing an siRNA-resistant ZNF207 ORF into the cells (**Figure 3E**). These findings confirm that ZNF207 depletion enhances canonical LMNA splicing, validating the results of our screen.

To determine whether depleting ZNF207 could rescue molecular phenotypes associated with progeria, we assessed its impact on hallmark disease features in immortalized fibroblasts. Progerin expression disrupts nuclear envelope morphology and impacts key epigenetic markers, such as H3K27me3 and H4K16ac, alongside reduced levels of lamina-associated polypeptide 2α (LAP2α) and Lamin B1 (LB1).^71–74^ The overall phenotype observed in the HGPS patient-derived fibroblasts was relatively mild (**Figure S5H**). Long-term (i.e., 15 days) knockdown of ZNF207 in immortalized fibroblasts from healthy individuals results in a marked decrease of LAP2α, H3K27me3, and H4K16ac which is typically observed in cells undergoing senescence. Consistent with its essential nature, ZNF207 depletion in HGPS-derived fibroblasts also lowered LAP2α and H4K16ac levels and did not restore H3K27me3 or LB1 levels (**Figure S5H**). Taken together, our data indicate that ZNF207 depletion effectively corrects *LMNA* aberrant splicing. However, sustained depletion of ZNF207 impairs cell fitness, consistent with its role as an essential gene.^75^

### ZNF207 depletion results in widespread alternative splicing changes

To investigate the broader impact of ZNF207 on alternative splicing regulation, we depleted ZNF207 in HEK293T cells using three independent siRNAs. Our alternative splicing analysis identified thousands of affected splicing events, with the majority being skipped alternative cassette exons. However, we also observed alternative 5′ and 3′ splice sites, as well as some instances of intron retention (**Figure 4A; Table S4**). We observed a significant overlap in the regulated splicing events across the three siRNA treatments, and the PSI correlation between them shows a strong positive relationship (**Figures S6A-B**).

**Figure 4:**
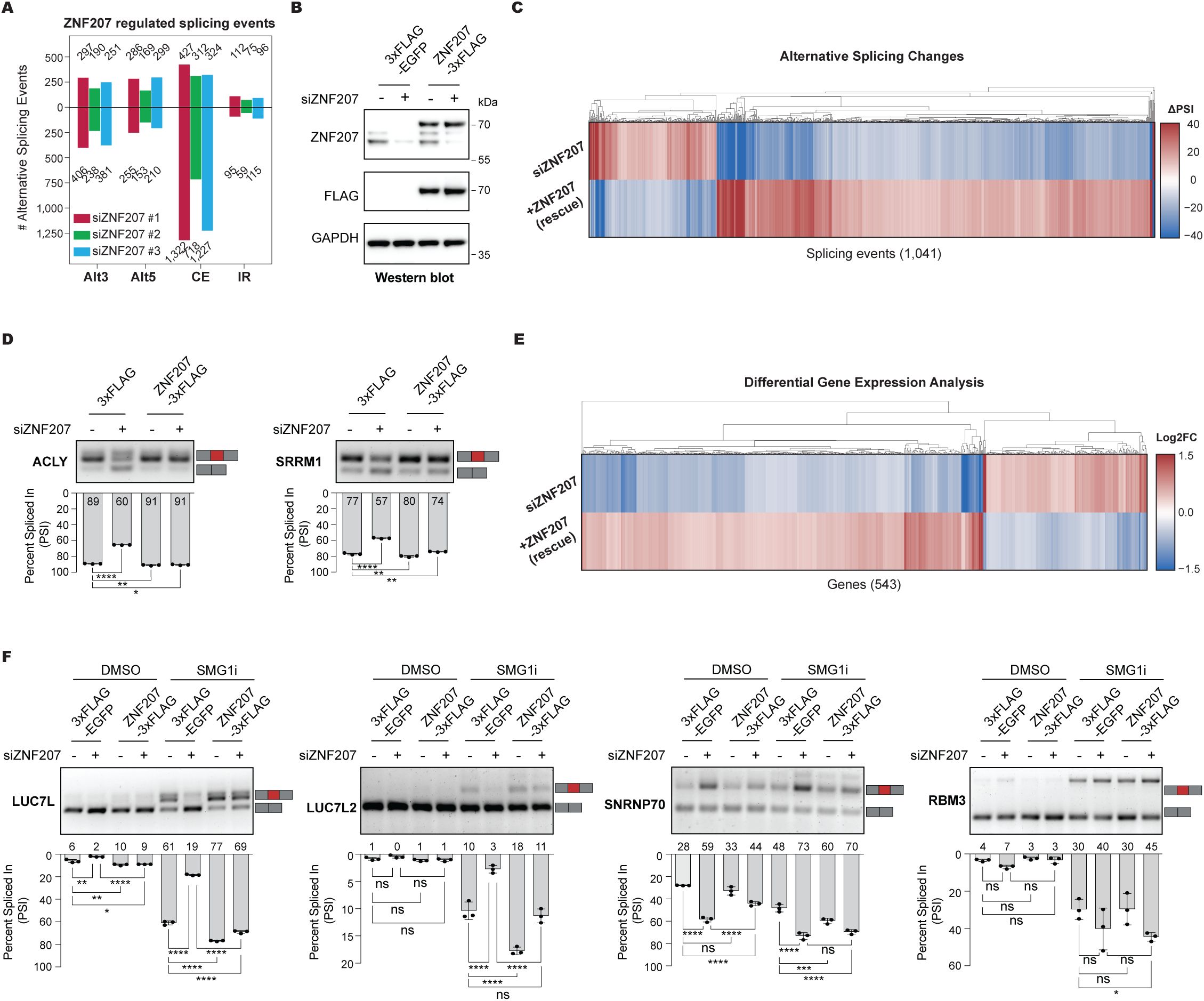
ZNF207 depletion induces widespread alternative splicing changes. **(A)** Distribution of alternative splicing event types (CE: cassette exons, Alt3: alternative 3′ splice sites, Alt5: alternative 5′ splice sites, IR: retained introns) detected by RNA-seq following *ZNF207* knockdown for 48 hours with three independent siRNAs in HEK293T cells. Events with significant splicing changes (|ΔPSI| ≥ 10; probability ≥ 0.95) upon siZNF207 knockdown are included. **(B)** Western blot analysis of ZNF207 in HEK293 Flp-In cells expressing doxycycline-inducible, siRNA-resistant 3×FLAG C-terminal tagged ZNF207 ORF. Cells were treated with control siRNA or siRNA targeting endogenous ZNF207 (siZNF207) for 48 hours. Blots were probed with antibodies against ZNF207, FLAG, and GAPDH (loading control). **(C)** RNA-seq analysis of splicing changes (ΔPSI) upon *ZNF207* knockdown in HEK293 Flp-In cells. Comparisons include knockdown of *ZNF207* relative to non-targeting siRNA treatment without induction (siZNF207) and ZNF207 ORF rescue expression in siRNA-treated cells compared to expression of empty vector (siZNF207 + rescue). Events with significant splicing changes (|ΔPSI| ≥ 10; probability ≥ 0.95) upon siZNF207 knockdown and rescue by ZNF207 ORF expression are highlighted. **(D)** RT-PCR analysis of ACLY (left) and SRRM1 (right) alternative splicing in HEK293 Flp-In cells treated with ZNF207-targeting siRNA and/or expressing siRNA-resistant C-terminally 3xFLAG-tagged ZNF207 ORF. Quantification of PSI values from three independent experiments are shown below the gel. Data are presented as mean ± SD. *p < 0.05, **p < 0.01, ****p < 0.0001; Dunnett’s multiple comparisons test following one-way ANOVA. **(E)** Gene expression analysis (Z score normalized) from RNA-seq profiling of HEK293 Flp-In cells treated with ZNF207-targeting siRNA and/or rescued with siRNA-resistant ZNF207 ORF expression. Only genes with significant expression changes (adjusted p < 0.05; |log2FC| ≥ 0.35) upon siZNF207 and rescued by ZNF207 expression are shown. **(F)** RT-PCR analysis of LUC7L, LUC7L2, SNRNP70, and RBM3 alternative splicing in HEK293 Flp-In cells treated with ZNF207-targeting siRNA and/or expressing siRNA-resistant ZNF207 ORF. Experiments were conducted with and without the NMD inhibitor SMG1i (0.5 µM, 6 hrs). Quantification of PSI values from three independent experiments are shown below the gel. Data are presented as mean ± SD. *p < 0.05, **p < 0.01, ***p < 0.001, ****p < 0.0001; Šídák’s multiple comparisons test following one-way ANOVA.

To rule out the possibility that these splicing changes are due to siRNA off-target effects, we complemented this approach by employing the Flp-In system to reintroduce an siRNA-resistant ZNF207 ORF with a C-terminal FLAG tag. ZNF207-FLAG expression was maintained at levels comparable to its endogenous counterpart (**Figure 4B**). As anticipated, the majority of alternative splicing changes are successfully rescued identifying more than 1,000 splicing events controlled by ZNF207 (**Figure 4C; Table S5**). Additionally, we validated the RNA sequencing data through focused RT-PCR assays on a few selected events. These assays confirmed our splicing analyses, and we were able to replicate the alternative splicing patterns and rescue them by reintroducing the ZNF207-FLAG ORF (**Figures 4D**). Notably, attempting to rescue using an N-terminal FLAG-tagged ZNF207 ORF failed, despite successful expression of the ZNF207 transgene, suggesting that the N-terminal FLAG tag may interfere with the splicing activity of the protein (**Figures S6C-D**). Collectively, these data confirm that ZNF207 plays a significant role in influencing alternative splicing choices.

Previous studies suggest that ZNF207 functions as a transcription factor, and that it impacts alternative splicing indirectly by regulating the expression of splicing factors.^62,63^ To investigate this, we analyzed our ZNF207-depleted and rescued transcriptomic data to identify genes whose expression is regulated by ZNF207. This analysis revealed 543 protein-coding regulated genes (**Figure 4E; Table S6**), with a modest enrichment for those involved in amino acid metabolism (**Figure S6E**). Only a few of those genes are linked to pre-mRNA splicing, including *LUC7L* and *LUC7L2*, which show increased expression following ZNF207 depletion, and *SNRNP70* and *RBM3*, which exhibit reduced expression. These findings were confirmed by qRT-PCR (**Figure S6F**). However, all these genes also display ZNF207-regulated splicing events predicted to affect transcript stability via NMD (**Figure S6G**). We validated these splicing events through RT-PCR (**Figure 4F**) and demonstrated that most of them indeed influence transcript stability via NMD (**Figures 4F and S6F**). These findings suggest that ZNF207 does not influence splicing by controlling the transcription of splicing-related genes, but it rather affects transcript stability, at least in part, through alternative splicing coupled to NMD.

### ZNF207 binds regulated exons and associates with the splicing machinery

To explore the mechanisms by which ZNF207 influences alternative splicing, we employed proximity biotin labeling using TurboID followed by streptavidin capture and mass spectrometry to identify ZNF207-associated proteins (**Figure 5A**). We generated HEK293 Flp-In cell lines expressing miniTurbo-tagged ZNF207 at near endogenous levels (**Figure S7A**). Biotin labelling experiments were performed in triplicate, and the detected mass spectrometry peptides were scored for significant enrichment. Only proteins detected and enriched in all three replicates were considered as ZNF207 proximal interactors (see Methods). In total we identified 187 proteins to be in close proximity with ZNF207 (**Figure 5B; Table S7**). In line with ZNF207’s known role in kinetochore function through its stably association with BUB3^60,61^, the TurboID analysis identified BUB3 and a few additional kinetochore-related proteins. We also detected several transcription and chromatin-related factors, consistent with ZNF207’s previously reported function as a transcription factor (**Figure 5B**).^62^

**Figure 5:**
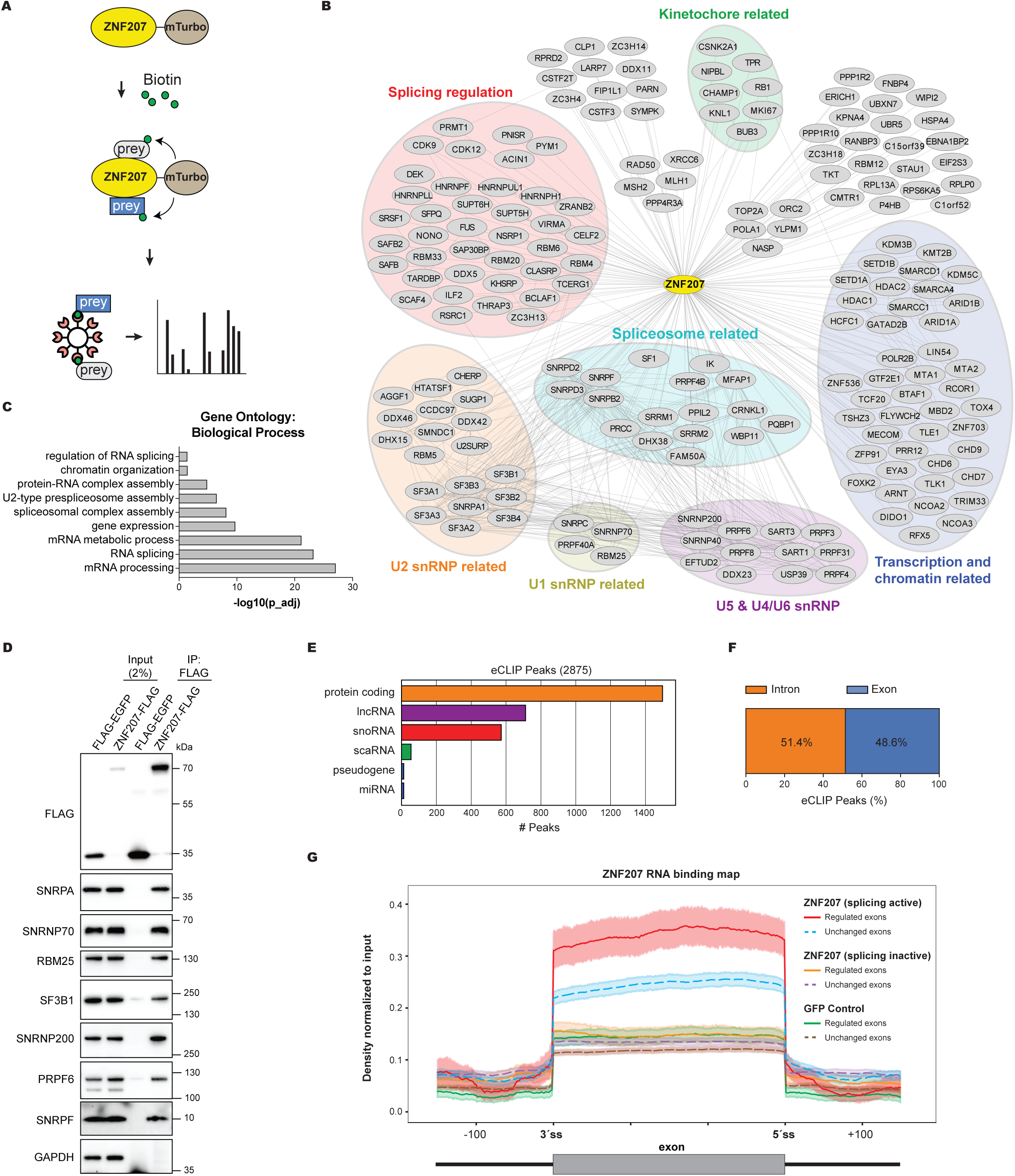
ZNF207 associates with the splicing machinery and binds to regulated splicing targets. **(A)** Overview of the TurboID experimental pipeline. HEK293 Flp-In cells expressing C-terminal miniTurbo-tagged ZNF207 were induced for 24 hours and incubated with biotin for 2 hours. Proximal biotinylated proteins were purified and analyzed via mass spectrometry. **(B)** Proximal interaction network of ZNF207 identified by TurboID. Only significant interactors consistently detected across all three replicates are shown (Table S7). **(C)** Gene Ontology (GO) enrichment analysis of biological processes among ZNF207 TurboID proximal interactors with FDR < 0.05. **(D)** Western blot analysis of total cell lysates (input) treated with benzonase, and FLAG immunoprecipitates (IP: FLAG-M2) from HEK293 Flp-In cells expressing 3×FLAG-ZNF207. Blots were probed with antibodies against FLAG, SNRPA, SNRNP70, RBM25, SF3B1, SNRNP200, PRPF6, SNRPF, and GAPDH (negative control). **(E)** Bar plot showing the distribution of ZNF207 eCLIP peaks across RNA biotypes. Only biotypes with at least 10 ZNF207 peaks are included. eCLIP reads were normalized to input, and peaks were called using the DEWSeq pipeline (see Methods for details). **(F)** Stacked bar chart showing the percentage of intronic and exonic ZNF207 binding peaks. **(G)** Average eCLIP signal profiles of C-terminal FLAG-tagged ZNF207 for ZNF207-regulated cassette exons (n = 715) and unchanged alternative exons (n = 3,745). Profiles include the negative control GFP and N-terminal FLAG-tagged ZNF207 (devoid of splicing activity) for the same exon subsets.

Notably, the TurboID data reveal that ZNF207-associated proteins are very highly enriched in factors involved in pre-mRNA splicing, including U1 snRNP, U2 snRNP, U4/U6.U5 tri-snRNP, and numerous alternative splicing auxiliary factors, extending previous observations (**Figure 5B**).^66,67^ Gene ontology analysis further supports this, showing strong enrichment for terms related to RNA processing and splicing, including complexes such as the spliceosome, 17S U2 snRNP, and C splicing complexes (**Figures 5C and S7B**).

To validate these findings, we performed co-immunoprecipitation experiments using HEK293 Flp-In cells expressing FLAG-tagged ZNF207 (**Figure 4B**). These experiments confirmed that ZNF207 co-purifies with several endogenous splicing factors tested, including U1-related SNRPA, SNRNP70, RBM25; U2-related SF3B1; U5-related SNRNP200, and PRPF6 as well as SNRPF, in an RNA-independent manner (**Figure 5D**). Collectively, our results reveal that ZNF207 associates with the splicing machinery, suggesting a direct role in the regulation of alternative splicing.

To further investigate the direct role of ZNF207 in alternative splicing regulation, we performed enhanced crosslinking and immunoprecipitation followed by sequencing (eCLIP-Seq). We used our HEK293 Flp-In cells stably expressing FLAG-tagged ZNF207 for RNA immunoprecipitation. As controls, we included cells expressing GFP-FLAG as well as N-terminal FLAG-tagged ZNF207, which is splicing-incompetent (**Figures S6C-D**).

The eCLIP-Seq analysis reveals that ZNF207 preferentially binds to protein coding genes (**Figure 5E**). Among the detected peaks we observed an approximately equal distribution between exonic and intronic binding (**Figure 5F**). Strikingly, the binding maps for ZNF207 show a strong binding enrichment over the coding sequences of regulated exons, as compared to exons unaffected by ZNF207 depletion (**Figures 5G and S7C**). This binding pattern is unique to the splicing-competent C-terminal FLAG-tagged ZNF207 and absent in the splicing-incompetent N-terminal FLAG-tagged variant or the GFP-FLAG negative controls. These results suggest that ZNF207 binding to regulated exons may be important for its role in splicing regulation.

### CRASP-Seq coupled to high-throughput mutagenesis identifies key regions of ZNF207 for alternative splicing regulation

To gain deeper mechanistic insights into ZNF207’s role in splicing regulation, we conducted a high-throughput tiling mutagenesis screen to map critical functional regions, leveraging recent advances in genome editing.^76–78^ For this screen, we replaced the CHyMErA loss-of-function hgRNA cassette in our CRASP-Seq vectors with a tiling Cas9 base editor library (**Figure 6A**). In addition to *ZNF207*, this library also targets representative genes from U1, U2, and U5 snRNP complexes, identified through TurboID-based ZNF207 interaction profiling, along with top hits from our original LMNA CRASP-Seq screen, yielding a total of 39 target genes. The final library comprises 8,594 Cas9 gRNAs, including 1,520 intergenic and 90 non-targeting controls (**Table S8**).

**Figure 6:**
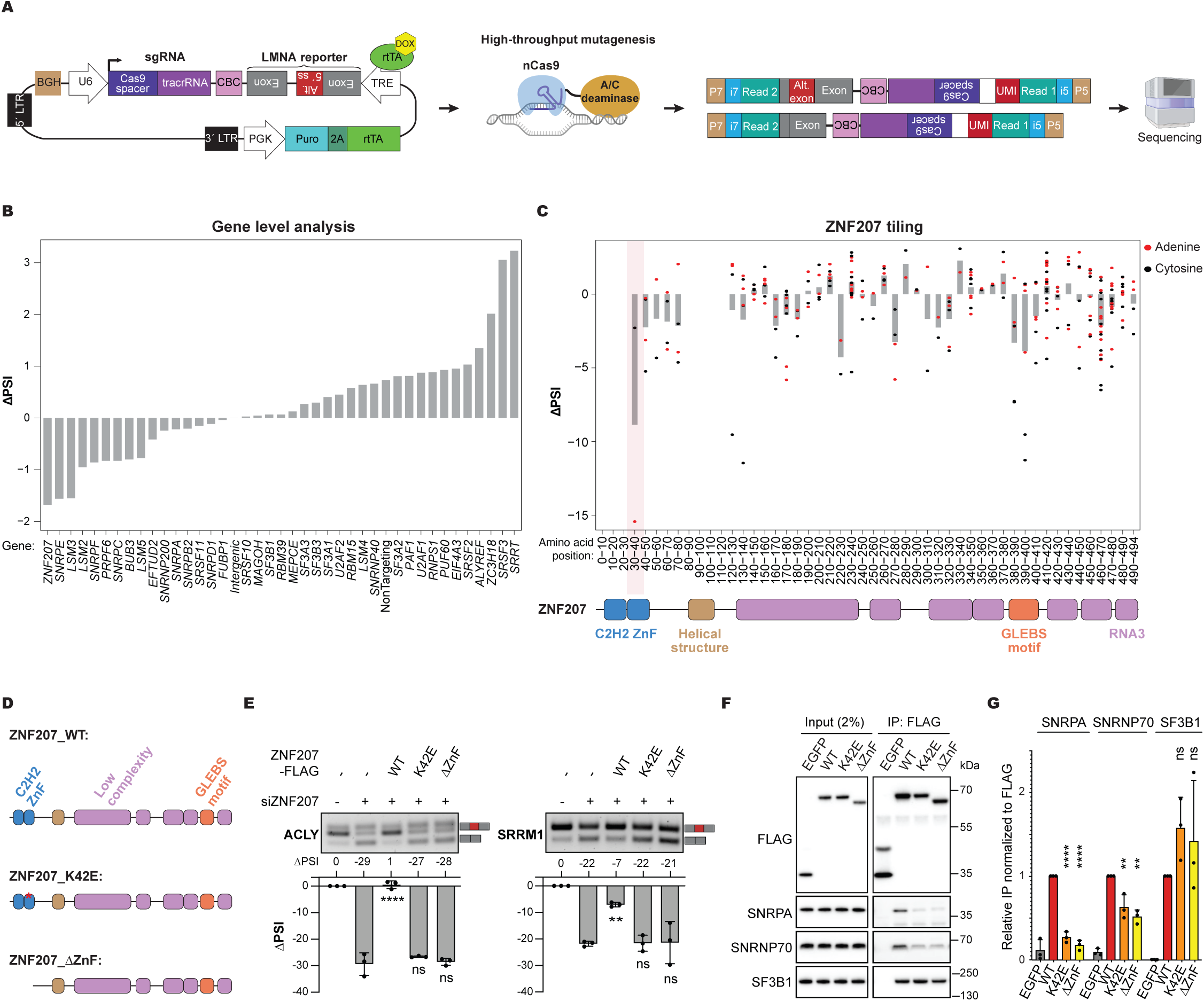
High-Throughput mutagenesis reveals a critical residue required for ZNF207-dependent splicing regulation. **(A)** Schematic of the CRASP-Seq base editor tiling approach. A lentiviral vector enables constitutive expression of sgRNAs while a doxycycline-inducible TRE promoter drives the expression of a mutant *LMNA* minigene reporter. Splicing-derived mRNA products are analyzed via Illumina paired-end high-throughput sequencing. **(B)** ΔPSI scores from gene-level analysis of all guides in the base editor library, including those targeting splice sites, across the 39 profiled genes. Intergenic and non-targeting controls are also included. The genes are arranged in ascending order of their ΔPSI scores, from left to right. **(C)** ΔPSI scores for base editor sgRNAs targeting ZNF207 coding sequence, excluding splice-site-overlapping guides. Average ΔPSI scores for individual sgRNAs within 10-amino-acid windows are plotted, with the scores of individual sgRNAs depicted as black dots. The ZNF207 protein is illustrated below, showing ZnF domains (blue), GLEBS motif (orange), and low-complexity regions (purple). **(D)** Schematic representation of ZNF207 constructs: full-length wild-type ZNF207 (top), full-length K42E point mutant (middle), and the ΔZnF truncation mutant lacking the zinc finger domain (bottom). **(E)** RT-PCR analysis of ACLY (left) and SRRM1 (right) alternative splicing in HEK293 Flp-In cells treated with ZNF207-targeting siRNA and/or expressing siRNA-resistant wild-type or truncation mutant ZNF207 ORFs. Quantification of ΔPSI values from three independent experiments are shown below the gel. Data are presented as mean ± SD. Statistical comparisons were performed relative to the siZNF207 sample without ZNF207 ORF expression. ****p < 0.0001; Dunnett’s multiple comparisons test following one-way ANOVA. **(F)** Western blot analysis of total cell lysates (input) treated with benzonase, and FLAG immunoprecipitates (IP: FLAG-M2) from HEK293 Flp-In cells expressing 3×FLAG-tagged ZNF207 variants. Blots were probed with antibodies specific for FLAG, SNRPA, SNRNP70, RBM25, SF3B1, and GAPDH (negative control). **(G)** Quantification of co-immunoprecipitated spliceosome components, normalized to FLAG-ZNF207 pulldown efficiency and wild-type (WT) ZNF207 levels. Data are presented as mean ± SD. Statistical comparisons were performed relative to WT-ZNF207. ***p < 0.001, **p < 0.01, *p < 0.05; One-way ANOVA with Dunnett’s multiple comparisons correction.

Following library cloning and LMNA incorporation into the CRASP-Seq vector, we conducted screens in HAP1 cells expressing highly efficient adenine (ABE8e) and cytosine (evoCDA1) base editors, as we established previously.^11^ Gene-level analysis revealed that guides targeting *ZNF207* lead to the greatest reduction in progerin isoform expression (**Figure 6B; Table S8**). In contrast, mutagenesis of *SRRT, SRSF3* and *ZC3H18* produce the largest progerin isoform increases, consistent with findings from the CHyMErA loss-of-function screen (**Figure 3A**). These concordant results underscore the robustness of our screening platforms, despite employing different gRNA libraries and approaches.

A detailed analysis of *ZNF207*-targeting gRNAs identified regions overlapping the ZnF domains as having the most pronounced impact on LMNA splicing when mutated (**Figure 6C; Table S8**). ZNF207 is predicted to be predominantly a disordered protein, containing multiple low-complexity regions^79^ (**Figures S8A-B**). Intriguingly, HydRA analysis, a tool for predicting RNA-binding proteins and associated domains^80^, identifies portions of these disordered regions, including the C-terminal region, as potential RNA-binding sites (**Figure S8B**).

To dissect the regions of ZNF207 responsible for RNA binding and interactions with splicing factors, we generated eight distinct ZNF207 truncation mutants targeting critical regions, including the ZnF domains, GLEBS motif, and C-terminal region (**Figure S8C**). We initially evaluated the ability of these ZNF207 mutants to rescue splicing alterations caused by the knockdown of endogenous *ZNF207*. Transient transfection of ZNF207 truncation mutants into HEK293T cells revealed that only the ZnF domains are critical for alternative splicing activity (**Figures S8D–E**). This finding aligns with our CRASP-Seq base editor mutagenesis screen, which identified gRNAs targeting the ZnF domains as having the strongest impact on LMNA minigene reporter splicing (**Figure 6C**). To address concerns that the observed rescue of splicing phenotypes might result from high expression levels achieved via transient transfection, we generated stable HEK293 Flp-In cell lines expressing these mutants at near-endogenous levels (**Figure S8F**). RT-PCR rescue assays corroborated our initial findings, confirming that the ZnF domains are the sole essential region of the ZNF207 protein required for splicing catalysis (**Figure S8G**).

Our CRASP-Seq high-throughput mutagenesis screen identified a gRNA overlapping the second ZnF domain of ZNF207 that had the strongest impact on ZNF207’s ability to promote aberrant LMNA splicing. This guide is predicted to mutate the region encoding amino acids H41, K42, and K43. To validate the functional importance of this region, we generated a panel of eleven ZNF207 mutants, each containing single, dual or triple predicted amino acid substitutions within this span. These constructs were expressed in cells depleted of endogenous ZNF207 to assess their ability to rescue splicing activity. Remarkably, constructs bearing even a single amino acid substitution (i.e., K42E) completely lost splicing regulatory activity, as measured by the modulation of endogenous ZNF207 splicing targets (**Figure S8H-I**). To confirm these findings, we generated stable cell lines expressing either wild-type or K42E-mutant ZNF207 (**Figure 6D**) at near-endogenous levels and evaluated their ability to rescue splicing events upon depletion of endogenous ZNF207. Consistently, only the WT, and not the K42E mutant, restored normal splicing of ACLY and SRRM1 transcripts (**Figure 6E**). Notably, K42 lies within the β-sheet of the second ZnF domain, and its substitution with glutamic acid is not predicted to alter the overall folding of the domain. These results underscore the power of the CRASP-Seq platform to pinpoint individual amino acid residues that are critical for the functional activity of splicing regulators.

Taken together, these data support that the ZnF domains are essential for ZNF207’s ability to regulate splicing. To test whether the ZnF domains are also sufficient, we generated truncation mutants containing either the ZnF domains alone or the ZnF domains followed by the adjacent helical region (ZnF+Hel; **Figures S9A**). While the ZnF domains alone were unable to rescue the splicing changes induced by siRNA-mediated knockdown of *ZNF207*, the ZnF+Hel truncation mutant fully restored splicing following transient transfection in a K42-dependent manner (**Figures S9B**). However, when expressed at near-endogenous levels using Flp-In cells (**Figures S9C**), the ZnF+Hel mutant achieved only a partial rescue of splicing changes (**Figures S9D**). These results suggest that the C-terminal disordered region of ZNF207 it is also important for its full splicing regulatory activity.

To further investigate the mechanism by which the ZNF207 regions influence alternative splicing, we leveraged the splicing-inactive mutants we have previously generated. Using stable Flp-In cell lines, we performed in vivo RNA crosslinking assays and co-immunoprecipitation experiments to assess the effects of these mutations on ZNF207’s interactions with RNA and spliceosomal proteins, respectively. RNA crosslinking experiments revealed no appreciable difference in RNA binding between ZNF207 truncation mutants lacking splicing regulatory activity and the wild-type protein (**Figure S9E**). In contrast, the ZnF+Hel mutant, which retains splicing competence, exhibits only weak RNA binding (**Figure S9F**). These findings align with the HydRA analysis and indicate that ZNF207 associates with RNA through multiple, potentially redundant segments within its C-terminal disordered region. Moreover, they indicate that RNA binding alone is not sufficient to confer splicing regulatory activity. Consistently, the K42E point mutant, despite being functionally inactive in splicing regulation, also crosslinked to RNA at levels comparable to wild-type (**Figure S9F**).

We next examined how the ZNF207 mutations affect its interactions with spliceosomal proteins. Notably, the ZnF domains are required for interactions with a specific subset of splicing factors, particularly components of the U1 snRNP complex, such as SNRPA and SNRNP70 (**Figure S9G-H**). In contrast, truncation mutants lacking the ZnF domains retained the ability to bind the U2 snRNP component SF3B1. Strikingly, the K42E mutant showed significantly reduced interaction with both SNRPA and SNRNP70, while its association with SF3B1 remained largely unaffected (**Figure 6F-G**). Together with our RT-PCR analyses (**Figure 6E**), these findings suggest that ZNF207 regulates alternative splicing through specific protein-protein interactions with components of the U1 snRNP, and that disruption of these interactions impairs its splicing regulatory function.

In summary, we have developed CRASP-Seq, a robust method for systematically identifying splicing regulatory factors for specific events. Applying CRASP-Seq to study the *LMNA* 1824C>T mutation revealed ZNF207 as a key splicing regulator. In HGPS patient cells, depletion of ZNF207 corrects LMNA splicing which, as expected, leads to significantly reduced levels of progerin protein (**Figure 3E**). ZNF207 binds to regulated exons and participates in critical interactions with U1 snRNP components to broadly control alternative splicing decisions (**Figure 7**). These findings showcase the utility of CRASP-Seq in uncovering critical regulatory mechanisms in splicing.

**Figure 7:**
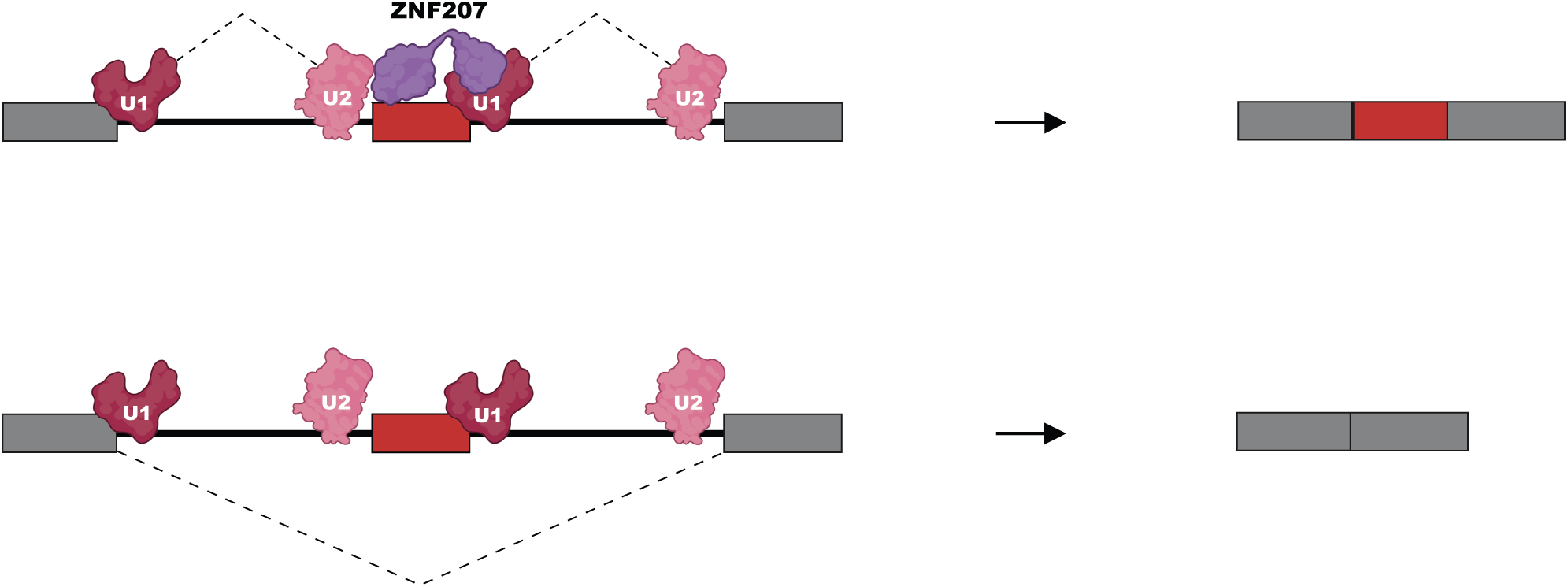
Model of ZNF207 as a regulator of alternative splicing. Illustration of the proposed mechanism by which ZNF207 regulates alternative splicing. ZNF207 interacts with U1 snRNP and other core spliceosome components, binding directly to target pre-mRNAs. These interactions influence splice site selection, promoting the activation of specific exons.

## DISCUSSION

In this study, we describe a CRISPR-based strategy for the effective identification of splicing regulatory factors and domains controlling phenotypically relevant alternative splicing events. Our method, CRASP-Seq, links CRISPR-induced genetic perturbations with quantifiable splicing phenotypes using RNA high-throughput sequencing as a readout. This approach allows precise measurement of the impact of each protein-coding gene in the human genome on the splicing outcome of a reporter, all from a single RNA extraction. We have demonstrated that CRASP-Seq is a versatile and powerful platform capable of inducing diverse perturbations—including gene knockouts and high-throughput mutagenesis—and of capturing a wide range of alternative splicing events, such as cassette exons, mutually exclusive exons, and alternative 5′ splice sites. These examples of CRASP-Seq to examine cancer-related and progeria-associated splicing events highlights the utility of our screening platform in identifying novel splicing regulators with significant implications for health and disease.

Several advantages distinguish our methodology from existing approaches. CRISPR technology, when compared to RNAi screens, offers increased sensitivity and fewer off-target effects.^81^ CRASP-Seq directly reads out RNA phenotypes using sequencing, similar to recent methods applied in yeast to measure gene expression.^82,83^ Therefore, in contrast to most CRISPR-based screening approaches applied in mammalian cells that require specialized equipment to isolate cells with a desired phenotype via microscopy or cell sorting (e.g., flow cytometry)^84–86^, CRASP-Seq only necessitates access to a high-throughput sequencer. This approach not only streamlines the screening workflow but also enhances the accessibility and affordability of such experiments while providing more precise splicing information. Unlike previously described CRISPR screening methods, which rely on detecting gRNA abundance changes in fluorescence-sorted populations^84–86^, CRASP-Seq enables the direct analysis of splice junctions in regulated splicing events. This provides highly quantitative data, allowing accurate determination of splicing changes, such as ΔPSI values. By utilizing a single RNA extraction from a pooled cell population transduced with the CRASP-Seq lentiviral vector, this approach is both cost-efficient and practical, avoiding the need for extensive robotics or thousands of RNA extractions typically required for arrayed screens.^15,53^ Importantly, CRASP-Seq provides a powerful platform for high-resolution mutational scanning of splicing modulators, enabling the identification of critical protein regions and individual amino acid residues involved in splicing regulation. CRASP-Seq is straightforward to implement, and the entire process—from cloning the minigene reporter into a CRISPR perturbation vector to obtaining screening results—can be completed within 1-2 weeks. Consequently, CRASP-Seq represents a highly accessible and economical tool for dissecting the regulatory factors and protein regions governing functionally significant splicing events.

Using our CRASP-Seq methodology, we identified genes that influence the aberrant splicing event in *LMNA* that cause the rare pediatric premature aging disorder HGPS. Among the significant hits, we identified several splicing-related genes, including *SRSF6*, a known negative regulator of progerin isoform expression.^87^ In contrast, ZNF207 and several core spliceosome proteins emerged as positive regulators of the progerin isoform. ZNF207 is a zinc finger protein previously shown to participate in kinetochore formation^60,61^ and function as a transcription factor.^62^ Our results identify ZNF207 as a regulator of alternative splicing. This is further supported by recent temporal mRNA interactome data, which reveal a binding profile similar to that of pre-mRNA splicing factors.^69^

Our findings suggest that ZNF207 functions as a bona fide regulator of alternative pre-mRNA splicing, with roles that extend beyond the aberrant splicing event observed in *LMNA*. Transcriptomic analyses identified over 1,000 splicing events influenced by ZNF207, with these effects reversed upon reintroduction of the ZNF207 ORF, reinforcing the validity of the observations. Additionally, several genes exhibiting differential expression following ZNF207 depletion display associated splicing changes predicted to induce NMD, suggesting that these splicing events contribute to the observed gene expression alterations. We propose that “splicing regulator” should join and perhaps surpass “kinetochore-related protein” and “transcription factor” as functional descriptors of ZNF207.

Our data suggest that ZNF207 regulates alternative splicing by mediating interactions with U1 snRNP components, as mutants unable to interact with U1 snRNP fail to rescue splicing alterations. These findings align with recent systematic studies aiming to map the functional network of splicing-related factors, suggesting ZNF207 as functionally associated with U1-related components, such as SNRNP70 and RBM25, particularly in the context of cassette exon inclusion.^17^ Interestingly, our study uncovers a feedback compensatory mechanism in which ZNF207 protein levels regulate the expression of auxiliary U1 splicing regulators, such as LUC7L and LUC7L2, by modulating poison exon inclusion in these genes.^88–90^ This finding further underscores the intricate link between ZNF207 and the U1 snRNP complex.

ZNF207 is an intrinsically disordered protein prone to phase separation and coacervation.^79^ Multivalent assemblies of ZNF207 may promote exon inclusion by stabilizing cross-exon interactions with U2 snRNP and RNA, following its initial recruitment through binding to U1 snRNP via its ZnF domains. Our mutational analyses indicate that ZNF207’s RNA-binding capacity is not localized to a single defined region but instead arises from various disordered regions that bind RNA in a redundant manner. This is consistent with our eCLIP sequencing data, which reveal preferential association of ZNF207 with the coding sequences of regulated splicing events. These findings suggest that ZNF207 associates with RNA in a largely non-sequence-specific manner, likely as a secondary consequence of its critical protein–protein interactions with the spliceosomal machinery, particularly U1 snRNP. Future structural studies of spliceosomal assemblies containing ZNF207 could provide deeper insights into these multifaceted interactions and their impact on splicing regulation.

Notably, BUB3, a stable interactor of ZNF207 and well-characterized kinetochore protein^60,61^, exerts only a modest effect on LMNA splicing and other ZNF207-regulated splicing events (**Figure S10A-C**). This observation aligns with the finding that the GLEBS domain, responsible for BUB3 interaction, is dispensable for ZNF207’s splicing activity (**Figure S8C-G**). Together, these data support the notion that ZNF207’s roles in kinetochore assembly and splicing regulation are functionally distinct. This functional uncoupling has potential therapeutic implications. Specifically, small molecules that selectively disrupt the interaction between ZNF207 and the U1 snRNP—without interfering with its kinetochore function—could modulate splicing in a targeted manner without affecting cell division. When combined with recent advancements in antisense oligonucleotides^71^ and base-editing technologies^91^, which specifically address the HGPS mutation, these approaches could contribute to a more effective therapeutic toolkit.

ZNF207 joins an expanding list of C2H2 zinc finger proteins with RNA splicing activity.^53,57,92^ Our high-throughput mutagenesis CRASP-Seq experiment revealed that the ZnF domains are essential for ZNF207 splicing activity, a conclusion further supported by truncation mutant assays. While ZnFs are predominantly recognized for mediating nucleic acid interactions, several examples of ZnFs facilitating protein-protein interactions have been documented.^93^ Consistent with this, the ZnF domains of ZNF207 enable its association U1 snRNP factors, further highlighting the versatility of ZnFs in molecular interactions. We anticipate that applying CRASP-Seq to additional alternative splicing events will uncover more previously overlooked splicing regulators.

---

**Supplementary Figure 1:**
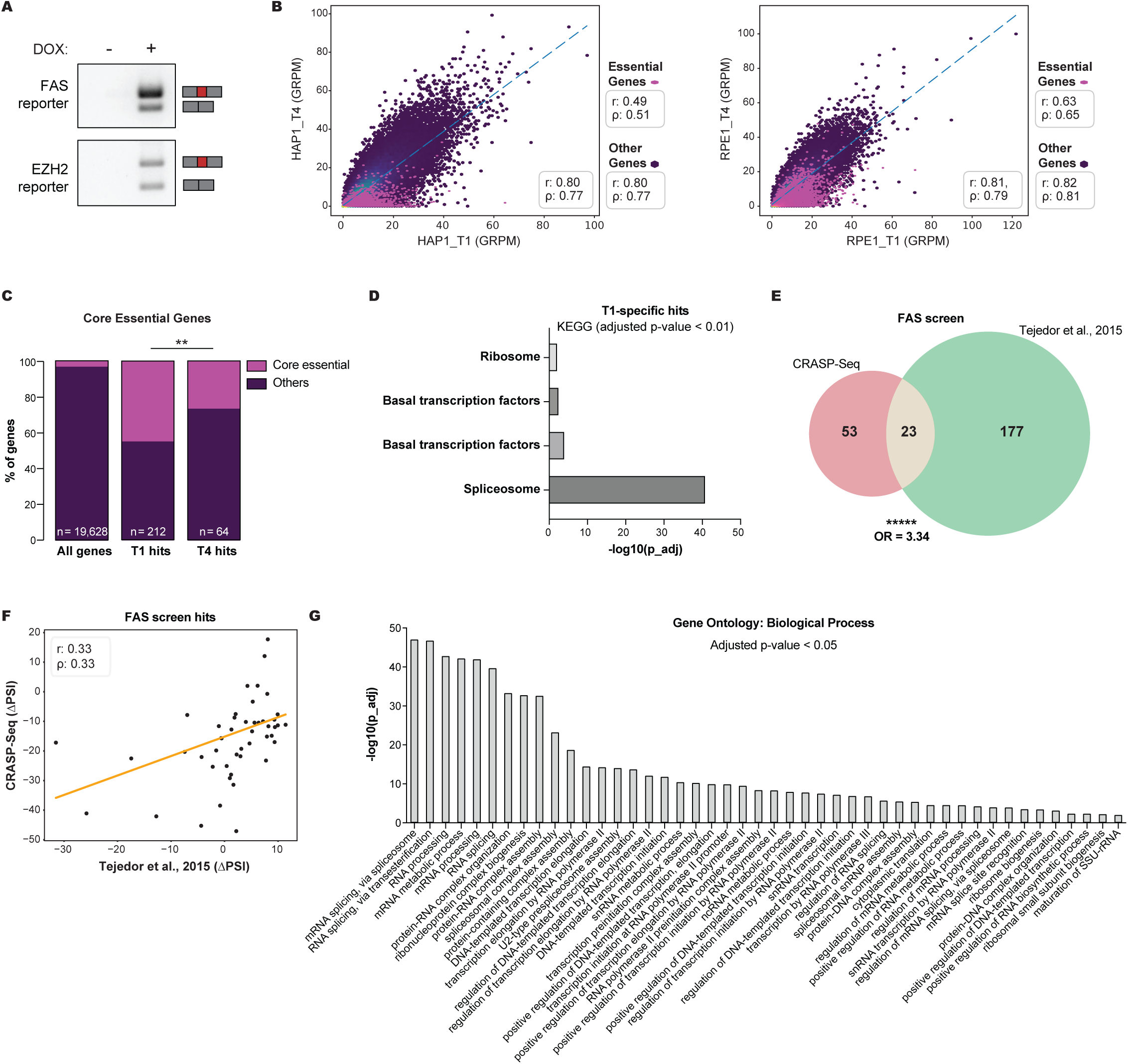
CRASP-Seq effectively identifies splicing regulatory factors. Related to Figure 1. (A) RT-PCR analysis of FAS and EZH2 minigene reporters expressed in HAP1 cells transduced with CRASP-Seq vectors. Cells were treated with 1 µg/mL doxycycline (+) or not (–) for 20 hours before RNA extraction. (B) Correlation analysis of guide read counts per million (GRPM) at early (T1) and late (T4) time points in the FAS CRASP-Seq screen. Scatter plots and Pearson correlation coefficients are shown for HAP1 (left) and RPE1 (right) cell lines, with separate analysis of common essential genes versus other genes. “r” denotes the Pearson correlation coefficient and “ρ” represents the Spearman’s rank correlation coefficient. (C) Bar plots showing the percentage of core-essential genes94 among all library-targeted genes, T1 CRASP-Seq hits, and T4 CRASP-Seq hits. **p-value = 0.0092; odds ratio = 2.25; Fisher’s exact test. (D) Enrichment of Kyoto Encyclopedia of Genes and Genomes (KEGG) biological pathways among T1-specific (i.e., lost in the T4 point) screen hits. Pathways with adjusted p-value < 0.01 are displayed. (E) Venn diagram comparing FAS screen hits identified by CRASP-Seq with those from a genome-wide siRNA robotics screen.15 ****p-value < 0.00001; OR = 3.34; Fisher’s exact test. (F) Correlation plot of differential splicing (ΔPSI) values for gene hits identified by CRASP-Seq. ΔPSI values are shown for CRASP-Seq versus the siRNA robotics screen.15 “r” denotes the Pearson correlation coefficient and “ρ” represents the Spearman’s rank correlation coefficient. (G) Gene Ontology (GO) analysis of biological processes enriched among regulators of FAS and EZH2 splicing events identified via the CRASP-Seq pipeline (related to Figure 1D). Hits with adjusted p-value < 0.05 are displayed.

**Supplementary Figure 2:**
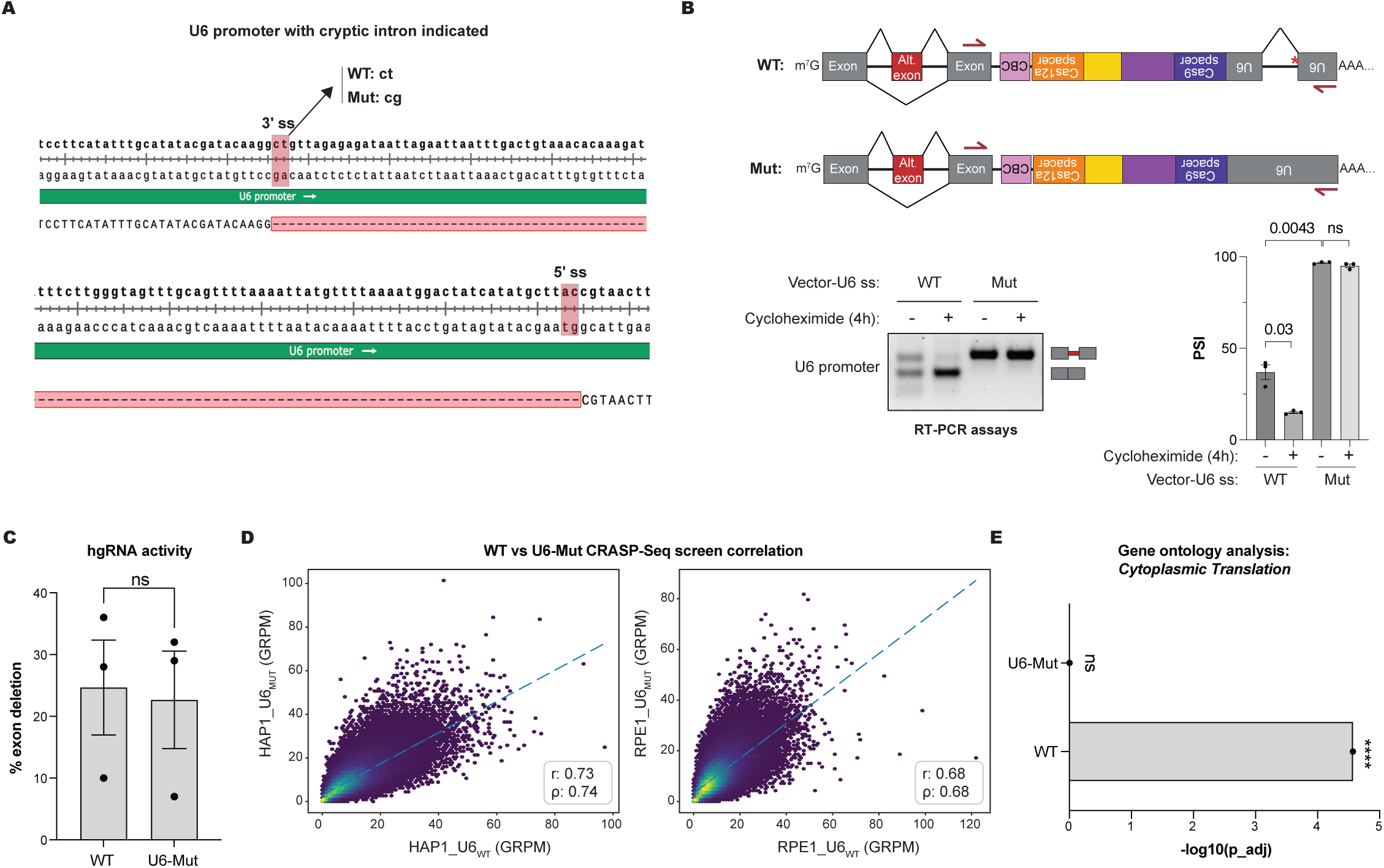
A point mutation in the CRASP-Seq vector eliminates a cryptic 3′ splice site and prevents spurious NMD. Related to Figure 1. **(A)** Sequence of the U6 promoter region showing the cryptic 3’ and 5’ splice sites and the associated intron. The position of the introduced point mutation designed to inactivate the cryptic splice site is highlighted. WT: wild-type; Mut: U6 point mutation. **(B)** RT-PCR analysis (bottom left) and PSI quantification (bottom right) for RNA extracted from HAP1 cells transduced with wild-type (WT) or U6 point-mutated (Mut) CRASP-Seq vectors. Cells were treated with 100 µg/mL cycloheximide or DMSO for 4 hours prior to RNA extraction. A schematic illustrating the primer annealing sites is shown at the top. Data from independent replicate experiments are shown. Statistical significance was assessed using a two-tailed Welch’s t-test. **(C)** Editing efficiency of wild-type (WT) and U6 point-mutated (Mut) CRASP-Seq vectors was evaluated using HPRT1 exon deletion as a readout.33 Three independent HPRT1 exon deletion hgRNAs were transduced, and deletion efficiency was measured from PCR-amplified genomic DNA. Statistical significance was assessed using a two-way paired t-test. **(D)** Correlation of guide read counts per million (GRPM) between CRASP-Seq screens conducted with wild-type (WT) and U6 point-mutated (U6-Mut) vectors. Scatter plots and Pearson correlation coefficients are shown for HAP1 (left) and RPE1 (right) cell lines. “r” denotes the Pearson correlation coefficient and “ρ” represents the Spearman’s rank correlation coefficient. **(E)** Bar plot showing the –log10 transformed adjusted p-values for enrichment of cytoplasmic translation-related terms among CRASP-Seq screen hits identified using wild-type (WT) versus U6 point-mutated (U6-Mut) lentiviral libraries.

**Supplementary Figure 3:**
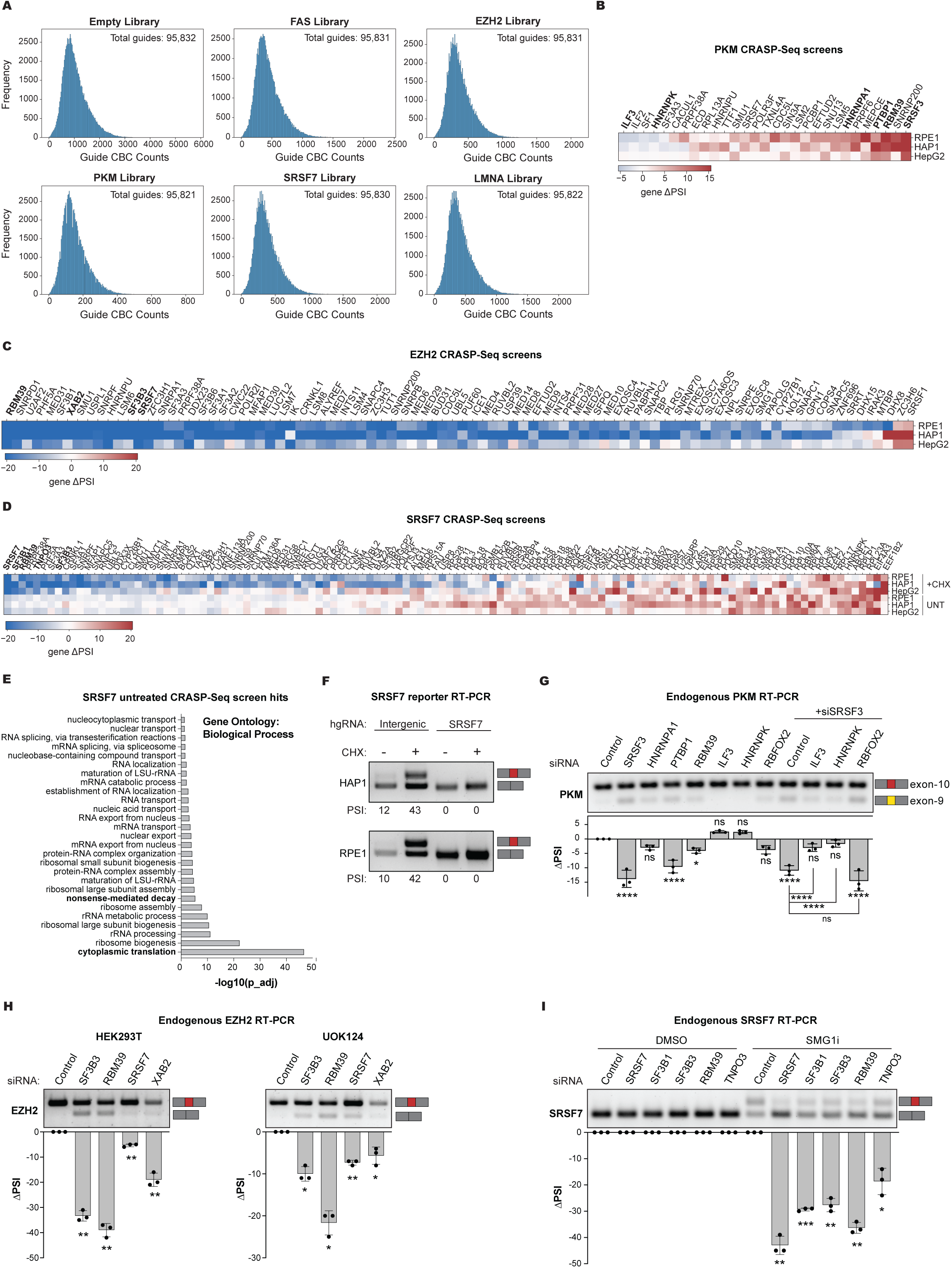
Genome-wide CRASP-Seq screens identify regulators of disease-related splicing events. Related to Figure 2. **(A)** Histogram showing the distribution of read counts for the 95,893 hgRNAs in the CRASP-Seq library. Separate histograms are displayed for the empty library as well as all five reporter libraries. **(B)** Heatmap of genes identified by CRASP-Seq screens as regulators of *PKM* alternative splicing. Differential splicing levels (ΔPSI) relative to intergenic controls are shown. Genes promoting exon-10 inclusion (PKM2 isoform) with positive ΔPSI values are highlighted in red, while genes promoting exon-9 inclusion (PKM1 isoform) are highlighted in blue. Genes selected for RT-PCR validation are highlighted in bold. **(C)** Heatmap of genes identified as regulators of EZH2 exon-14 alternative splicing in CRASP-Seq screens. Differential splicing levels (Δ PSI) relative to intergenic controls are shown, with genes promoting exon-14 inclusion highlighted in blue. Genes selected for RT-PCR validation are highlighted in bold. **(D)** Heatmap of genes identified as regulators of SRSF7 poison exon splicing in CRASP-Seq screens. Differential splicing levels (ΔPSI) relative to intergenic controls are shown both with and without cycloheximide treatment. Genes promoting poison exon inclusion are highlighted in blue. Genes selected for RT-PCR validation are highlighted in bold. **(E)** Gene Ontology (GO) analysis of biological processes enriched among regulators of the *SRSF7* poison exon identified via the CRASP-Seq pipeline in untreated cells. Hits with adjusted p-value < 0.05 are displayed. **(F)** RT-PCR validation of alternative splicing for SRSF7 poison exon using the CRASP-Seq *SRSF7* minigene reporter in HAP1 (top) and RPE1 (bottom) cells co-expressing intergenic or SRSF7-targeting hgRNAs. Cells were treated with 100 µg/mL cycloheximide or DMSO for 4 hours prior to RNA extraction. **(G)** RT-PCR analysis of endogenous PKM alternative splicing in HEK293 cells. The indicated genes (or combinations) were knocked down using siRNAs for 50 hours before RNA extraction. Quantification of ΔPSI values from three independent experiments are displayed below the gel. Data are represented as mean ± standard deviation (SD). *p-value < 0.05, ****p-value < 0.0001; two-way ANOVA. **(H)** RT-PCR analysis of endogenous EZH2 alternative splicing in HEK293 (left) and UOK124 renal cell carcinoma (right) cell lines. The indicated genes were knocked down using siRNA for 48 hours before RNA extraction. Quantification of ΔPSI values from three independent experiments are displayed below the gel. Data are presented as mean ± SD. *p-value < 0.05, **p-value < 0.01; Welch’s t-test with Holm-Šídák method multiple correction testing. **(I)** RT-PCR analysis of endogenous SRSF7 poison exon splicing in HEK293 cells. Cells were treated with 0.5 µM SMG1i or DMSO for 8 hours prior to RNA extraction. Quantifications of ΔPSI values from three independent experiments are displayed below the gel. Data are presented as mean ± SD. *p-value < 0.05, **p-value < 0.01, ***p-value < 0.001; Welch’s t-test with Holm-Šídák method multiple correction testing.

**Supplementary Figure 4:**
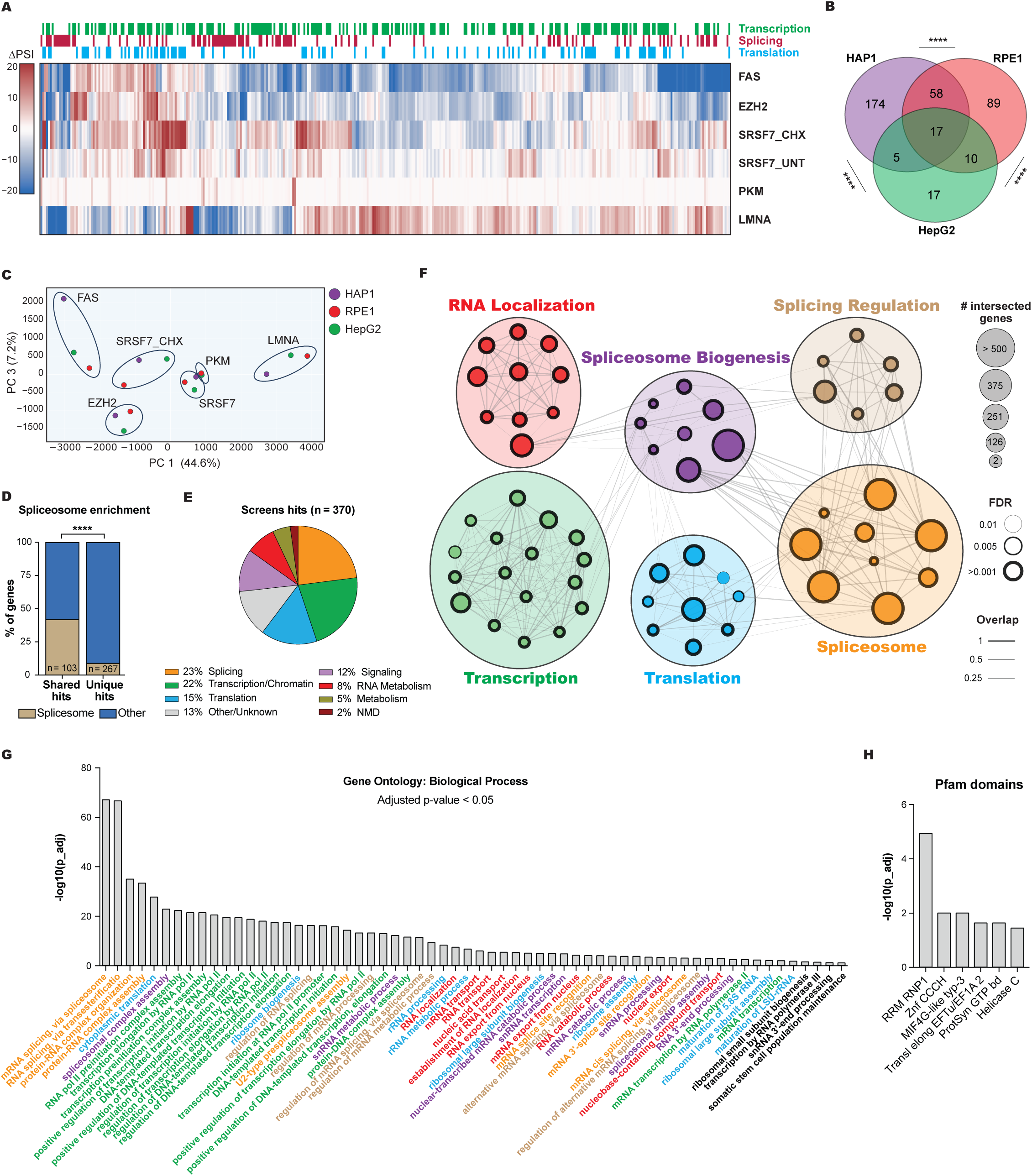
Genome-wide identification and functional characterization of alternative splicing regulators. Related to Figure 2. **(A)** Heatmap of ΔPSI values for CRASP-Seq screen hits across the five alternative splicing events in HepG2 cells. Functional annotations of genes are shown above the heatmap. **(B)** Venn diagram showing the overlap of CRASP-Seq screen hits identified across all five splicing reporters in three different cell lines. ****p-value < 0.0001; Fisher’s exact test. **(C)** Principal component analysis (PCA) of the six CRASP-Seq screens across the three different cell lines. **(D)** Bar plots depicting the percentage of core spliceosome genes (as defined by containing CORUM spliceosome terms) among shared hits identified across two or more reporters compared to unique hits identified by only one reporter. p-value < 0.0001; odds ratio = 7.32; Fisher’s exact test. **(E)** Pie chart depicting the functional classification of genes identified as hits in the genome-wide CRASP-Seq screens. Categories were curated through literature review and functional enrichment analysis using the g:Profiler Gene Ontology (GO) database. **(F)** GO enrichment analysis of regulators identified by all CRASP-Seq regulatory screens. Only significantly enriched biological processes with adjusted p-value < 0.01 are displayed. **(G)** Detailed GO enrichment analysis of biological processes associated with regulators of all reporters identified via the CRASP-Seq pipeline. Hits with adjusted p-value < 0.05 are shown. GO terms are color-coded to align with the functional categories shown in Supplementary Figure 4F. **(H)** Pfam domain enrichment analysis of regulators identified by all CRASP-Seq regulatory screens using Enrichr.95 Only significantly enriched domains with adjusted p-value < 0.05 are displayed.

**Supplementary Figure 5:**
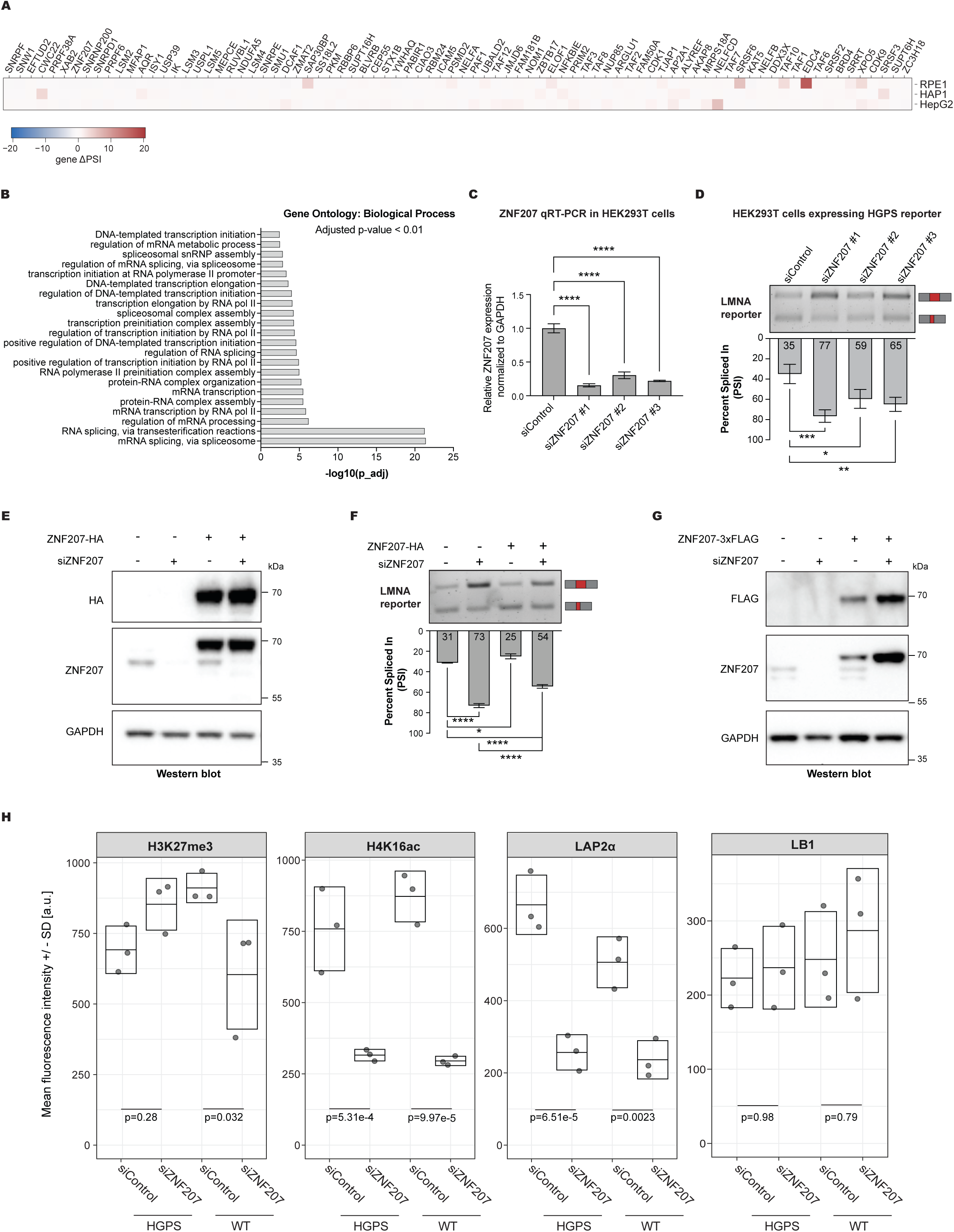
ZNF207 is a positive regulator of progerin aberrant splicing. Related to Figure 3. **(A)** Heatmap showing ΔPSI values for wild-type *LMNA* CRASP-Seq screens for the hits identified in the mutant *LMNA* screens across HAP1, RPE1, and HepG2 cell lines. **(B)** Gene Ontology (GO) enrichment analysis of biological processes among *LMNA* CRASP-Seq screen hits with adjusted p-value < 0.01. **(C)** Real-time quantitative RT-PCR measuring ZNF207 transcript depletion in HEK293 cells transfected with three independent siRNAs. Quantification of transcript levels from three independent experiments are displayed. Data are presented as mean ± SD. ****p-value < 0.0001; Dunnett’s multiple comparisons test following one-way ANOVA. **(D)** RT-PCR analysis of aberrant *LMNA* splicing in RNA from HEK293T cells transduced with the mutant *LMNA* minigene reporter and treated with three independent siRNAs targeting ZNF207. Quantification of PSI values from three independent experiments are shown below the gel. Data are presented as mean ± SD. *p-value < 0.05, **p-value < 0.01, ***p-value < 0.001; Dunnett’s multiple comparisons test following one-way ANOVA. **(E)** Western blot analysis of ZNF207 in HEK293T cells expressing the *LMNA* minigene reporter, transiently transfected with either a siRNA-resistant 3×HA C-terminal-tagged ZNF207 ORF or an empty vector. Cells were treated with either non-targeting siRNA control or siRNA targeting endogenous ZNF207 (siZNF207). Blots were probed with antibodies against ZNF207, HA tag, and GAPDH (loading control). **(F)** RT-PCR analysis of *LMNA* splicing in HEK293 cells expressing the *LMNA* minigene reporter and treated with ZNF207-targeting siRNA and/or ectopically expressing an siRNA-resistant ZNF207 ORF. Quantification of PSI values from three independent experiments are shown below the gel. Data are presented as mean ± SD. **p-value < 0.01, ****p-value < 0.0001; Dunnett’s multiple comparisons test following one-way ANOVA. **(G)** Western blot analysis of ZNF207 in HGPS patient-derived immortalized fibroblasts stably transduced with either a siRNA-resistant 3×FLAG C-terminal-tagged ZNF207 ORF or an empty vector. Cells were treated with either non-targeting siRNA control or siRNA targeting endogenous ZNF207 (siZNF207). Blots were probed with antibodies against ZNF207, FLAG, and GAPDH (loading control). **(H)** Single cell high throughput immunofluorescence quantification of LAP2α, H3K27me3, H4K16ac and LB1 expression levels in immortalized human fibroblasts treated with siZNF207 or siCTL for 15 days. At least 600 cells were analyzed in each experiment and represented as the distribution of average mean fluorescence intensity. The values from three independent experiments ± SD are plotted (see Methods). Statistical differences were analyzed by one-way ANOVA followed by the Dunnett’s test using the WT siCTL cell line as the negative control for WT siZNF207 and the HGPS siCTL as the negative control for HGPS siZNF207.

**Supplementary Figure 6:**
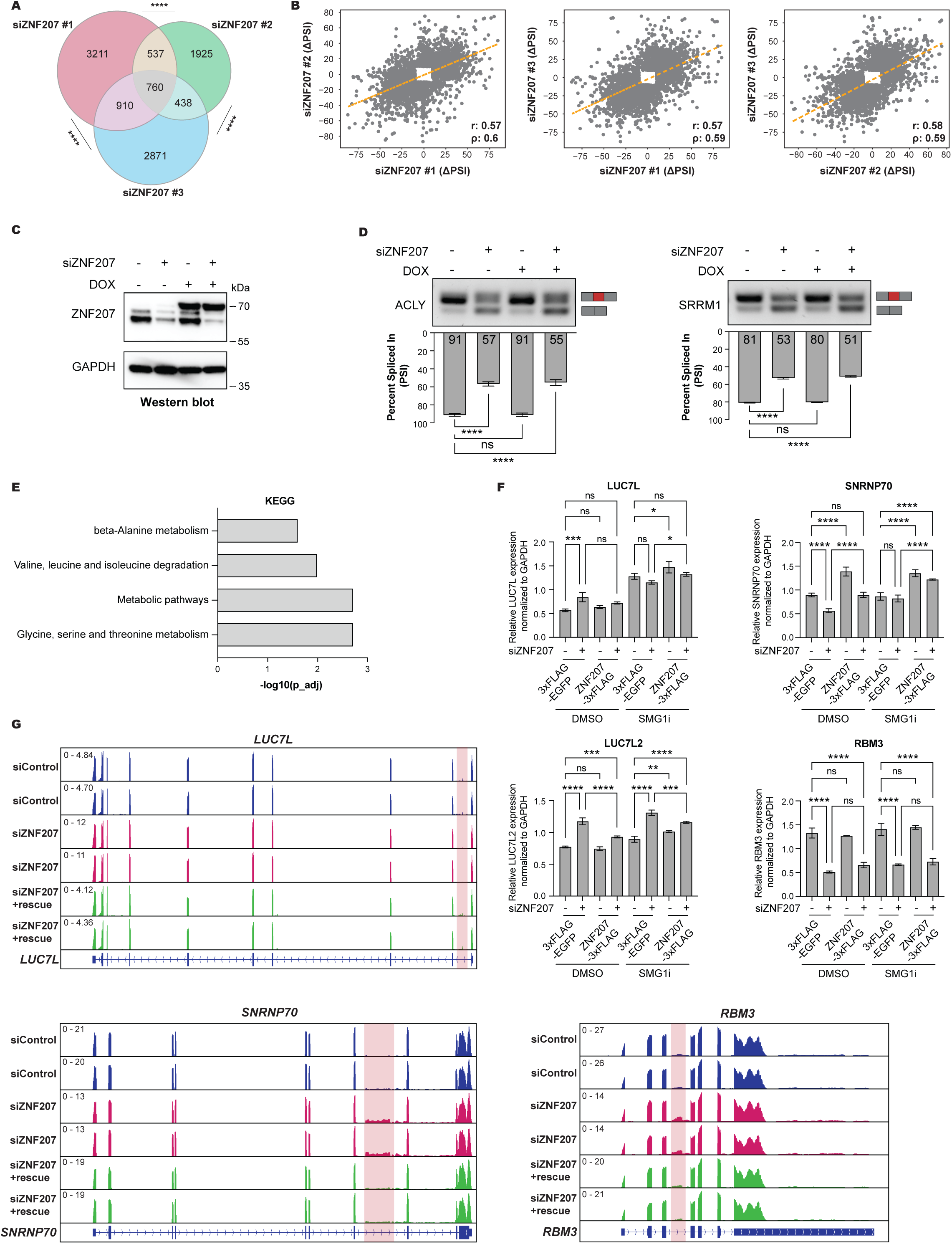
ZNF207 is an alternative splicing regulator. Related to Figure 4. **(A)** Venn diagram illustrating the overlap of regulated splicing events (|ΔPSI| ≥ 10 and probability ≥ 0.95) across the three independent siRNA sequences targeting ZNF207. ****p-value < 0.0001; Fisher’s exact test. **(B)** Correlation of PSI changes (ΔPSI) in HEK293T cells transfected with three independent siRNA sequences targeting ZNF207. Significant events for each siRNA treatment are highlighted using distinct colors. “r” denotes the Pearson correlation coefficient and “ρ” represents the Spearman’s rank correlation coefficient. **(C)** Western blot analysis of ZNF207 in HEK293 Flp-In cells expressing doxycycline-inducible, siRNA-resistant 3×FLAG N-terminal tagged ZNF207 ORF. Cells were treated with non-targeting siRNA control or siRNA targeting endogenous ZNF207 (siZNF207). Blots were probed with antibodies against ZNF207, and GAPDH (loading control). **(D)** RT-PCR analysis of alternative splicing for *ACLY* (left) and *SRRM1* (right) in HEK293 Flp-In cells treated with ZNF207-targeting siRNA and/or expressing siRNA-resistant N-terminally tagged ZNF207 ORF. Quantification of PSI values from three independent experiments are shown below the gel. Data are presented as mean ± SD. ****p < 0.0001; Dunnett’s multiple comparisons test following one-way ANOVA. **(E)** Enrichment of KEGG biological pathways among genes showing differential expression after ZNF207 depletion, which are rescued by ZNF207 reintroduction (as shown in Figure 4E). Only terms with adjusted p-value < 0.05 are shown. **(F)** Real-time quantitative RT-PCR analysis of transcript levels for *LUC7L*, *LUC7L2*, *RBM3*, and *SNRNP70* in HEK293 Flp-In cells expressing a doxycycline-inducible, siRNA-resistant 3×FLAG C-terminal tagged ZNF207 ORF. Cells were treated with either non-targeting siRNA controlor siRNA targeting endogenous ZNF207 (siZNF207) and further treated with either 0.5 µM SMG1 NMD inhibitor (SMG1i) or DMSO for 6 hours prior to RNA extraction. Quantification of transcript levels from three independent experiments are displayed. Data are presented as mean ± SD. ****p-value < 0.0001, ***p-value < 0.001, **p-value < 0.01, *p-value < 0.05; Šídák’s multiple comparisons testing following one-way ANOVA. **(G)** Genome browser tracks highlighting alternative splicing events in *LUC7L*, *SNRNP70*, and *RBM3*, genes whose expression is regulated by ZNF207. Tracks represent data from two independent RNA-Seq replicates under three conditions: cells treated with control siRNAs (siControl), siRNAs targeting ZNF207 (siZNF207), and siZNF207-treated cells rescued by induced ZNF207 expression. Normalized reads per million (RPM) are displayed on the left of each track. The regulated splicing events are highlighted.

**Supplementary Figure 7:**
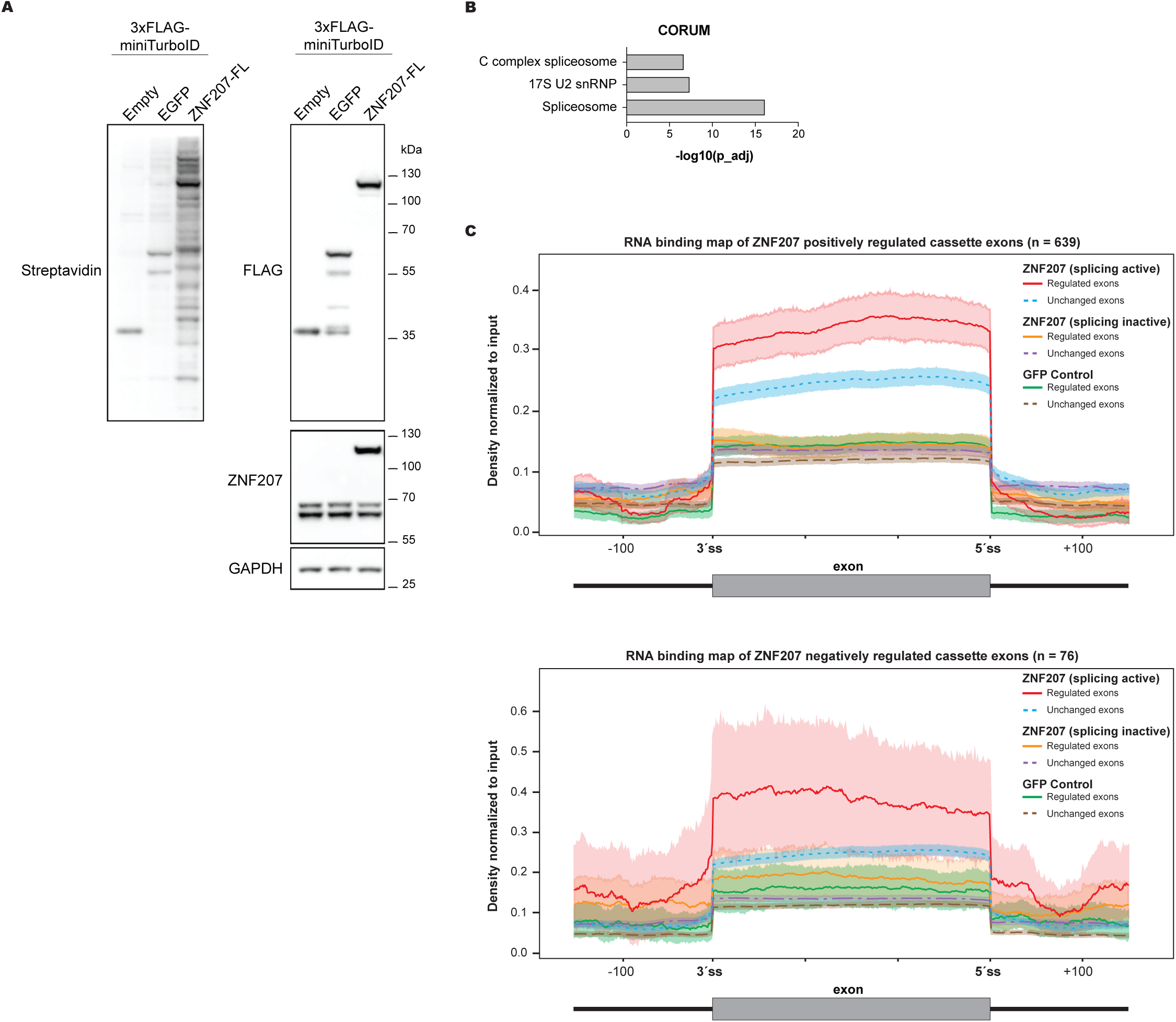
ZNF207 directly impacts alternative splicing. Related to Figure 5. **(A)** Western blot analysis of HEK293 Flp-In cells expressing 3×FLAG-miniTurboID-tagged constructs, including an empty vector, EGFP, or ZNF207 (C-terminal tagged with 3×FLAG-miniTurboID). The blot was probed with streptavidin (for biotinylation detection), ZNF207, FLAG, and GAPDH antibodies. **(B)** CORUM complex enrichment analysis of ZNF207 proximal interactors identified via TurboID experiments. All enriched terms have adjusted p-value < 0.0001. **(C)** Average eCLIP signal profiles for C-terminal FLAG-tagged ZNF207 over exons positively (top; n = 639) or negatively (bottom; n = 76)) regulated by ZNF207 and over unchanged alternative exons (n = 3,745). Signal profiles are compared to the negative control GFP and N-terminal FLAG-tagged ZNF207 (lacking splicing activity) for the same exon subsets.

**Supplementary Figure 8:**
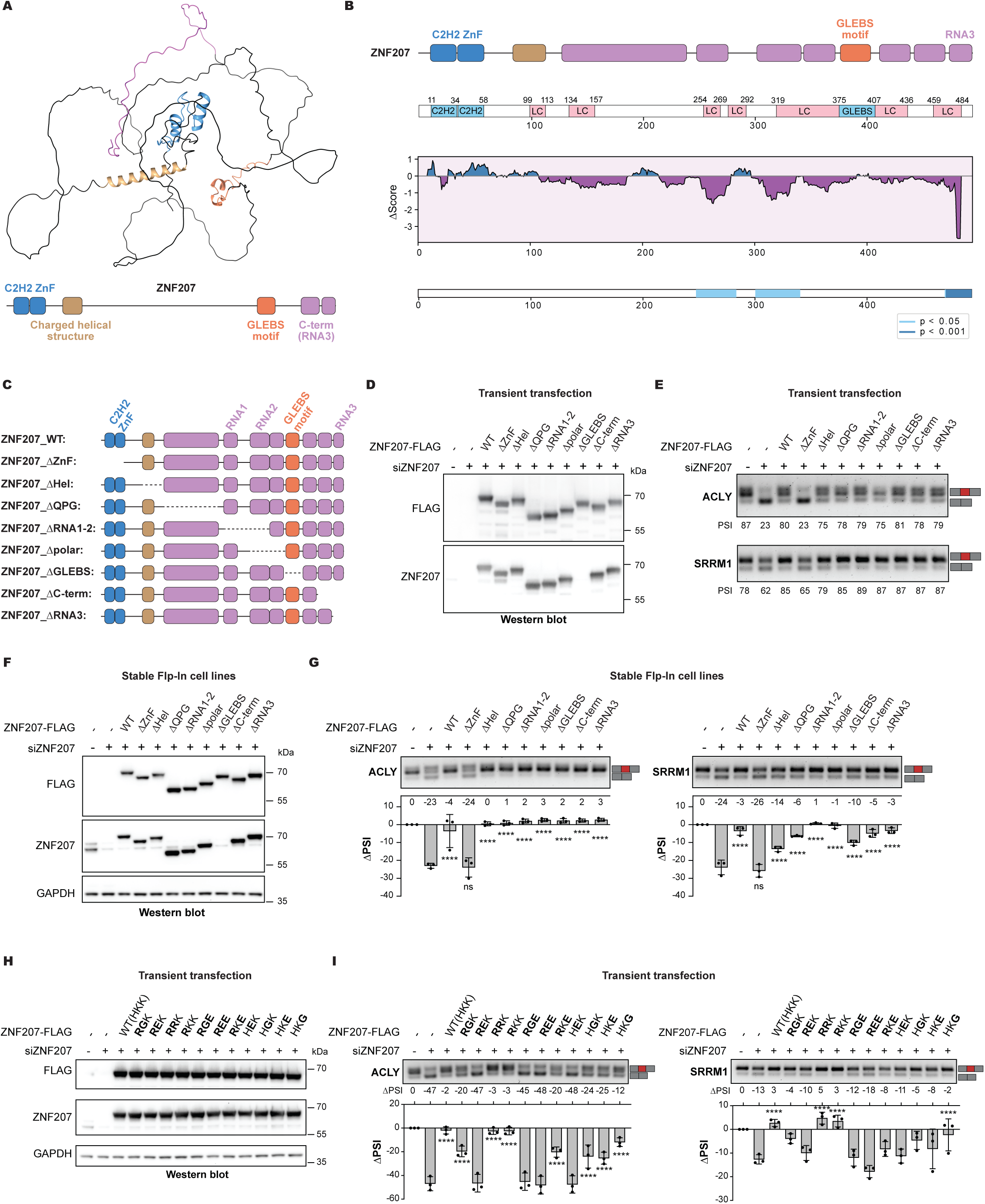
Zinc finger domains are essential for ZNF207 splicing regulatory activity. Related to Figure 6. **(A)** AlphaFold prediction of the full-length ZNF207 protein. Key structural regions are highlighted: the C2H2 zinc finger domain in blue, the helical structure in brown, the GLEBS motif in orange, and the C-terminal regions, predicted to be disordered and RNA-binding, in pink. Corresponding features are similarly annotated in the schematic below. **(B)** HydRA analysis predicting RNA-binding regions within ZNF207. Occlusion maps illustrate regions with differential RNA-binding activity: purple regions represent Δscore < 0, while blue regions represent Δscore > 0. Statistically significant regions (p < 0.05 or p < 0.001) are highlighted in light blue and dark blue, respectively, on the bottom track. The coordinates of protein domains and annotated regions are displayed at the top. **(C)** Schematic representation of ZNF207 truncation mutants used in the figure. **(D)** Western blot analysis of ZNF207 truncation mutants transiently transfected into HEK293T cells. Cells were treated with control siRNA (siControl) or siRNA targeting endogenous ZNF207 (siZNF207). Blots were probed with antibodies against ZNF207, and FLAG. **(E)** RT-PCR analysis of ACLY (top) and SRRM1 (bottom) alternative splicing in HEK293 cells treated with ZNF207-targeting siRNA and transiently transfected with siRNA-resistant, C-terminally 3×FLAG-tagged ZNF207 truncation mutants. Quantification of PSI values are shown below the gel. **(F)** Western blot analysis of siRNA-resistant 3×FLAG C-terminal tagged ZNF207 truncation mutants expressed in HEK293 Flp-In cells under doxycycline induction. Cells were treated with non-targeting siRNA control or siRNA targeting endogenous ZNF207 (siZNF207). Western blots were probed with antibodies against ZNF207, FLAG (to detect tagged constructs), and GAPDH (loading control). **(G)** RT-PCR analysis of ACLY (left) and SRRM1 (right) alternative splicing in HEK293 Flp-In cells treated with ZNF207-targeting siRNA and/or expressing siRNA-resistant wild-type or truncation mutant ZNF207 ORFs. Quantification of ΔPSI values from three independent experiments are shown below the gel. Data are presented as mean ± SD. Statistical comparisons were performed relative to the siZNF207 sample without ZNF207 ORF expression. ****p < 0.0001; Dunnett’s multiple comparisons test following one-way ANOVA. **(H)** Western blot analysis of ZNF207 mutants at residues H41, K42, and K43 (wild-type: HKK), transiently expressed in HEK293T cells. Cells were transfected with either control siRNA (siControl) or siRNA targeting endogenous ZNF207 (siZNF207). Blots were probed with antibodies against ZNF207, FLAG, and GAPDH (loading control). **(I)** RT-PCR analysis of ACLY (left) and SRRM1 (right) alternative splicing in HEK293 cells treated with ZNF207-targeting siRNA and transiently transfected with siRNA-resistant, C-terminally 3×FLAG-tagged ZNF207 mutants. Quantification of ΔPSI values from three independent experiments are shown below the gel. Data are presented as mean ± SD. Statistical comparisons were performed relative to the siZNF207 sample without ZNF207 ORF expression. ****p < 0.0001; Dunnett’s multiple comparisons test following one-way ANOVA.

**Supplementary Figure 9:**
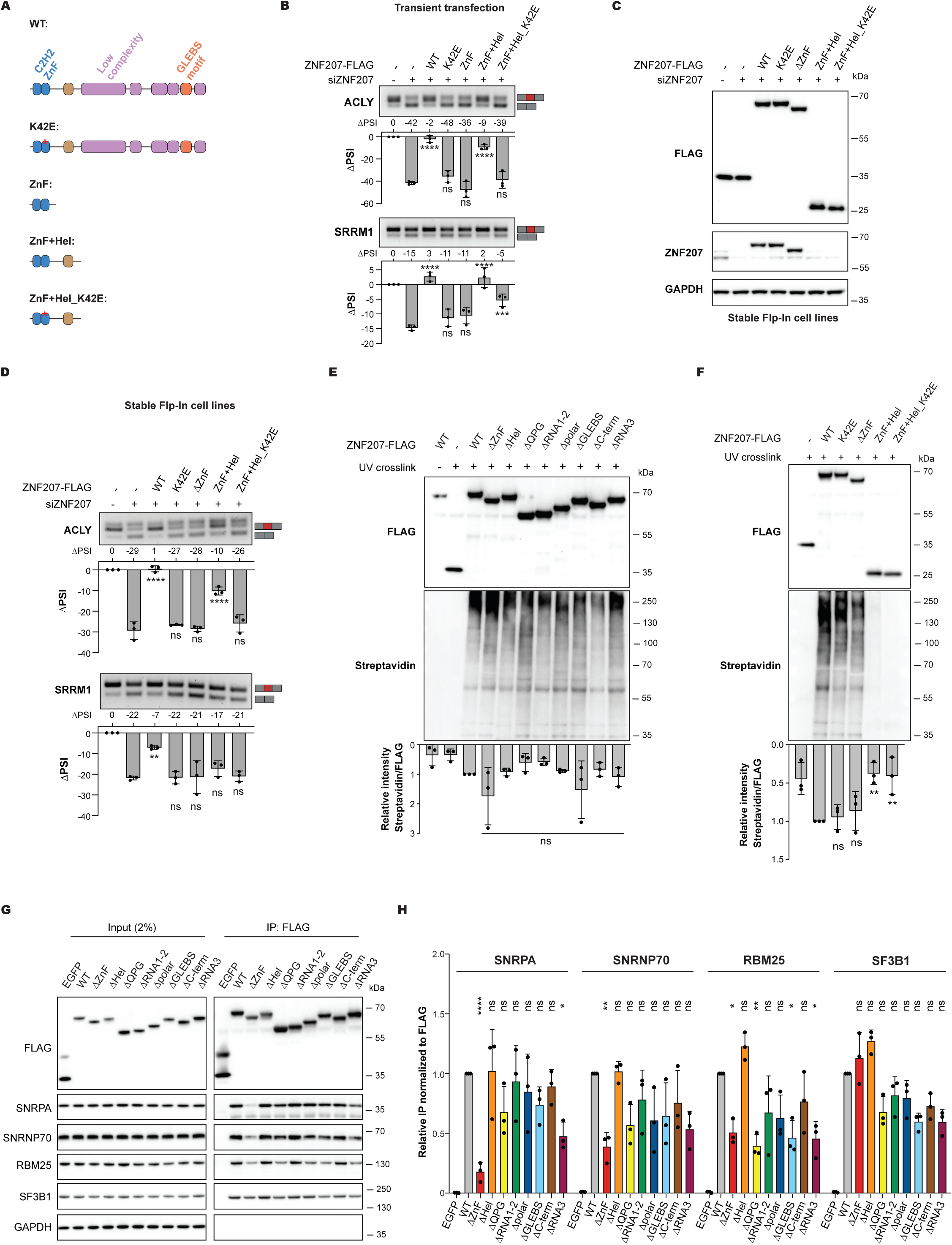
ZNF207 zinc finger domain facilitates interactions with U1 snRNP components. Related to Figure 6. **(A)** Schematic representation of ZNF207 variants. **(B)** RT-PCR analysis of ACLY (top) and SRRM1 (bottom) alternative splicing in HEK293 cells treated with ZNF207-targeting siRNA and transiently transfected with siRNA-resistant, C-terminally 3×FLAG-tagged ZNF207 mutants. Quantification of ΔPSI values from three independent experiments are shown below the gel. Data are presented as mean ± SD. Statistical comparisons were performed relative to the siZNF207 sample without ZNF207 ORF expression. ****p < 0.0001; Dunnett’s multiple comparisons test following one-way ANOVA. **(C)** Western blot analysis of siRNA-resistant 3×FLAG C-terminal tagged ZNF207 mutants expressed in HEK293 Flp-In cells under doxycycline induction. Cells were treated with non-targeting siRNA control or siRNA targeting endogenous ZNF207 (siZNF207). Western blots were probed with antibodies against ZNF207, FLAG (to detect tagged constructs), and GAPDH (loading control). **(D)** RT-PCR analysis of ACLY (top) and SRRM1 (bottom) alternative splicing in HEK293 Flp-In cells treated with ZNF207-targeting siRNA and/or expressing siRNA-resistant wild-type or mutant ZNF207 ORFs. Quantification of ΔPSI values from three independent experiments is shown below the gels. Data are presented as mean ± SD. Statistical comparisons were performed relative to the siZNF207 sample without ZNF207 ORF expression. ****p < 0.0001; Dunnett’s multiple comparisons test following one-way ANOVA. **(E-F)** Biotin-based RNA labeling of UV-crosslinked RNA bound to immunoprecipitated ZNF207 truncation mutants. Pull-down efficiency of each construct was assessed by FLAG Western blot. Quantification of relative RNA signal from three independent experiments is shown below the gel. Data are presented as mean ± SD. Statistical analysis was performed using one-way ANOVA followed by Dunnett’s multiple comparisons test. **(G)** Representative western blot analysis of total cell lysates (input) treated with benzonase, and FLAG immunoprecipitates (IP: FLAG-M2) from HEK293 Flp-In cells expressing 3×FLAG-tagged ZNF207 truncation mutants shown in S8C. Blots were probed with antibodies specific for FLAG, SNRPA, SNRNP70, RBM25, SF3B1, and GAPDH (negative control). **(H)** Quantification of co-immunoprecipitated spliceosome components, normalized to FLAG-ZNF207 pulldown efficiency and wild-type (WT) ZNF207 levels. Data are presented as mean ± SD. Statistical comparisons were performed relative to WT-ZNF207. ***p < 0.001, **p < 0.01, *p < 0.05; Šídák’s multiple comparisons test following two-way ANOVA.

**Supplementary Figure 10:**
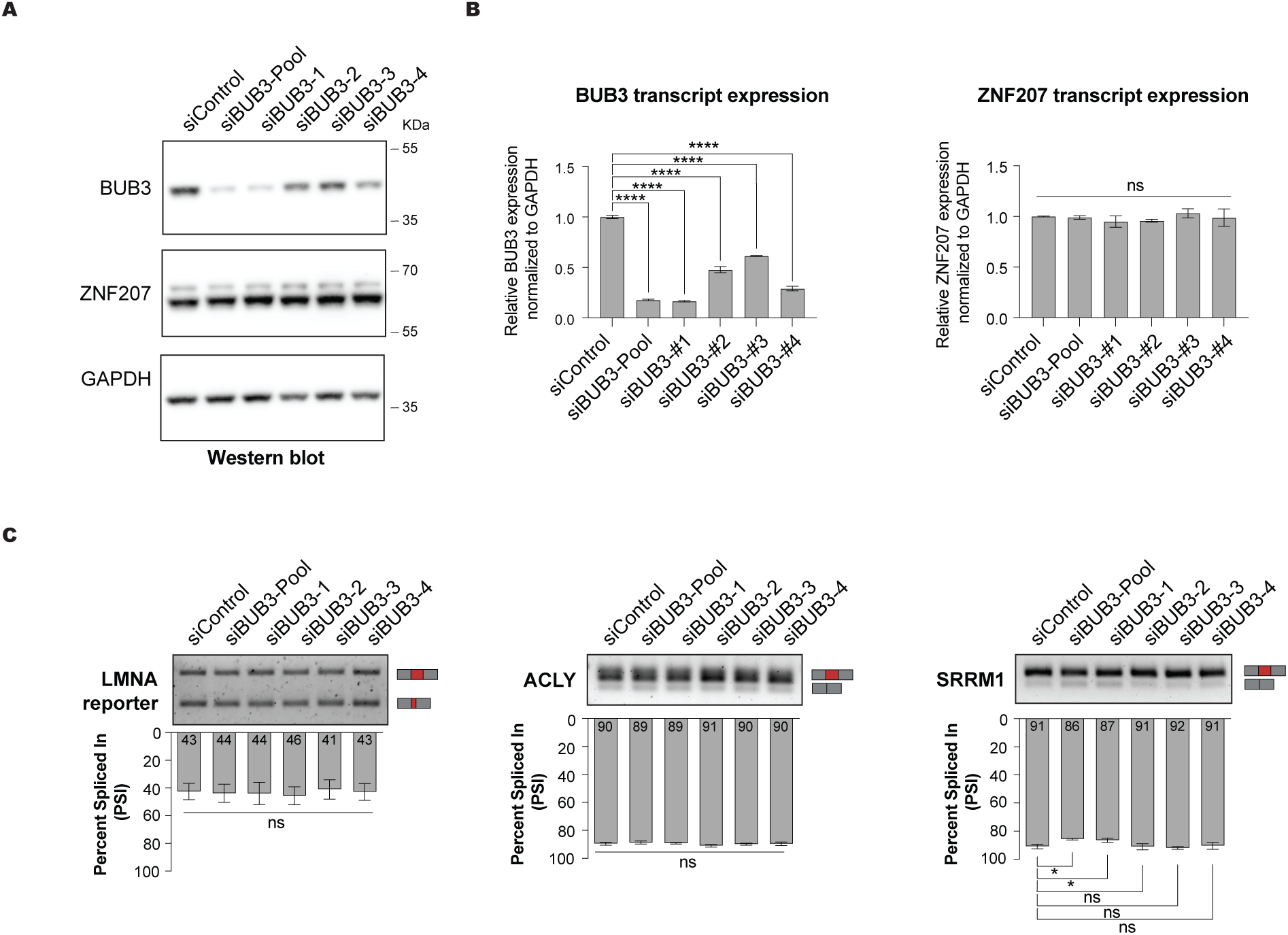
BUB3 knockdown has minimal impact on ZNF207-driven splicing. Related to Discussion. **(A)** Western blot analysis of BUB3 and ZNF207 in HEK293T cells treated with non-targeting siRNA control (siCTL), four independent siRNAs targeting BUB3 (siBUB3), or a pool of these siRNAs. Blots were probed with antibodies against BUB3, ZNF207, and GAPDH (loading control). **(B)** Real-time quantitative RT-PCR analysis of BUB3 and ZNF207 transcript levels in HEK293T cells treated with either control siRNA (siCTL) or siRNAs targeting BUB3 (siBUB3), as in panel B. Quantification of transcript levels from three independent experiments are displayed. Data are presented as mean ± SD. ****p-value < 0.0001; one-way ANOVA with Dunnett’s multiple comparisons test. **(C)** RT-PCR analysis of *LMNA* minigene reporter (left), endogenous *ACLY* (middle), and endogenous *SRRM1* (right) alternative splicing in HEK293T cells treated with BUB3-targeting siRNAs, as in panel B. Quantification of PSI values from three independent experiments are shown below the gel. Data are presented as mean ± SD. *p-value < 0.05; one-way ANOVA with Dunnett’s multiple comparisons test.

## SUPPLEMENTARY TABLE LEGENDS

**Table S1: Genome-wide CRASP-Seq CHyMErA Library. Related to Figure 1**.

Comprehensive annotation of the CHyMErA genome-wide knockout library. Includes gRNA sequences, target gene information, gRNA classification, guide sequences, predicted cut sites, on- and off-target score evaluations, fraction of targeted transcript isoforms, relative position of targeted site with the coding sequence for both Cas9 and Cas12a gRNA sequence of each hgRNA. The designed 187-nucleotide oligo pool used for cloning is included.

**Table S2: Oligos, DNA Fragments, and Addgene Plasmids Used in This Study. Related to Methods.**

(Sheet 1) Sequences of oligonucleotides utilized in this study, including primers for RT-PCR, qPCR, and gRNA sequences.

(Sheet 2) Sequences of oligonucleotides utilized in this study for CRASP-Seq Illumina library preparation.

(Sheet 3) Sequences of the six minigene splicing reporters included in our CRASP-Seq libraries. Intronic regions are represented in lowercase letters.

(Sheet 4) Details of newly cloned plasmids and libraries deposited to Addgene.

**Table S3: Genome-wide knockout CRASP-Seq data. Related to Figure 2**.

(Sheet 1) ΔPSI values for each hgRNA across 25 screens conducted in this study.

(Sheet 2) ΔPSI values for 370 gene hits identified across the five splicing reporter screens.

(Sheets 3-7) ΔPSI values for gene hits from individual splicing reporter screens: FAS (Sheet 3), EZH2

(Sheet 4), PKM (Sheet 5), SRSF7 (Sheet 6), and LMNA (Sheet 7).

**Table S4: Transcriptome-wide Effects of ZNF207 on Alternative Splicing. Related to Figure 4**.

(Sheet 1) Whippet analysis of alternative splicing changes in HEK293T cells following ZNF207 knockdown with three independent siRNAs. Includes gene ID, event coordinates, event size, event type, gene type, gene FPKM value, absolute PSI, ΔPSI, and probability scores for endogenous splicing events.

**Table S5: Transcriptome-wide Effects of ZNF207 on Alternative Splicing. Related to Figure 4**.

(Sheet 1) Whippet analysis of alternative splicing changes in HEK293 Flp-In cells following ZNF207 knockdown and rescue with ZNF207 open reading frame (ORF). Includes gene ID, event coordinates, event size, event type, gene type, gene FPKM value, absolute PSI, ΔPSI, and probability scores for endogenous splicing events.

(Sheet 2) List of ZNF207 regulated events in protein coding genes. Includes gene ID, event coordinates, event size, event type, gene type, gene FPKM value, absolute PSI, ΔPSI, and probability scores for endogenous splicing events.

FLAG EV: FLAG empty vector.

**Table S6: Transcriptome-wide Effects of ZNF207 on Gene Expression. Related to Figure 4**.

(Sheet 1) Results from DESeq2 analysis of gene expression in HEK293 Flp-In cells following ZNF207 knockdown and subsequent rescue with the ZNF207 ORF. The table includes gene names, gene IDs, gene types, log2 fold-change (LFC) values, and adjusted p-values.

(Sheet 2) List of ZNF207 regulated protein-coding genes. The table includes gene names, gene IDs, gene types, log2 fold-change (LFC) values, and adjusted p-values.

FLAG EV: FLAG empty vector.

**Table S7: ZNF207 miniTurboID data. Related to Figure 5**.

Summary of identified peptides and their counts from ZNF207 miniTurboID proximity labeling mass spectrometry analysis.

(Sheet 1) Comprehensive list of all proteins and peptide counts identified in the miniTurboID MS data.

(Sheet 2) Filtered dataset of ZNF207 proximal interactors consistently identified across all three replicates.

**Table S8: Base Editing CRISPR Screens Identifying ZNF207 Regions Critical for Splicing Activity. Related to Figure 6**.

(Sheet 1) Detailed list of all Cas9 sgRNAs used in the base editor high-throughput mutagenesis CRASP-Seq screen. Each entry includes the gene name, gene ID, Cas9 guide sequence, guide coordinates, and on- and off-target scores. The 144-nucleotide oligo pool designed for cloning is also provided.

(Sheet 2) Base editing screen data, including the position of each guide with the potential to mutate adenines (A) or cytosines (C), and the length of guides overlapping coding sequences (CDS). The table specifies the targeted amino acid positions for each guide, determined based on the longest protein and transcript isoforms (both specified). The amino acid corresponding to position 1 of each guide is also noted. Additionally, the table includes PSI and ΔPSI values for each gRNA in the library and for each targeted gene.

## ACKNOWLEDGEMENTS

The authors thank Michael Aregger, Colin Wu, Moonsup Lee, and Brian Yee, for constructive discussions and technical assistance. We also thank members of the Gonatopoulos-Pournatzis and Aregger groups, as well as the RNA Biology Laboratory at NCI-Frederick. We are also grateful to the CCR Sequencing Facility, as well as the NIH/NCI/CCR High Throughput Imaging Facility (HiTIF) for help with high throughput imaging and Gianluca Pegoraro (NIH/NCI/CCR HiTIF) for help with writing the R code for high throughput image analysis. The NMD inhibitor SMG1i was kindly provided by the Cystic Fibrosis Foundation Chemical Compound Program. Figures 1A, 6A, 7 and the graphical abstract include elements from BioRender.com. This work was supported by the NCI/NIH Intramural Research Program (Project ZIA BC012019).

## AUTHOR CONTRIBUTIONS

Conceptualization and study design: T.G.-P.; Data curation: J.J.K., S.K., T.G.-P., A.P.D., S.V., A.K.B., B.K., E.D., M.-S.X., G.D.; Formal Analysis: A.K.B., J.J.K., T.G.-P., B.K., T.A., A.P.D., G.D., S.V.; Funding acquisition: T.G.-P., T.M.; Supervision: T.G.-P., T.M.; Visualization: A.K.B., J.J.K., T.G.-P., S.K., B.K., A.P.D., S.V.; Writing – original draft: T.G.-P.; Writing – review & editing: A.K.B., J.J.K., T.M., A.P.D., T.A., S.V., B.K., G.D..

## RESOURCE AVAILABILITY

### Lead contact

For additional details and inquiries regarding resources and reagents, please reach out to the lead contact, Thomas Gonatopoulos-Pournatzis.

### Materials availability

All generated key plasmids and libraries have been deposited to Addgene (see Table S2).

Plasmids: https://www.addgene.org/depositing/85289/

Libraries: https://www.addgene.org/pooled-library/gonatopoulos-pournatzis-human-crispr-crasp-seq/

Any other reagent is available upon reasonable request.

### Data availability

- CRASP-Seq screening data generated by this study have been deposited at GE Omnibus (GEO) and are available. Accession number is: GSE284628.
- The GEO accession numbers for the ZNF207 related RNA-Seq data are: GSE284629 and GSE284630.
- The accession number for the eCLIP-Seq data is: [GSE284631](https://www.ncbi.nlm.nih.gov/geo/query/acc.cgi?acc=GSE284630).
- The ZNF207 mass spectrometry data associated with this study have been deposited to the ProteomeXchange consortium through partner MassIVE (massive.ucsd.edu): [MassIVE Dataset Summary](https://massive.ucsd.edu/ProteoSAFe/private-dataset.jsp?task=4c8496f4558d4612a12414ee5e12be77) (Submission ID: **MSV000096693).**

## METHODS SECTION

### Cell culture

HEK293T, HEK293 Flp-In, HAP1, and RPE1 cells were maintained in Dulbecco’s Modified Eagle’s Medium (DMEM; Gibco #11995073) supplemented with 10% heat-inactivated fetal bovine serum (HI-FBS; Gibco #16140071) and 1% penicillin-streptomycin (Gibco #15140122). HepG2 cells were cultured in Eagle’s Minimum Essential Medium (EMEM; ATCC #30-2003) with 10% fetal bovine serum (FBS; Gibco #A5670701). hTERT-immortalized fibroblasts, including wild-type (WT) and Hutchinson-Gilford Progeria Syndrome (HGPS) patient-derived cells, were cultured in Minimum Essential Medium (MEM; ThermoFisher Scientific #11090081) supplemented with 10% HI-FBS, 1% penicillin-streptomycin, and 2 mM L-glutamine (ThermoFisher Scientific #25030081). UOK124 cells were cultured in DMEM supplemented with 1% non-essential amino acids (MEM; Gibco #11140050), 10% HI-FBS; Gibco and 1% penicillin-streptomycin. All cell lines were grown at 37°C in a humidified 5% CO₂ atmosphere. Cells were routinely passaged using 0.25% trypsin-EDTA (Gibco #25200056) and seeded at appropriate densities for each experimental condition in tissue culture-treated plates. The culture medium was refreshed every 2–3 days, and cell confluency was monitored with an EVOS™ M5000 Imaging System (Invitrogen). Cell viability was assessed using a Vi-Cell BLU Cell Viability Analyzer (Beckman Coulter).

### Generation of HEK293 Flp-In cell lines expressing ZNF207

The CHyMErA and base editor stable cell lines were generated and maintained as described previously.^11^ To generate HEK293T cells stably expressing the LMNA minigene reporter, 350,000 HEK293T cells were seeded and transduced with lentiviral supernatant in growth medium supplemented with polybrene (1:1000 dilution) to enhance infection efficiency. Approximately 22 hours post-transduction, puromycin was added to the medium to select for stable integrands.

For generating Flp-In cell lines expressing ZNF207, one million HEK293 Flp-In cells were transfected with pcDNA5 ZNF207 plasmids or controls using the Lipofectamine 2000 transfection reagent (ThermoFisher Scientific #11668027) following manufacturer’s recommendation. After 4.5 hours, the transfection media was replaced with fresh growth media to reduce Lipofectamine-associated toxicity. Two days post-transfection, cells were treated with 200 µg/mL Hygromycin B (ThermoFisher Scientific #10687010) in DMEM. The hygromycin-containing media was refreshed every 4–5 days to select for stably transfected cells. Once cells reached confluency, various concentrations of doxycycline were tested to optimize expression of FLAG-tagged proteins. Final concentrations of 0.001 µg/mL doxycycline for FLAG-Empty or FLAG-EGFP and 0.5 µg/mL doxycycline for FLAG-ZNF207 constructs were selected for consistent protein expression.

### Cloning CRASP-Seq lentiviral vectors

The CRASP-Seq lentiviral vector was constructed using the pLCHKOv3 backbone (Addgene #209025) as a starting point. First, the two existing BpiI restriction sites, intended for the subsequent two-step library cloning, were removed via site-directed mutagenesis. Second, the hgRNA expression cassette was inverted from the minus to the plus strand orientation. Third, a minigene cloning sequence, including a FLAG-tag and followed by unique Eco47III and Eco32I restriction sites (inserted for the subsequent cloning of cell barcodes and minigene reporters), was inserted downstream of the hgRNA expression cassette on the minus strand. Fourth, a doxycycline-inducible tetracycline response element (TRE) was cloned on the minus strand between the hgRNA expression cassette and the minigene cloning sequence. Fifth, 2A peptide sequence fused to rtTA ORF was inserted immediately upstream of the stop codon of the puromycin resistance cassette. Finally, a BGH poly(A) signal was incorporated into the minus strand upstream of the hgRNA expression cassette (Figure 1A). This CRASP-Seq backbone vector, named pLCHKO-CRASP, has been deposited to Addgene (#231928).

Using pLCHKO-CRASP, we applied NEBuilder HiFi DNA Assembly Master Mix (New England Biolabs #E2621L) to clone a cell barcode library containing 12 random nucleotides. A 62-nucleotide (nt) oligonucleotide, ordered from IDT, included 25-nt flanking sequences around the 12 random nucleotides at the Eco47III (ThermoFisher Scientific # FD0324) restriction site. The Eco47III-digested vector was used in 16x NEBuilder reactions following the manufacturer’s protocol. The reaction mixture was precipitated with sodium acetate and ethanol, as previously described^34^, and the purified product was transformed into Endura competent cells (LGC Biosearch Technologies #60242-2) by electroporation (1 mm cuvette, 25 µF, 200 Ω, 1,600 V). Sufficient cells were plated on 15-cm LB agar plates containing 100 µg/mL carbenicillin to achieve a library coverage of 2,000-fold. After overnight incubation at 30°C, colonies were scraped, pooled, and the bacterial pellets were collected. Plasmids were extracted using the EndoFree Plasmid Maxi Kit (QIAGEN #12362), and the resulting library plasmids were deposited to Addgene (Addgene #232068 – CRASP-Seq CBC backbone).

The genome-wide CHyMErA KO library was constructed as described previously.^11^ Briefly, a single oligo pool of 187 nt, including a 20-nt Cas9 guide sequence, followed by the Cas9 tracrRNA, the AsCas12a direct repeat (DR), and a 23-nt Cas12a guide sequence, was synthesized by TWIST Biosciences (**Table S1**). These guide sequences were flanked by BveI/BspMI restriction sites. The oligo pool, comprising 95,893 guide sequences, was amplified by PCR using KAPA HiFi HotStart DNA polymerase (Roche #KK2601). Four 50 µL PCR reactions were set up with 10 nM oligo pool and 0.35 µM of each primer (**Table S2**) under the following conditions: initial denaturation at 98°C for 3 minutes, followed by 10 cycles of 98°C for 10 seconds, 62°C for 15 seconds, and 72°C for 20 seconds. Amplified oligos were purified using a PCR purification kit (ThermoFisher Scientific #K0701), and quality was confirmed via 2% agarose gel electrophoresis. Oligos were then digested with BveI (ThermoFisher Scientific #FD1744) and ligated into 2 µg of digested CRASP-Seq CBC backbone using T4 DNA ligase (New England Biolabs #M0202) in a Golden Gate reaction (step 1: 37°C for 30 minutes; step 2: 37°C for 30 minutes, 22°C for 30 minutes, repeated for 16 cycles; step 3: 37°C for 15 minutes; step 4: 65°C for 20 minutes) at a 1:9 vector-to-insert molar ratio. The ligation mix was precipitated and purified, and Endura competent cells were transformed by electroporation as described above to achieve a library coverage of >1,000-fold. After incubation at 30°C, colonies were pooled, and the plasmid library was extracted using the EndoFree Plasmid Maxi Kit (QIAGEN #12362). The library plasmids were deposited to Addgene (#232069 – CRASP-Seq Gene KO v1 empty).

As a final step, we cloned various minigene reporters into the CRASP-Seq Gene KO v1 empty library. DNA fragments containing the desired sequences, including complete native intronic sites and neighboring exons, were either synthesized (TWIST Biosciences) or amplified from genomic DNA. NEBuilder was used to insert these fragments into the Eco32I-digested CRASP-Seq Gene KO v1 empty vector with reactions yielding at least 250-fold coverage as described above.

### Cloning base editor library

2.5 µg of CRASP-Seq CBC backbone plasmid was digested using the BveI restriction enzyme at 37°C for 2 hours in a total of eight reactions (total plasmid = 20 µg). The 144-nucleotide gRNA library oligo pool (TWIST Biosciences; Table S8) was amplified in a thermocycler using the following protocol: initial denaturation at 95°C for 3 minutes, followed by six cycles of 98°C for 15 seconds, 65°C for 15 seconds, and 65°C for 45 seconds using KAPA HiFi HotStart DNA polymerase. The TKO_F1 and TKO_R2 primers (Table S2) were used for this amplification. Both the digested plasmids and the amplified gRNA library oligos underwent PCR purification using the QIAquick PCR Purification Kit (QIAGEN #28104). Purified plasmids were then further purified via gel purification using GeneJET Gel Extraction Kit (ThermoFisher Scientific # K0691).

Golden gate cloning was performed with ∼66 fmol of purified plasmids and ∼330 fmol of large-scale oligos, followed by ethanol precipitation and reconstitution in 10 µL of TE buffer. For transformation, 90 µL of Endura electrocompetent cells were used, following the manufacturer’s protocol, and plated on Carbenicillin (100 µg/mL) agar plates. The calculated library coverage was approximately 3,500x. Colonies were collected and pelleted, and the CRASP-Seq BE Tiling v1 empty library (Addgene # 232070) was purified using the EndoFree Plasmid Maxi Kit (QIAGEN #12362).

We next cloned the oligo library into the LMNA minigene reporter. 20 µg of the CRASP-Seq BE Tiling v1 empty library were digested with Eco32I at 37°C for 2 hours, followed by PCR and gel purification (ThermoFisher Scientific # K0691). The LMNA mutant splicing reporter (Table S2; ∼46 fmol per reaction) was inserted into the digested plasmid (∼16 fmol per reaction) using the NEBuilder HiFi DNA assembly reaction (8 reactions; NEB, Cat# E2621L), followed by ethanol precipitation and reconstitution in TE buffer. Subsequently, 120 µL of Endura electrocompetent cells were transformed with the assembled constructs and plated on Carbenicillin agar plates. After colony collection and pelleting, the LMNA splicing reporter plasmid library was obtained by the EndoFree Plasmid Maxi Kit (QIAGEN #12362).

### Lentivirus production

Lentivirus production was carried out as described previously.^11^ For gRNA library virus production, 8 million HEK293T cells were seeded per 15-cm plate in DMEM with 10% HI-FBS. Twenty-four hours post-seeding, cells were transfected with a mix containing 8 μg of the lentiviral CRASP-Seq library vector, 6.5 μg of packaging plasmid psPAX2 (Addgene #12260), 4 μg of envelope plasmid pMD2.G (Addgene #12260), 48 μL of X-tremeGENE 9 DNA Transfection Reagent (Sigma-Aldrich #6365809001), and 1.4 mL of Opti-MEM (Gibco #31985062). After 24 hours, the medium was replaced with serum-free, high-BSA growth medium (DMEM with 1.1 g/100 mL BSA and 1% penicillin-streptomycin). Virus-containing medium was collected 48 hours post-transfection, centrifuged at 475 rcf for 5 minutes at 4°C, aliquoted, and stored at −80°C.

For viral titer determination, target cells were transduced with serial dilutions of the lentiviral hgRNA library in the presence of polybrene (8 μg/mL; Sigma-Aldrich #H9268). After 24 hours, the virus-containing medium was replaced with fresh medium containing puromycin (1–2 μg/mL; ThermoFisher Scientific #A1113803). Cells were incubated for an additional 48 hours. The multiplicity of infection (MOI) was determined 72 hours post-infection by comparing the survival rate of puromycin-selected cells to infected but non-selected control cells.

For producing lentivirus with the LMNA mutant splicing reporter, 750,000 HEK293T cells were seeded in a 6-well plate and subsequently transfected with 2.5 μg of the lentiviral vector containing the LMNA mutant splicing reporter, 600 ng of psPAX2, and 400 ng of pMD2.G using the Lipofectamine 2000 transfection reagent (ThermoFisher Scientific #11668027), following the manufacturer’s protocol.

### RT-PCR assays

RNA was extracted from frozen cell pellets using RNeasy Plus Universal Mini Kit (QIAGEN #74136) according to manufacturer’s protocol. 40 ng of RNAs were used for RT-PCR reaction using QIAGEN OneStep RT-PCR Kit (QIAGEN #210215). The primers are provided in Table S2. For RT-PCR targeting LMNA, ACLY and SRRM1, 30 cycles were used. 26 cycles were used to amplify LMNA splicing reporter. For RT-PCR targeting LUC7L, LUC7L2, SNRNP70, and RBM3, 100 ng of RNAs were used. 29 cycles were used for LUC7L. 31 cycles were used for LUC7L2. 32 cycles were used for SNRNP70 and RBM3.

### Immunoblotting

For experiments with HEK293T and HEK293 Flp-In cells, cell pellets were collected and lysed on ice for 10 minutes in F buffer (10 mM Tris pH 7.05, 50 mM NaCl, 30 mM Na4 pyrophosphate, 50 mM NaF, 5 µM ZnCl2, 10% glycerol, 0.5% Triton X-100), supplemented with a protease inhibitor cocktail (Roche #11836170001). Lysates were then centrifuged at 18,500 rcf for 5 minutes at 4°C, and the supernatants were collected. Samples were prepared by heating at 70°C for 5 minutes in NuPAGE LDS Sample Buffer (ThermoFisher Scientific #NP0007) with DTT (100 mM; ThermoFisher Scientific #R0861). Protein concentrations were determined using Bradford reagent (Bio-Rad #5000006), and equal amounts of protein (10–30 µg) were loaded and separated on 4–12% Bis-Tris gels (ThermoFisher Scientific #NP0323BOX). Proteins were then transferred to PVDF membranes, blocked with 5% milk for 1 hour at room temperature, and probed with primary antibodies including ZNF207 (1:2,000, Invitrogen #703747), FLAG M2 (1:2,000, Sigma #F3165), GAPDH (1:2500, Proteintech #10494-1-AP), SF3B1 (1:2,000, Proteintech #27684-1-AP), PRPF6 (1:1,000, Proteintech #23929-1-AP), SNRNP200 (1:1,000, Proteintech #23875-1-AP), RBM25 (1:2,000, Proteintech #25297-1-AP), SNRNP70 (1:1,000, Abcam #ab83306), SNRPA (1:2,000, Proteintech #10212-1-AP), SNRPF (1:1,000, Proteintech #14977-1-AP), and BUB3 (1:1,000, FisherFisher Scientific #BDB611730). All primary antibodies were prepared in 5% milk unless noted otherwise. After primary antibody incubation, membranes were washed and incubated with HRP-conjugated secondary antibodies (anti-Rabbit, Cell Signaling Technology #7074; anti-Mouse, Cell Signaling Technology #7076) at a 1:5,000 dilution for 1 hour at room temperature. Following additional washes, chemiluminescence detection was performed using SuperSignal West Pico PLUS (ThermoFisher Scientific #34580) or Femto (ThermoFisher Scientific #A45916), and images were captured using the Invitrogen iBright CL1500 Imaging System (ThermoFisher Scientific # A44114).

For HGPS patient-derived immortalized fibroblast cells, equal cell numbers (25,000–50,000) were pelleted and lysed in F buffer (1–2 µL of lysis buffer per 1,000 cells) with a protease inhibitor cocktail. After a 10-minute ice incubation, lysates were sonicated using a Bioruptor Plus (Diagenode # B01020002) at high power for 14 cycles (20 seconds on, 10 seconds off). Sonicated lysates were then mixed with NuPAGE LDS Sample Buffer (ThermoFisher Scientific #NP0007) with DTT (100 mM; ThermoFisher Scientific #R0861) and heated at 95°C for 5 minutes. The initial sample volume was used to determine the amount of beta-actin (1:5,000 in 5% milk; Proteintech #20536-1-AP), while loading volumes for progerin detection were adjusted based on beta-actin levels using Lamin A/C antibody (1:10,000 in 5% BSA; Santa Cruz Biotechnology #sc376248).

To assess biotinylation by miniTurbo Tag, proteins transferred to PVDF membranes were blocked for 1 hour in 5% BSA. The membranes were then incubated with Streptavidin-HRP (1:5,000 in 5% BSA; ThermoFisher Scientific #SA10001) for 1 hour at room temperature or with specific primary antibodies overnight at 4°C. Following washes, chemiluminescence detection was performed as described, with images acquired using the iBright CL1500 Imaging System.

### Co-immunoprecipitation

Six million HEK293 Flp-In cells expressing FLAG-EGFP, FLAG-ZNF207 WT, or FLAG-ZNF207 truncated mutants were plated and treated with doxycycline (0.5 µg/mL) for 20 hours. After treatment, cells were washed with PBS, centrifuged, and the resulting cell pellets were lysed in IGEPAL buffer (50 mM Tris-HCl, pH 7.5, 150 mM NaCl, 1% IGEPAL CA-630, 5% glycerol, with protease inhibitors) and incubated on a rotator at 4°C for 15 minutes. For RNA degradation, 125 units of benzonase were added to each lysate. For immunoprecipitation, protein lysates were incubated at 4°C for 4.5 hours with anti-FLAG M2 Magnetic Beads (Sigma-Aldrich #M8823). After incubation, the beads were washed four times with fresh IGEPAL buffer to remove non-specifically bound proteins. The bound protein complexes were then eluted by boiling in NuPAGE LDS Sample Buffer (ThermoFisher Scientific #NP0007) supplemented with DTT (100 mM; ThermoFisher Scientific, #R0861) at 70°C for 10 minutes at 1,200 rpm. The eluted proteins were analyzed by western blot using the indicated primary antibodies.

### Cloning ZNF207

ZNF207 (ENST00000394670) was cloned into the pDONR223 entry vector using BP Clonase II Enzyme Mix (ThermoFisher Scientific #11789020) following PCR amplification and purification of the target sequences. The BP reaction product was transformed into NEB Stable Competent *E. coli* cells (New England Biolabs #C3040H) with transformants selected on LB agar plates containing spectinomycin. The resulting pDONR223-ZNF207 construct was then modified by site-directed mutagenesis using the Q5 Site-Directed Mutagenesis Kit (NEB #E0552S) to introduce silent mutations for resistance to the most effective ZNF207-targeting siRNA (#1: CAACTAGTGCAACCAGTAA).

Next, the modified ZNF207 sequences were transferred from the pDONR223 entry vector to the pcDNA5-miniTurboID and pcDNA5-3xFLAG destination vectors via Gateway LR Clonase II Enzyme Mix (ThermoFisher Scientific #11791020). For the pcDNA5-3xFLAG constructs, vectors enabling either N-terminal or C-terminal FLAG tags were used. The LR reaction mix was transformed into NEB Stable Competent *E. coli* cells, and transformants were selected on LB agar plates containing 100 µg/mL ampicillin. Only the functional pcDNA vector encoding C-terminal FLAG-tagged ZNF207 was deposited to Addgene (Addgene #231929 – pcDNA5-ZNF207-3xFLAG).

Site-directed mutagenesis of the siRNA-resistant pDONR-ZNF207 plasmid was then performed to generate truncated mutant constructs of ZNF207, including ΔZnF, ΔMB, ΔQPGM, ΔRNA1-2, ΔGLEBS, ΔCterm, and ΔRNA3 (Figure 6D). Domain boundaries for ZnF (zinc finger), MB, QPGM, and GLEBS were assigned based on UniProt annotations (uniprot.org), while the RNA1-2 and RNA3 domains were mapped using HydRA^80^, and the C-terminal domain was predicted based on structural modeling with AlphaFold.^96,97^ All the destination vectors are submitted to Addgene (see Table S2 for Addgene IDs).

For lentiviral vector construction, an N-terminal FLAG pLX304 plasmid was digested with AfeI and SrfI restriction enzymes. The cut plasmid was then assembled with a DNA fragment synthesized by Twist Bioscience using the NEBuilder HiFi DNA Assembly Kit to generate a C-terminal FLAG-tagged pLX304 plasmid. Gateway LR Clonase II Enzyme Mix was used to generate the pLX304 C-terminal FLAG ZNF207 construct by recombining the siRNA-resistant ZNF207 sequence from the pDONR223 vector into the pLX304 C-terminal FLAG backbone (Addgene #231939 – pLenti-ZNF207-3xFLAG). All cloned constructs were confirmed by long read nanopore sequencing.

### ZNF207 rescue experiments

For transient transfection experiments, 700,000 HEK293T cells expressing the LMNA mutant minigene were plated for non-targeting control siRNA (siControl) treatment, and 1 million cells were plated for siRNA targeting ZNF207 (siZNF207). Sixteen hours post-siRNA transfection (25 nM siRNA) using Lipofectamine RNAiMAX (ThermoFisher Scientific #13778150), cells were transfected with plasmids using X-tremeGENE9 (Sigma #6365779001). Cells were collected 48 hours after plasmid transfection for further analysis.

For experiments with stable ZNF207-expressing HEK293T Flp-In cells, 700,000 cells were plated for non-targeting control siRNA (siControl), and 1 million cells were plated for siZNF207. Six hours after siRNA transfection, doxycycline was added as previously described to induce ZNF207 expression. Cells were collected and frozen 48 hours post-siRNA transfection for downstream applications.

### miniTurboID mass spectrometry

Eight million HEK293 Flp-In cells were plated for each cell line (miniTurbo-Empty, miniTurbo-EGFP, or ZNF207-miniTurbo) in 15 cm cell culture dishes. After 48 hours, doxycycline was added to the media at a final concentration of 0.0005 µg/mL for miniTurbo-Empty and miniTurbo-EGFP (Addgene #209059), and 0.5 µg/mL for ZNF207-miniTurbo (Addgene #231938). Twenty hours after doxycycline treatment, D-Biotin (ThermoFisher Scientific #B20656) was added to the media at a final concentration of 50 µM. After an additional 4 hours of incubation, approximately 30 million cells were harvested and frozen for BioID analysis, while ∼50,000 cells were collected for immunoblotting.

HEK293 Flp-In cells lines were lysed in 1 mL of 0.5% NP40 lysis buffer (150 mM NaCl, 50 mM HEPES pH 7.2, 0.5% NP40 [Sigma-Aldrich #I3021]) containing protease inhibitors (Roche cOmplete #11836145001; 1 tablet per 50 mL Lysis Buffer). Cell pellets were resuspended by gentle pipetting until homogenous, followed by pulse sonication to shear DNA. Lysates were clarified by centrifugation at 18,500

× g for 10 minutes at 4°C. Protein concentration was determined using a BCA assay, and 1 mg of total protein per sample was adjusted to 1 mL with cold 0.5% NP40 lysis buffer. Protein lysates were incubated with 60 µL of a 50% slurry of Streptavidin Sepharose Beads (ThermoFisher Scientific #20353) overnight at 4°C with rotation. Beads were washed sequentially: three times with 1 mL cold 0.5% NP40 lysis buffer, three times with ice-cold 1× TBS, and once with 50 mM HEPES (pH 8.0). Washed beads were resuspended in 50 mM HEPES (pH 8.0) and stored at −80°C.

Beads were thawed on ice, heated to 95°C for 4 minutes, and cooled to room temperature. Each sample was digested with 2 µg trypsin (Promega #V5111) overnight at 37°C with gentle rocking. Peptides were cleaned using EasyPep™ MS Sample Prep Kits (ThermoFisher Scientific #A57864) according to the manufactures instructions and dried in a speedvac.

Dried peptides were resuspended in 15 µL of 0.1% TFA and analyzed on an EASY-nLC 1200 system coupled to a Q Exactive HF mass spectrometer (ThermoFisher Scientific) with an EasySpray ion source. Peptides were loaded onto an Acclaim PepMap 100 C18 trap column (75 µm × 2 cm; ThermoFisher Scientific) and separated on a PepMap RSLC C18 analytical column (75 µm × 25 cm). Elution was performed using a two-step gradient: 5–27% acetonitrile with 0.1% formic acid over 60 minutes, followed by 27–40% acetonitrile with 0.1% formic acid over 45 minutes, at a flow rate of 300 nL/min. MS1 scans were acquired at 60,000 resolution over a mass range of 380–1580 m/z, with a maximum injection time of 120 ms and an AGC target of 3 × 10⁶. MS2 scans were performed at 15,000 resolution, using a normalized collision energy of 27, maximum injection time of 50 ms, and an AGC target of 2 × 10⁵.

Raw MS data were processed using Proteome Discoverer 2.4 (ThermoFisher Scientific) with the Sequest search engine. Data were searched against the UniProt human database using fully tryptic peptides, allowing up to 2 missed cleavages. Peptide length was restricted to 6–40 amino acids, with precursor and fragment mass tolerances of 10 ppm and 0.02 Da, respectively. FDR was set to ≤ 1% using Percolator.

Protein intensities were normalized to the sample with the highest total intensity. For each protein, median intensities within experimental groups were compared to control groups (e.g., empty vector or EGFP) using a one-tailed t-test assuming equal variance.

Proximal interactors consistently identified across all three experimental replicates were selected for network visualization (Figure 5B) and gene ontology (GO) analysis (Figures 5C and S7B). For GO analysis, enrichment of biological processes and CORUM protein complexes was performed using g:Profiler.^98^ The background set included all proteins detected in the mass spectrometry experiments (Table S7), including those from control samples. Proteins included in the Figure 5B network were restricted to those with a median peptide count of five or higher. Network visualization in Figure 5B was created using the Cytoscape STRING application^99,100^, for importing STRING networks.

### qPCR validations

For most of qPCR validation, 40 ng of RNA was used in the SensiFAST SYBR No-ROX One-Step Kit (Thomas Scientific #C755H72) according to the manufacturer’s protocol. For SMG1 NMD inhibitor treatment experiments (qPCR targeting. LUC7L, LUC7L2, SNRNP70 and RBM3), cDNAs were synthesized using 1ug of RNAs and Maxima H Minus First Strand cDNA Synthesis Kit (ThermoFisher Scientific #K1651) followed by 1:25 dilution. cDNA equivalent to 4 ng of RNA (2 uL after dilution) was used in the SensiFAST SYBR No-ROX Kit (Thomas Scientific #C755J02). Two to three technical replicates were performed for each sample to ensure accuracy, and outlier values were excluded from the analysis to obtain reliable results. The primers are listed in Table S2.

### Immunofluorescence staining

Cells were grown on 384-well (Perkin Elmer, #6057300) microplates and fixed for 15 min with 4% paraformaldehyde (PFA, Electron Microscopy Sciences). Fixed cells were then washed one time with PBS, permeabilized for 10 min (PBS/0.5% Triton X-100) and washed again one time with PBS. Next, cells were incubated with primary antibodies diluted in blocking buffer (PBS, 0.05% Tween20, 5% bovine serum albumin) for 1.5 hour. After two consecutive washes with PBS, cells were incubated with secondary antibodies diluted in blocking buffer for 1 hour and washed once with PBS. Cells were then counterstained with DAPI (Biotium, #40043) for 30 min and washed once with PBS. After washing, cells could be stored for extended periods of time in PBS at 4°C. All steps for IF staining were performed at ambient temperature. Primary antibodies used for immunofluorescence were: α-LAP2alpha rabbit polyclonal (1:400, Abcam, ab5162), α-Lamin B1 rabbit polyclonal (1:200, Abcam, #ab16048), α-Histone H3 (tri methyl K27) rabbit monoclonal (1:400, Abcam, #ab192985) and α-acetyl-Histone H4 (Ac-Lys16) rabbit recombinant monoclonal (1:400, Abcam, #ab109463). The secondary antibody used for immunofluorescence detection was Alexa Fluor Goat-anti-Rabbit 594 (1:400, Invitrogen, #A11012).

### High-throughput image acquisition, quantification and statistical analysis

Cells were imaged in two channels (405 and 561 nm excitation lasers) in an automated fashion using a dual spinning disk high-throughput confocal microscope (Yokogawa CV7000S) with a 405/488/561/640 nm excitation dichroic mirror, a 40 air objective lens (NA 0.95), a 568 nm excitation dichroic mirror, and two 16-bit sCMOS cameras (Andor, 2560 × 2160 pixels) with binning set to 2 (Pixel size: 0.325 nm for 40X and 0.216 nm for 60X nm). Emission bandpass filters were used for each channel: 445/45 and 600/37 nm. Sixteen randomly selected fields of view were imaged per well in a single optimal focal plane. Images were corrected on the fly using a geometric correction for camera background, illumination field (vignetting), camera alignment, and chromatic aberrations. Corrected images were saved and stored as 16-bit TIFF files.

Captured images from high-throughput imaging experiments were analyzed using SiMA 1.3 (Revvity Signals). Briefly, nuclei regions of interest (ROI) were segmented using the DAPI (405 nm) channel and nucleus ROIs adjacent to the image edges were excluded from subsequent image analysis steps. The mean fluorescence intensity for the nucleus ROI was measured in the 561 nm channel. SiMA results were exported as comma-separated text files and analyzed using R. 600-1000 cells were analyzed per condition.

High-throughput imaging experiments were performed on three technical replicates (wells) and averaged to obtain the measurement for that experiment. The experiments were repeated independently three times (n = 3), and the values for each experiment were averaged. Single cell data analysis was performed using R software and the data are shown as the average mean values of 3 different experiments +/- standard deviation (SD). Statistical analysis for the average mean values of single cell data was performed by an ANOVA one-way test, followed by a post-hoc Dunnett’s test using the cell line as the predictor variable and fixing wild-type (WT) fibroblasts treated with control siRNas (siCTL) as the negative control for the WT cells treated with siZNF207 and HGPS fibroblasts treated with siCTL as the negative control for HGPS siZNF207. Corresponding p-value is shown in every figure.

### eCLIP sequencing and analysis

Twelve million cells were plated per cell line (FLAG-EGFP, FLAG-ZNF207 N-terminus, or ZNF207-FLAG C-terminus) in 15 cm cell culture dishes. After 24 hours, doxycycline was added to the media at a final concentration of 0.5 µg/mL to induce protein expression. Cells were washed twice with PBS and resuspended in 3 mL of PBS. UV-crosslinking was performed at 254 nm with an energy setting of 400 mJ/cm². Following crosslinking, cells were counted, and approximately 21 million viable cells were pelleted, frozen, and sent to EclipsBio for further assays, library generation and sequencing. eCLIP was performed based on the single-end seCLIP protocol^101^ with specific modifications. Samples were lysed using 1 mL of eCLIP lysis buffer containing 6 µL of Proteinase Inhibitor Cocktail and 20 µL of Murine RNase Inhibitor. Lysates underwent two rounds of sonication (4 minutes total) with 30-second ON/OFF cycles at 75% amplitude. For immunoprecipitation, a pre-validated anti-FLAG antibody was pre-coupled to Anti-Rabbit IgG Dynabeads (ThermoFisher Scientific #11202D). The antibody-bead complex was added to the lysate and incubated overnight at 4°C. Before immunoprecipitation, 2% of the lysate was set aside as the paired input control. The remaining sample was subjected to magnetic separation and washed thoroughly with eCLIP high-stringency wash buffers. Immunoprecipitation and input samples were excised from the membrane at a size corresponding to ∼75 kDa above the expected protein band. Subsequent steps, including RNA adapter ligation, IP-Western blot validation, reverse transcription, DNA adapter ligation, and PCR amplification, were performed as previously described in the seCLIP protocol.^101^

We used FastQC and MultiQC^102^ for read quality assessments at various stages of the eCLIP-Seq analytical pipeline. UMI (Unique Molecular Identifier) barcodes were identified and annotated using UMI-tools^103^. Cutadapt^104^ was used for the read trimming. Trimmed reads were first aligned to repetitive elements of human genome (sourced from RepBase, GIRI (Genetic Information Research Institute) (https://www.girinst.org/server/RepBase/)) so that they can be later separated to increase accuracy of the eCLIP-Seq peak identification in the subsequent steps. The unaligned reads from this step were used to perform the main alignment with human genome (hg38) using gencode v44 primary assembly genome fasta and GTF files. All the mapping steps were performed with STAR^105^. The mapped reads were deduplicated using UMI-tools, which considers UMI annotations from the preprocessing step and alignment positions to remove duplicate reads from the samples. Thus, generated final alignments were considered for peak calling and generation of metagene coverage plots. We used SAMtools^106^ to merge, filter, and index Binary Alignment Files (BAM) and BEDTools^107^ to manage BED files. We used the HTSeq-CLIP tool (v2.19)^108^ with default parameters for peak calling, followed by statistical analysis using the DEWSeq tool (v1.18)^109^. Regions of enrichment (enrichedWindowRegions) were identified based on a log2 fold change (L2FC) ≥ 0.5 and an adjusted p-value (padj) ≤ 0.05. These regions were further analyzed to determine the RNA biotypes, genomic locations, and gene categories associated with the bound targets.

DeepTools^110^ was used to generate ZNF207 metagene coverage plots using a subtraction-based IP_vs_Input normalization method. We determined ZNF207 regulated cassette exons by assessing siZNF207 responsive splicing events that are rescued by ZNF207_FL overexpression from RNA-Seq data analysis. We selected all the alternative exons that are not regulated by ZNF207 as a control for comparison with following criteria: (1) 90 ≥ PSI ≥ 10 for siZNF207 knock-down condition (2) |dPSI| < 5 and probability < 0.95 for siZNF207 knock-down condition. We used cassette exons of size ≥ 6bp of protein coding genes with minimum FPKM expression value of 0.1 to perform comparative metagene enrichment analysis for ZNF207 eCLIP-Seq data.

### Design of genome-wide knock-out library

The CHyMErA hybrid guide RNA (hgRNA) gene knockout library was constructed to enable dual-targeting of individual genes and combinatorial targeting of paralog gene pairs using Cas9 and Cas12a enzymes. The process began with the curation of protein-coding genes from the Gencode Comprehensive Annotation v43. Genes and transcripts were filtered to ensure high-quality annotations. Only genes labeled as protein-coding were retained, while polymorphic pseudogenes and transcript isoforms with fewer than two untranslated regions (UTRs) were excluded, except when all transcripts for a gene had fewer than two UTRs. Transcripts with a transcript support level of 3 or higher were removed unless their exclusion would eliminate a gene entirely. Additionally, fusion genes and annotations within the mitochondrial genome were excluded.

Single guide RNAs (sgRNAs) targeting human protein-coding genes were then selected from a database of all possible Cas9 and Cas12a sgRNAs, based off the presence of “NGG” and “TTTV” PAMs, respectively. Only sgRNAs with upper and lower cut sites within the coding sequence (CDS) of a gene were considered. These sgRNAs were required to target a single site in the genome (Hamming.0 = 1) without any genomic sites within one mismatch (Hamming.1 = 0), and Cas12a spacers with no more than 2 sites in the genome with 2 mismatches (Hamming.2 <= 2). Guide RNAs with off-target GuideScan scores^111,112^ exceeding 0.16 for both enzymes (Cas9 and Cas12a) were retained. On-target efficiency was assessed using Cas9 Rule Set 2 scores^113^ and Cas12a-specific CHyMErA_Net scores^33^, both of which were required to exceed 0.1. Additional filtering was applied to eliminate guide RNAs containing or creating BveI and Eco47III restriction sites, which are critical for downstream library construction and reporter cloning.

Each Cas9 and Csa12a sgRNA for every unique Ensembl GeneID was ranked based on its quality, assessed by transcript coverage (Transcript_Percentage_Targeted), cut site location (N..C), spacer overlap, and On-Target Score. The percentage of transcript isoforms targeted by each guide was evaluated, with optimal percentage of transcripts targeted being 100%, cut sites located at least 10% downstream of the start codon (N-terminal) and no more than 50% upstream of the stop codon (C-terminal) of every transcript, and new spacers with no shared genomic location with other chosen guides. Guides were iteratively filtered, with quality thresholds reducing by fifths (Transcript_Percentage_Targeted from 1 -> 0.8, cut sites within 8% downstream of the start codon and 40% upstream of the stop codon, and minimal spacer overlaps from 0->4) with guides passing each quality threshold ordered by On-Target Score until all guides have been ranked. Based on this assessment, sgRNAs assigned quality ranks from 1 (highest) to 6 (lowest) based off the Round of quality each guide passed. The top 25 sgRNAs for each gene were selected for both Cas9 (a1-a25) and Cas12a (b1-b25).

To construct hybrid guides RNAs (hgRNAs), the top 25 sgRNAs for each enzyme were paired combinatorially. hgRNAs were evaluated based on the same criteria as sgRNAs with the added quality criteria of minimal genomic distance between Cas9 and Cas12a cut sites. hgRNA pairs targeting all transcript isoforms in the upstream-middle of the CDS with no spacer overlap and an optimal genomic distance of at least 150 base pairs were prioritized. If these criteria were not met, iterative quality thresholds were applied to relax the selection criteria, with lower thresholds reducing the stringency of Percentage Transcript_Targeted, transcript CDS cut site, spacer overlap and hgRNA cut genomic distance requirements by one fifth until 4 hgRNAs were found. Each hgRNA was labeled according to the quality rank of its Cas9 and Cas12a components (e.g., a6-b3, where a6 represents the 6th-ranked Cas9 guide and b3 the 3rd-ranked Cas12a guide). The selection process ensured that the best pairs were prioritized based on a balance of these criteria.

To ensure diversity among hgRNAs, selected pairs were shuffled and reanalyzed to maximize the distribution of quality sgRNAs while maintaining overall hgRNA quality. For example, if the initial selection yielded pairs such as a1-b1, a2-b2, a3-b3, and a4-b4, a subsequent reshuffling could generate pairs such as a1-b4, a2-b3, a3-b2, and a4-b1, provided the reshuffled pairs maintain hgRNA quality thresholds. In cases with limited sgRNA availability, specific rules guided the selection process. When only three Cas9 and three Cas12a sgRNAs were available, only three hgRNAs were selected. For three Cas9 and two Cas12a sgRNAs, two hgRNAs were chosen along with a Cas9-only guide paired with a Cas12a intergenic guide, and vice versa for two Cas9 and three Cas12a sgRNAs. If three Cas9 and one Cas12a sgRNAs were available, one hgRNA was formed along with two Cas9-only guides paired with two Cas12a intergenic guides and vice versa for one Cas9 and three Cas12a sgRNAs. When two Cas9 and two Cas12a sgRNAs were available, two hgRNAs were chosen alongside the same Cas9 sgRNAs paired with Cas12a intergenic guides. For two Cas9 and one Cas12a sgRNA, one hgRNA was formed alongside two Cas9 sgRNAs paired with 2 Cas12a intergenic guides and one Cas12a sgRNA paired with one Cas9 intergenic guide and vice versa for 1 Cas9 and 2 Cas12a sgRNAs. If only one Cas9 and one Cas12a guide were available, three guide combinations were formed: one hgRNA, one Cas9-only guide, and one Cas12a-only guide. For cases where two or more Cas9 sgRNAs were available but no Cas12a sgRNAs, intergenic Cas12a guides were added, and the reverse was applied for cases where two or more Cas12a sgRNAs were available but no Cas9 sgRNAs. No hgRNAs were selected for genes where only a single Cas9 or Cas12a sgRNA was available. This approach allowed flexibility while maintaining the library’s robustness and comprehensiveness.

The targeted paralogs were chosen based on definitions established in previous studies.^33,114–116^ Additionally, 624 hand-picked gene pairs, primarily involved in splicing-related processes, were included to enhance the focus on splicing regulation. The construction of the paralog knockout hybrid guide RNA (hgRNA) library utilized computational criteria to identify optimal Cas9 and Cas12a sgRNAs for targeting paralog gene pairs. To begin, information such as gene IDs and genomic coordinates was extracted using the BioMart R package with the hsapiens_gene_ensembl dataset (version 104). Key attributes, including chromosome name, start position, HGNC symbol, Ensembl gene ID, description, external gene source, and gene biotype, were retrieved using the getBM function with filters “hgnc_symbol” (or “external_synonym”), “chromosome_name,” and “biotype.” Paralog pairs were then sorted based on their library of origin, which included both previously generated paralog combinations and handpicked paralog pairs.

For each paralog pair, the selection process aimed to identify the best Cas9 and Cas12a sgRNAs for targeting Paralog A and Paralog B based on the same criteria used to rank sgRNAs for the dual knockout hgRNA library detailed above. When at least two Cas9 and Cas12a sgRNAs were available for both paralogs, hgRNA combinations were extracted, targeting each paralog twice for a total of four hybrid guides. Two Cas9 (A1- and A2-) and two Cas12a (-A1 and -A2) guides targeting gene A and two Cas9 (B1- and B2-) and Cas12a (-B1 and -B2) guides targeting gene B will make hybrid guide RNA combinations (A1-B1, A2-B2, B1-A1, and B2-A2) with Cas9 guides in the first position and Cas12a guides in the second position). If no Cas9 guides targeted Paralog A but Cas9 guides were available for Paralog B, Cas12a guides targeting Paralog A were used to form hybrid combinations (e.g. B1-A1, B2-A2, B3-A3, B4-A4). By default, if only Cas9 guides or Cas12a are available to target gene A and B, no hgRNAs will be chosen.

The full library was further supplemented with 331 fully intergenic hgRNAs and 99 non-targeting hgRNAs to serve as controls. A pool of 95,893 oligos, each 187 nucleotides in length, was designed to include a 20-nt Cas9 guide sequence, followed by the Cas9 tracrRNA, the AsCas12a direct repeat (DR), and a 23-nt Cas12a guide sequence (Table S1). The oligos were synthesized by TWIST Biosciences and cloned into the CRASP-Seq vector as described above.

### Design base editor library

We designed an sgRNA tiling library to target the coding sequences of all isoforms for 39 selected genes. Exon coordinates were derived from the GRCh38 assembly using Gencode version 46 (GFF3 file). To ensure compatibility with downstream cloning steps and maintain sgRNA efficiency, guides containing homopolymer stretches (AAAAAA, TTTTT, CCCCCC, GGGGGG) or restriction motifs (e.g., GATATC, GCAGGT, ACCTGC) were excluded. Additional filters removed sgRNAs with sequences starting with CAGGT or ATATC, or ending with GCAG, to prevent interference with EcoRV and BfuAI/BveI recognition sites. To enable base editing and ensure effective targeting, only sgRNAs overlapping at least 1 nucleotide of an exon and containing an A or C residue within positions 1–12 of the spacer sequence were retained. Both plus- and minus-strand sgRNAs were included. Off-target effects were minimized by applying GuideScan thresholds: an off-target score > 0.16 and a Hamming distance ≤ 2, as established previously.^112^ In total, the library consists of 6,984 sgRNAs targeting the 39 genes, supplemented by 1,520 intergenic and 92 non-targeting control sgRNAs, that were selected from our previous Cas9 sgRNA libraries (Table S8).^11^ The library oligo pool consists of 144-nucleotide sequences, encompassing the Cas9 protospacer region followed by a modified tracrRNA sequence.^117^ These sequences are flanked by stuffer regions containing BveI restriction sites, enabling efficient cloning into the digested CRASP-Seq vector.

### CRASP-Seq screening protocol

The generated lentiviral library was used to infect HAP1, RPE1, or HepG2 cells expressing *Sp*Cas9 and opCas12a nucleases^11,37^ at a multiplicity of infection (MOI) of 0.2, ensuring a coverage of at least 250 cells per hgRNA. After 24 hours, selection was initiated with 2 µg/mL puromycin. Forty-eight hours post-selection (T0), the surviving cells were pooled, plated, and treated with doxycycline to induce reporter expression. After 24 hours (T1), cells were harvested, aliquoted into 10 million-cell samples, and prepared for further analysis. For the SRSF7 poison exon screen, 100 µg/mL cycloheximide was added 4 hours before harvesting to inhibit NMD.

Total RNA was extracted using the RNeasy Plus Mini Kit (QIAGEN #74136) according to the manufacturer’s instructions. mRNA was then isolated using oligo(dT) Dynabeads (ThermoFisher Scientific #61002), followed by cDNA synthesis using the Maxima H Minus First Strand cDNA Synthesis Kit (ThermoFisher Scientific #K1651) with a custom primer (**Table S2**). The resulting cDNA was purified using the QIAquick PCR Purification Kit (QIAGEN #28104). PCR was optimized and performed using the NEBNext Ultra II Q5 Master Mix (New England Biolabs #M0544X) under the following conditions: 98°C for 30 seconds (initial denaturation), followed by 14-17 cycles of 98°C for 10 seconds, 60°C for 20 seconds, and 72°C for 30 seconds, with a final extension at 72°C for 1 minute. Illumina barcoded forward and reporter-specific reverse primer pairs were utilized (**Table S2**). The PCR products were visualized on a 1-1.5% agarose gel stained with SYBR Safe (ThermoFisher Scientific #S33102). The desired bands were excised and purified using a gel extraction kit (ThermoFisher Scientific #K0691). The extracted libraries were quantified using the Qubit dsDNA HS Assay (ThermoFisher Scientific #Q32851) and Agilent TapeStation (Agilent #5067-5582 and #5067-5583). The validated sequencing libraries were pooled and sequenced on the NovaSeq 6000 platform using 300-cycle kits, with a 20-25% PhiX spike-in. The sequencing strategy most commonly employed was: Read 1: 210, Index Read 1: 8, Index Read 2: 8, Read 2: 104.

### Analysis of CRASP-Seq data

The CRASP-Seq data were analyzed using a custom bioinformatics pipeline consisting of several sequential steps. Briefly, the pipeline involves quantifying the Percent Spliced In (PSI) index for the minigene reporter associated with each hgRNA expressed in the cells. UMIs introduced during cDNA synthesis is used to remove PCR duplicates, while CBCs are applied to randomly generate two replicate samples. By applying a 10% false discovery rate (FDR) threshold, we establish a PSI cutoff to determine whether individual hgRNAs positively or negatively regulate a minigene reporter. Positive and negative regulators are defined as those with at least two hgRNAs detected to affect reporter splicing. Only regulators identified in both experimental replicates were considered significant hits.

More specifically, we first we performed mapping and guide annotation by retrieving each component of the CRISPR-Cas system and splicing reporter cassette, including the Cas9 guide, Cas12a guide, CBC, and UMIs from Read 1, as well as minigene splicing event monitoring from Read 2. These components were extracted from sequencing reads using plasmid backbone-adjacent anchor sequences and mapped with the aligner tool STAR^105^ to annotate the guide sequences based on the genome-wide CRISPR library. Alignment files were then managed using SAMtools^106^ to facilitate downstream applications. PCR-mediated sequence deduplication was addressed by monitoring CBCs and UMIs, allowing a hamming distance of 1 for each sequence element to remove duplicates. Only fully annotated and unique reads were retained for quantifying splicing regulation and linking splicing outcomes to specific genetic perturbations. To assess splicing outcomes, Percent Splicing In (PSI) and ΔPSI scores were calculated as follows:

- PSI (Percent Splicing In) = (IE/(IE+EE)) * 100, where IE represents inclusion event counts and EE represents exclusion event counts.
- ΔPSI was defined as PSI_guide - PSI_intergenic, where PSI_guide represents the PSI value for a particular guide, and PSI_intergenic represents the cumulative PSI value of all intergenic guides in the sequencing library.

To identify individual hgRNAs affecting the splicing reporter, we calculated an empirical 10% FDR by analyzing intergenic hgRNAs and setting PSI cutoffs to identify significant hgRNAs as those at the 5% extremes of the intergenic distribution. At the gene level, we further required that the ΔPSI for each gene (calculated across all guides targeting the same gene) met this cutoff, with more than half of the guide RNAs for the gene exceeding this threshold (e.g., 3 out of 4 guides or 2 out of 3 guides). Genes with at least two guides identified as hits were assigned as splicing regulator hits. Additionally, genes were considered hits only if they met these criteria in both experimental replicates, as determined by splitting data based on the CBC barcode, to ensure robust hit calling.

The base editor screen was analyzed by estimating ΔPSI scores for all sgRNAs and genes (by merging all the sgRNAs targeting the same gene). To plot the sequence-level tiling profile of ZNF207, the protein region corresponding to the guide RNAs were retrieved using Ensembldb package (version 2.30.0). We extracted genome coordinates of ZNF207 (Transcript ENST00000394670.9) from Ensembl 113 using AnnotationHub and then used the genomeToProtein function to map the guide RNA (lying completely within the exons) to the positions of protein sequence. For the genes with forward orientation, we defined the positions by considering the start index of mapped positions while for the reverse orientation last index was taken. The ΔPSI score is the average of the ΔPSI adenine and cytosine base editor screens calculated for each guide RNA as explained above. For gene-level analysis, the average adenine and cytosine ΔPSI scores for each gene were calculated as described above. These scores were then used to plot bar graphs, with genes arranged in ascending order of their ΔPSI scores.

### Gene expression and splicing analysis from RNA-sequencing data

The raw FASTQ reads were trimmed with Cutadapt^104^ tool and mapped to human genome (hg38) with STAR aligner^105^. We used gencode v44 reference files (primary assembly genome fasta and primary assembly GTF (Gene Transcript Features) for mapping and transcript annotation steps. STAR generated gene count files were used for differential gene expression analysis with DESeq2^118^. We used SAMtools^106^ to manage alignment files for various steps such as merging and indexing Binary Alignment Files (BAM). Following statistical parameters were calculated to study differential gene expression either through custom scripts or through existing R packages such as DESeq2: (1) FPKM (Fragments per Kilo-base of Gene per Million Reads): ((Paired Reads/library size) *1000000)/ gene size) *1000 (2) L2FC (Log2 Fold Change): - log2 (Test Counts / Control Counts). ZNF207-regulated genes were defined as those exhibiting an absolute log2 fold change (LFC) difference of ≥ 0.35 between siControl and siZNF207 conditions, with an adjusted p-value < 0.05. Additionally, these genes were required to show similar differences between siZNF207-treated cells expressing the empty vector and those expressing C-terminally tagged ZNF207. Only genes with FPKM values > 0.1 were included in the analysis. For GO analysis (Figure S6E), enrichment of biological processes was performed using g:Profiler.^98^ The background set comprised all protein-coding genes with FPKM values greater than 0.1.

Whippet^119^ tool was used to perform differential splicing analysis with trimmed FASTQ files and STAR generated alignment files to identify known and novel splicing events in any test_vs_control comparison using above mentioned reference files. ZNF207-regulated splicing events were identified as those displaying an absolute splicing change (ΔPSI) of ≥10 between siControl and siZNF207 conditions, with a probability score of ≥0.95. Furthermore, these events had to exhibit consistent differences between siZNF207-treated cells expressing an empty vector and those expressing C-terminally tagged ZNF207. Only genes with FPKM values exceeding 0.1 were included in the analysis.

### Gene ontology enrichment analysis

Gene ontology (GO) term enrichment analysis of significant hits from the CRASP-Seq screens was conducted using g:Profiler^98^ with the protein-coding gene set targeted by the CRISPR library serving as the background reference (Table S1). Network views (Figures 1D and S4F) were visualized in Cytoscape^99,120^ using the Enrichment Map plugin, which organizes categories based on shared gene overlaps to highlight related functional clusters. For the annotation of heatmap subclusters in Figure 2E, we conducted Gene Set Enrichment Analysis (GSEA) using the enricher function from the clusterProfiler^121^ tool with default parameters.

## BIBLIOGRAPHY

1. Kastner, B., Will, C.L., Stark, H., and Lührmann, R. (2019). Structural Insights into Nuclear pre-mRNA Splicing in Higher Eukaryotes. Cold Spring Harb Perspect Biol 11. 10.1101/cshperspect.a032417.

2. Wan, R., Bai, R., Zhan, X., and Shi, Y. (2020). How Is Precursor Messenger RNA Spliced by the Spliceosome? Annu Rev Biochem 89, 333–358. 10.1146/annurev-biochem-013118-111024.

3. Tholen, J., and Galej, W.P. (2022). Structural studies of the spliceosome: Bridging the gaps. Curr Opin Struct Biol 77, 102461. 10.1016/j.sbi.2022.102461.

4. Martínez-Lumbreras, S., Morguet, C., and Sattler, M. (2024). Dynamic interactions drive early spliceosome assembly. Curr Opin Struct Biol 88, 102907. 10.1016/j.sbi.2024.102907.

5. Ule, J., and Blencowe, B.J. (2019). Alternative Splicing Regulatory Networks: Functions, Mechanisms, and Evolution. Mol Cell 76, 329–345. 10.1016/j.molcel.2019.09.017.

6. Marasco, L.E., and Kornblihtt, A.R. (2023). The physiology of alternative splicing. Nat Rev Mol Cell Biol 24, 242–254. 10.1038/s41580-022-00545-z.

7. Rogalska, M.E., Vivori, C., and Valcárcel, J. (2023). Regulation of pre-mRNA splicing: roles in physiology and disease, and therapeutic prospects. Nat Rev Genet 24, 251–269. 10.1038/s41576-022-00556-8.

8. Baralle, F.E., and Giudice, J. (2017). Alternative splicing as a regulator of development and tissue identity. Nat Rev Mol Cell Biol 18, 437–451. 10.1038/nrm.2017.27.

9. Pan, Q., Shai, O., Lee, L.J., Frey, B.J., and Blencowe, B.J. (2008). Deep surveying of alternative splicing complexity in the human transcriptome by high-throughput sequencing. Nature genetics 40, 1413–1415. 10.1038/ng.259.

10. Wang, E.T., Sandberg, R., Luo, S., Khrebtukova, I., Zhang, L., Mayr, C., Kingsmore, S.F., Schroth, G.P., and Burge, C.B. (2008). Alternative isoform regulation in human tissue transcriptomes. Nature 456, 470–476. 10.1038/nature07509.

11. Xiao, M.-S., Damodaran, A.P., Kumari, B., Dickson, E., Xing, K., On, T.A., Parab, N., King, H.E., Perez, A.R., Guiblet, W.M., et al. (2024). Genome-scale exon perturbation screens uncover exons critical for cell fitness. Molecular Cell 84, 2553–2572.e2519. 10.1016/j.molcel.2024.05.024.

12. Fu, X.D., and Ares, M., Jr. (2014). Context-dependent control of alternative splicing by RNA-binding proteins. Nat Rev Genet 15, 689–701. 10.1038/nrg3778.

13. Park, J.W., Parisky, K., Celotto, A.M., Reenan, R.A., and Graveley, B.R. (2004). Identification of alternative splicing regulators by RNA interference in Drosophila. Proc Natl Acad Sci U S A 101, 15974–15979. 10.1073/pnas.0407004101.

14. Brooks, A.N., Duff, M.O., May, G., Yang, L., Bolisetty, M., Landolin, J., Wan, K., Sandler, J., Booth, B.W., Celniker, S.E., et al. (2015). Regulation of alternative splicing in Drosophila by 56 RNA binding proteins. Genome Res 25, 1771–1780. 10.1101/gr.192518.115.

15. Tejedor, J.R., Papasaikas, P., and Valcárcel, J. (2015). Genome-wide identification of Fas/CD95 alternative splicing regulators reveals links with iron homeostasis. Mol Cell 57, 23–38. 10.1016/j.molcel.2014.10.029.

16. Papasaikas, P., Tejedor, J.R., Vigevani, L., and Valcárcel, J. (2015). Functional splicing network reveals extensive regulatory potential of the core spliceosomal machinery. Mol Cell 57, 7–22. 10.1016/j.molcel.2014.10.030.

17. Rogalska, M.E., Mancini, E., Bonnal, S., Gohr, A., Dunyak, B.M., Arecco, N., Smith, P.G., Vaillancourt, F.H., and Valcárcel, J. (2024). Transcriptome-wide splicing network reveals specialized regulatory functions of the core spliceosome. Science 386, 551–560. 10.1126/science.adn8105.

18. Manning, K.S., and Cooper, T.A. (2017). The roles of RNA processing in translating genotype to phenotype. Nat Rev Mol Cell Biol 18, 102–114. 10.1038/nrm.2016.139.

19. Scotti, M.M., and Swanson, M.S. (2016). RNA mis-splicing in disease. Nature reviews. Genetics 17, 19–32. 10.1038/nrg.2015.3.

20. Gordon, L.B., Brown, W.T., and Collins, F.S. (1993). Hutchinson-Gilford Progeria Syndrome. In GeneReviews(®), M.P. Adam, J. Feldman, G.M. Mirzaa, R.A. Pagon, S.E. Wallace, L.J.H. Bean, K.W. Gripp, and A. Amemiya, eds. (University of Washington, Seattle Copyright © 1993-2024, University of Washington, Seattle. GeneReviews is a registered trademark of the University of Washington, Seattle. All rights reserved.).

21. Gordon, L.B., Rothman, F.G., López-Otín, C., and Misteli, T. (2014). Progeria: a paradigm for translational medicine. Cell 156, 400–407. 10.1016/j.cell.2013.12.028.

22. Bradley, R.K., and Anczuków, O. (2023). RNA splicing dysregulation and the hallmarks of cancer. Nat Rev Cancer. 10.1038/s41568-022-00541-7.

23. Bonnal, S.C., López-Oreja, I., and Valcárcel, J. (2020). Roles and mechanisms of alternative splicing in cancer - implications for care. Nat Rev Clin Oncol 17, 457–474. 10.1038/s41571-020-0350-x.

24. Kahles, A., Lehmann, K.V., Toussaint, N.C., Hüser, M., Stark, S.G., Sachsenberg, T., Stegle, O., Kohlbacher, O., Sander, C., Caesar-Johnson, S.J., et al. (2018). Comprehensive Analysis of Alternative Splicing Across Tumors from 8,705 Patients. Cancer Cell 34, 211–224.e216. 10.1016/j.ccell.2018.07.001.

25. Calabrese, C., Davidson, N.R., Demircioğlu, D., Fonseca, N.A., He, Y., Kahles, A., Lehmann, K.V., Liu, F., Shiraishi, Y., Soulette, C.M., et al. (2020). Genomic basis for RNA alterations in cancer. Nature 578, 129–136. 10.1038/s41586-020-1970-0.

26. Yoshimi, A., Lin, K.T., Wiseman, D.H., Rahman, M.A., Pastore, A., Wang, B., Lee, S.C., Micol, J.B., Zhang, X.J., de Botton, S., et al. (2019). Coordinated alterations in RNA splicing and epigenetic regulation drive leukaemogenesis. Nature 574, 273–277. 10.1038/s41586-019-1618-0.

27. Jung, H., Lee, D., Lee, J., Park, D., Kim, Y.J., Park, W.Y., Hong, D., Park, P.J., and Lee, E. (2015). Intron retention is a widespread mechanism of tumor-suppressor inactivation. Nat Genet 47, 1242–1248. 10.1038/ng.3414.

28. David, C.J., Chen, M., Assanah, M., Canoll, P., and Manley, J.L. (2010). HnRNP proteins controlled by c-Myc deregulate pyruvate kinase mRNA splicing in cancer. Nature 463, 364–368. 10.1038/nature08697.

29. Wang, Z., Jeon, H.Y., Rigo, F., Bennett, C.F., and Krainer, A.R. (2012). Manipulation of PK-M mutually exclusive alternative splicing by antisense oligonucleotides. Open Biol 2, 120133. 10.1098/rsob.120133.

30. Paronetto, M.P., Bernardis, I., Volpe, E., Bechara, E., Sebestyén, E., Eyras, E., and Valcárcel, J. (2014). Regulation of FAS exon definition and apoptosis by the Ewing sarcoma protein. Cell Rep 7, 1211–1226. 10.1016/j.celrep.2014.03.077.

31. Chen, K., Xiao, H., Zeng, J., Yu, G., Zhou, H., Huang, C., Yao, W., Xiao, W., Hu, J., Guan, W., et al. (2017). Alternative Splicing of EZH2 pre-mRNA by SF3B3 Contributes to the Tumorigenic Potential of Renal Cancer. Clin Cancer Res 23, 3428–3441. 10.1158/1078-0432.Ccr-16-2020.

32. Thomas, J.D., Polaski, J.T., Feng, Q., De Neef, E.J., Hoppe, E.R., McSharry, M.V., Pangallo, J., Gabel, A.M., Belleville, A.E., Watson, J., et al. (2020). RNA isoform screens uncover the essentiality and tumor-suppressor activity of ultraconserved poison exons. Nature Genetics 52, 84–94. 10.1038/s41588-019-0555-z.

33. Gonatopoulos-Pournatzis, T., Aregger, M., Brown, K.R., Farhangmehr, S., Braunschweig, U., Ward, H.N., Ha, K.C.H., Weiss, A., Billmann, M., Durbic, T., et al. (2020). Genetic interaction mapping and exon-resolution functional genomics with a hybrid Cas9–Cas12a platform. Nature Biotechnology 38, 638–648. 10.1038/s41587-020-0437-z.

34. Aregger, M., Xing, K., and Gonatopoulos-Pournatzis, T. (2021). Application of CHyMErA Cas9-Cas12a combinatorial genome-editing platform for genetic interaction mapping and gene fragment deletion screening. Nature Protocols 2021 16:10 16, 4722–4765. 10.1038/s41596-021-00595-1.

35. Ward, H.N., Aregger, M., Gonatopoulos-Pournatzis, T., Billmann, M., Ohsumi, T.K., Brown, K.R., Blencowe, B.J., Moffat, J., and Myers, C.L. (2021). Analysis of combinatorial CRISPR screens with the Orthrus scoring pipeline. Nature Protocols 2021 16:10 16, 4766–4798. 10.1038/s41596-021-00596-0.

36. Mu, W., Starmer, J., Yee, D., and Magnuson, T. (2018). EZH2 variants differentially regulate polycomb repressive complex 2 in histone methylation and cell differentiation. Epigenetics Chromatin 11, 71. 10.1186/s13072-018-0242-9.

37. Gier, R.A., Budinich, K.A., Evitt, N.H., Cao, Z., Freilich, E.S., Chen, Q., Qi, J., Lan, Y., Kohli, R.M., and Shi, J. (2020). High-performance CRISPR-Cas12a genome editing for combinatorial genetic screening. Nature communications 11, 3455–3455. 10.1038/s41467-020-17209-1.

38. Braun, S.E., Shi, X., Qiu, G., Wong, F.E., Joshi, P.J., Prasad, V.R., and Johnson, R.P. (2007). Instability of retroviral vectors with HIV-1-specific RT aptamers due to cryptic splice sites in the U6 promoter. AIDS Res Ther 4, 24. 10.1186/1742-6405-4-24.

39. Chen, M., David, C.J., and Manley, J.L. (2012). Concentration-dependent control of pyruvate kinase M mutually exclusive splicing by hnRNP proteins. Nat Struct Mol Biol 19, 346–354. 10.1038/nsmb.2219.

40. Wang, Z., Chatterjee, D., Jeon, H.Y., Akerman, M., Vander Heiden, M.G., Cantley, L.C., and Krainer, A.R. (2012). Exon-centric regulation of pyruvate kinase M alternative splicing via mutually exclusive exons. J Mol Cell Biol 4, 79–87. 10.1093/jmcb/mjr030.

41. Königs, V., de Oliveira Freitas Machado, C., Arnold, B., Blümel, N., Solovyeva, A., Löbbert, S., Schafranek, M., Ruiz De Los Mozos, I., Wittig, I., McNicoll, F., et al. (2020). SRSF7 maintains its homeostasis through the expression of Split-ORFs and nuclear body assembly. Nat Struct Mol Biol 27, 260–273. 10.1038/s41594-020-0385-9.

42. Lareau, L.F., Inada, M., Green, R.E., Wengrod, J.C., and Brenner, S.E. (2007). Unproductive splicing of SR genes associated with highly conserved and ultraconserved DNA elements. Nature 446, 926–929. 10.1038/nature05676.

43. Leclair, N.K., Brugiolo, M., Urbanski, L., Lawson, S.C., Thakar, K., Yurieva, M., George, J., Hinson, J.T., Cheng, A., Graveley, B.R., and Anczuków, O. (2020). Poison Exon Splicing Regulates a Coordinated Network of SR Protein Expression during Differentiation and Tumorigenesis. Molecular Cell 80, 648–665.e649. 10.1016/j.molcel.2020.10.019.

44. Karousis, E.D., and Mühlemann, O. (2019). Nonsense-Mediated mRNA Decay Begins Where Translation Ends. Cold Spring Harb Perspect Biol 11. 10.1101/cshperspect.a032862.

45. Kishor, A., Fritz, S.E., and Hogg, J.R. (2019). Nonsense-mediated mRNA decay: The challenge of telling right from wrong in a complex transcriptome. Wiley Interdiscip Rev RNA 10, e1548. 10.1002/wrna.1548.

46. Kurosaki, T., Popp, M.W., and Maquat, L.E. (2019). Quality and quantity control of gene expression by nonsense-mediated mRNA decay. Nat Rev Mol Cell Biol 20, 406–420. 10.1038/s41580-019-0126-2.

47. Loerch, S., Maucuer, A., Manceau, V., Green, M.R., and Kielkopf, C.L. (2014). Cancer-relevant splicing factor CAPERα engages the essential splicing factor SF3b155 in a specific ternary complex. J Biol Chem 289, 17325–17337. 10.1074/jbc.M114.558825.

48. Stepanyuk, G.A., Serrano, P., Peralta, E., Farr, C.L., Axelrod, H.L., Geralt, M., Das, D., Chiu, H.J., Jaroszewski, L., Deacon, A.M., et al. (2016). UHM-ULM interactions in the RBM39-U2AF65 splicing-factor complex. Acta Crystallogr D Struct Biol 72, 497–511. 10.1107/s2059798316001248.

49. Wang, E., Lu, S.X., Pastore, A., Chen, X., Imig, J., Chun-Wei Lee, S., Hockemeyer, K., Ghebrechristos, Y.E., Yoshimi, A., Inoue, D., et al. (2019). Targeting an RNA-Binding Protein Network in Acute Myeloid Leukemia. Cancer Cell 35, 369–384.e367. 10.1016/j.ccell.2019.01.010.

50. Haque, N., Will, A., Cook, A.G., and Hogg, J.R. (2023). A network of DZF proteins controls alternative splicing regulation and fidelity. Nucleic Acids Res 51, 6411–6429. 10.1093/nar/gkad351.

51. Kataoka, N., Bachorik, J.L., and Dreyfuss, G. (1999). Transportin-SR, a nuclear import receptor for SR proteins. J Cell Biol 145, 1145–1152. 10.1083/jcb.145.6.1145.

52. Lai, M.C., Lin, R.I., and Tarn, W.Y. (2001). Transportin-SR2 mediates nuclear import of phosphorylated SR proteins. Proc Natl Acad Sci U S A 98, 10154–10159. 10.1073/pnas.181354098.

53. Han, H., Braunschweig, U., Gonatopoulos-Pournatzis, T., Weatheritt, R.J., Hirsch, C.L., Ha, K.C.H., Radovani, E., Nabeel-Shah, S., Sterne-Weiler, T., Wang, J., et al. (2017). Multilayered Control of Alternative Splicing Regulatory Networks by Transcription Factors. Mol Cell 65, 539–553.e537. 10.1016/j.molcel.2017.01.011.

54. Sigova, A.A., Abraham, B.J., Ji, X., Molinie, B., Hannett, N.M., Guo, Y.E., Jangi, M., Giallourakis, C.C., Sharp, P.A., and Young, R.A. (2015). Transcription factor trapping by RNA in gene regulatory elements. Science 350, 978–981. 10.1126/science.aad3346.

55. Oksuz, O., Henninger, J.E., Warneford-Thomson, R., Zheng, M.M., Erb, H., Vancura, A., Overholt, K.J., Hawken, S.W., Banani, S.F., Lauman, R., et al. (2023). Transcription factors interact with RNA to regulate genes. Mol Cell 83, 2449–2463.e2413. 10.1016/j.molcel.2023.06.012.

56. Hendrickson, G.D., Kelley, D.R., Tenen, D., Bernstein, B., and Rinn, J.L. (2016). Widespread RNA binding by chromatin-associated proteins. Genome Biol 17, 28. 10.1186/s13059-016-0878-3.

57. Nabeel-Shah, S., Pu, S., Burns, J.D., Braunschweig, U., Ahmed, N., Burke, G.L., Lee, H., Radovani, E., Zhong, G., Tang, H., et al. (2024). C2H2-zinc-finger transcription factors bind RNA and function in diverse post-transcriptional regulatory processes. Mol Cell 84, 3810–3825.e3810. 10.1016/j.molcel.2024.08.037.

58. Takahashi, H., Takigawa, I., Watanabe, M., Anwar, D., Shibata, M., Tomomori-Sato, C., Sato, S., Ranjan, A., Seidel, C.W., Tsukiyama, T., et al. (2015). MED26 regulates the transcription of snRNA genes through the recruitment of little elongation complex. Nat Commun 6, 5941. 10.1038/ncomms6941.

59. Guiro, J., and Murphy, S. (2017). Regulation of expression of human RNA polymerase II-transcribed snRNA genes. Open Biol 7. 10.1098/rsob.170073.

60. Jiang, H., He, X., Wang, S., Jia, J., Wan, Y., Wang, Y., Zeng, R., Yates, J., 3rd, Zhu, X., and Zheng, Y. (2014). A microtubule-associated zinc finger protein, BuGZ, regulates mitotic chromosome alignment by ensuring Bub3 stability and kinetochore targeting. Dev Cell 28, 268–281. 10.1016/j.devcel.2013.12.013.

61. Toledo, C.M., Herman, J.A., Olsen, J.B., Ding, Y., Corrin, P., Girard, E.J., Olson, J.M., Emili, A., DeLuca, J.G., and Paddison, P.J. (2014). BuGZ is required for Bub3 stability, Bub1 kinetochore function, and chromosome alignment. Dev Cell 28, 282–294. 10.1016/j.devcel.2013.12.014.

62. Fang, F., Xia, N., Angulo, B., Carey, J., Cady, Z., Durruthy-Durruthy, J., Bennett, T., Sebastiano, V., and Reijo Pera, R.A. (2018). A distinct isoform of ZNF207 controls self-renewal and pluripotency of human embryonic stem cells. Nat Commun 9, 4384. 10.1038/s41467-018-06908-5.

63. Malla, S., Prasad Bhattarai, D., Groza, P., Melguizo-Sanchis, D., Atanasoai, I., Martinez-Gamero, C., Román Á, C., Zhu, D., Lee, D.F., Kutter, C., and Aguilo, F. (2022). ZFP207 sustains pluripotency by coordinating OCT4 stability, alternative splicing and RNA export. EMBO Rep 23, e53191. 10.15252/embr.202153191.

64. Baltz, A.G., Munschauer, M., Schwanhäusser, B., Vasile, A., Murakawa, Y., Schueler, M., Youngs, N., Penfold-Brown, D., Drew, K., Milek, M., et al. (2012). The mRNA-bound proteome and its global occupancy profile on protein-coding transcripts. Mol Cell 46, 674–690. 10.1016/j.molcel.2012.05.021.

65. Castello, A., Fischer, B., Eichelbaum, K., Horos, R., Beckmann, B.M., Strein, C., Davey, N.E., Humphreys, D.T., Preiss, T., Steinmetz, L.M., et al. (2012). Insights into RNA biology from an atlas of mammalian mRNA-binding proteins. Cell 149, 1393–1406. 10.1016/j.cell.2012.04.031.

66. Hegele, A., Kamburov, A., Grossmann, A., Sourlis, C., Wowro, S., Weimann, M., Will, C.L., Pena, V., Lührmann, R., and Stelzl, U. (2012). Dynamic protein-protein interaction wiring of the human spliceosome. Mol Cell 45, 567–580. 10.1016/j.molcel.2011.12.034.

67. Wan, Y., Zheng, X., Chen, H., Guo, Y., Jiang, H., He, X., Zhu, X., and Zheng, Y. (2015). Splicing function of mitotic regulators links R-loop-mediated DNA damage to tumor cell killing. J Cell Biol 209, 235–246. 10.1083/jcb.201409073.

68. Cho, N.H., Cheveralls, K.C., Brunner, A.D., Kim, K., Michaelis, A.C., Raghavan, P., Kobayashi, H., Savy, L., Li, J.Y., Canaj, H., et al. (2022). OpenCell: Endogenous tagging for the cartography of human cellular organization. Science 375, eabi6983. 10.1126/science.abi6983.

69. Choi, Y., Um, B., Na, Y., Kim, J., Kim, J.S., and Kim, V.N. (2024). Time-resolved profiling of RNA binding proteins throughout the mRNA life cycle. Mol Cell 84, 1764–1782.e1710. 10.1016/j.molcel.2024.03.012.

70. Wu, D., Yates, P.A., Zhang, H., and Cao, K. (2016). Comparing lamin proteins post-translational relative stability using a 2A peptide-based system reveals elevated resistance of progerin to cellular degradation. Nucleus 7, 585–596. 10.1080/19491034.2016.1260803.

71. Puttaraju, M., Jackson, M., Klein, S., Shilo, A., Bennett, C.F., Gordon, L., Rigo, F., and Misteli, T. (2021). Systematic screening identifies therapeutic antisense oligonucleotides for Hutchinson-Gilford progeria syndrome. Nat Med 27, 526–535. 10.1038/s41591-021-01262-4.

72. Scaffidi, P., and Misteli, T. (2005). Reversal of the cellular phenotype in the premature aging disease Hutchinson-Gilford progeria syndrome. Nat Med 11, 440–445. 10.1038/nm1204.

73. Scaffidi, P., and Misteli, T. (2006). Lamin A-dependent nuclear defects in human aging. Science 312, 1059–1063. 10.1126/science.1127168.

74. Vidak, S., Kubben, N., Dechat, T., and Foisner, R. (2015). Proliferation of progeria cells is enhanced by lamina-associated polypeptide 2α (LAP2α) through expression of extracellular matrix proteins. Genes Dev 29, 2022–2036. 10.1101/gad.263939.115.

75. Tsherniak, A., Vazquez, F., Montgomery, P.G., Golub, T.R., Boehm, J.S., Hahn, W.C., Tsherniak, A., Vazquez, F., Montgomery, P.G., Weir, B.A., et al. (2017). Defining a Cancer Dependency Map. Cell 170, 564–570.e516. 10.1016/j.cell.2017.06.010.

76. Beusch, I., Rao, B., Studer, M.K., Luhovska, T., Šukytė, V., Lei, S., Oses-Prieto, J., SeGraves, E., Burlingame, A., Jonas, S., and Madhani, H.D. (2023). Targeted high-throughput mutagenesis of the human spliceosome reveals its in vivo operating principles. Mol Cell 83, 2578–2594.e2579. 10.1016/j.molcel.2023.06.003.

77. Hanna, R.E., Hegde, M., Fagre, C.R., DeWeirdt, P.C., Sangree, A.K., Szegletes, Z., Griffith, A., Feeley, M.N., Sanson, K.R., Baidi, Y., et al. (2021). Massively parallel assessment of human variants with base editor screens. Cell 184, 1064–1080.e1020. 10.1016/j.cell.2021.01.012.

78. Cuella-Martin, R., Hayward, S.B., Fan, X., Chen, X., Huang, J.-W., Taglialatela, A., Leuzzi, G., Zhao, J., Rabadan, R., Lu, C., et al. (2021). Functional interrogation of DNA damage response variants with base editing screens. Cell 184, 1081–1097.e1019. 10.1016/j.cell.2021.01.041.

79. Jiang, H., Wang, S., Huang, Y., He, X., Cui, H., Zhu, X., and Zheng, Y. (2015). Phase transition of spindle-associated protein regulate spindle apparatus assembly. Cell 163, 108–122. 10.1016/j.cell.2015.08.010.

80. Jin, W., Brannan, K.W., Kapeli, K., Park, S.S., Tan, H.Q., Gosztyla, M.L., Mujumdar, M., Ahdout, J., Henroid, B., Rothamel, K., et al. (2023). HydRA: Deep-learning models for predicting RNA-binding capacity from protein interaction association context and protein sequence. Mol Cell 83, 2595–2611.e2511. 10.1016/j.molcel.2023.06.019.

81. Evers, B., Jastrzebski, K., Heijmans, J.P.M., Grernrum, W., Beijersbergen, R.L., and Bernards, R. (2016). CRISPR knockout screening outperforms shRNA and CRISPRi in identifying essential genes. Nature biotechnology 34, 631–634. 10.1038/nbt.3536.

82. Muller, R., Meacham, Z.A., Ferguson, L., and Ingolia, N.T. (2020). CiBER-seq dissects genetic networks by quantitative CRISPRi profiling of expression phenotypes. Science 370. 10.1126/science.abb9662.

83. Alford, B.D., Tassoni-Tsuchida, E., Khan, D., Work, J.J., Valiant, G., and Brandman, O. (2021). ReporterSeq reveals genome-wide dynamic modulators of the heat shock response across diverse stressors. Elife 10. 10.7554/eLife.57376.

84. Gonatopoulos-Pournatzis, T., Wu, M., Braunschweig, U., Roth, J., Han, H., Best, A.J., Raj, B., Aregger, M., O’Hanlon, D., Ellis, J.D., et al. (2018). Genome-wide CRISPR-Cas9 Interrogation of Splicing Networks Reveals a Mechanism for Recognition of Autism-Misregulated Neuronal Microexons. Molecular Cell 72, 510–524.e512. 10.1016/j.molcel.2018.10.008.

85. Benbarche, S., Pineda, J.M.B., Galvis, L.B., Biswas, J., Liu, B., Wang, E., Zhang, Q., Hogg, S.J., Lyttle, K., Dahi, A., et al. (2024). GPATCH8 modulates mutant SF3B1 mis-splicing and pathogenicity in hematologic malignancies. Mol Cell. 10.1016/j.molcel.2024.04.006.

86. Scarborough, A.M., Flaherty, J.N., Hunter, O.V., Liu, K., Kumar, A., Xing, C., Tu, B.P., and Conrad, N.K. (2021). SAM homeostasis is regulated by CFI(m)-mediated splicing of MAT2A. Elife 10. 10.7554/eLife.64930.

87. Lopez-Mejia, I.C., Vautrot, V., De Toledo, M., Behm-Ansmant, I., Bourgeois, C.F., Navarro, C.L., Osorio, F.G., Freije, J.M., Stévenin, J., De Sandre-Giovannoli, A., et al. (2011). A conserved splicing mechanism of the LMNA gene controls premature aging. Hum Mol Genet 20, 4540–4555. 10.1093/hmg/ddr385.

88. Jourdain, A.A., Begg, B.E., Mick, E., Shah, H., Calvo, S.E., Skinner, O.S., Sharma, R., Blue, S.M., Yeo, G.W., Burge, C.B., and Mootha, V.K. (2021). Loss of LUC7L2 and U1 snRNP subunits shifts energy metabolism from glycolysis to OXPHOS. Mol Cell 81, 1905–1919.e1912. 10.1016/j.molcel.2021.02.033.

89. Daniels, N.J., Hershberger, C.E., Gu, X., Schueger, C., DiPasquale, W.M., Brick, J., Saunthararajah, Y., Maciejewski, J.P., and Padgett, R.A. (2021). Functional analyses of human LUC7-like proteins involved in splicing regulation and myeloid neoplasms. Cell Rep 35, 108989. 10.1016/j.celrep.2021.108989.

90. Kenny, C.J., McGurk, M.P., Schüler, S., Cordero, A., Laubinger, S., and Burge, C.B. (2024). LUC7 proteins define two major classes of 5’ splice sites in animals and plants. bioRxiv, 2022.2012.2007.519539. 10.1101/2022.12.07.519539.

91. Koblan, L.W., Erdos, M.R., Wilson, C., Cabral, W.A., Levy, J.M., Xiong, Z.M., Tavarez, U.L., Davison, L.M., Gete, Y.G., Mao, X., et al. (2021). In vivo base editing rescues Hutchinson-Gilford progeria syndrome in mice. Nature 589, 608–614. 10.1038/s41586-020-03086-7.

92. Gosztyla, M.L., Zhan, L., Olson, S., Wei, X., Naritomi, J., Nguyen, G., Street, L., Goda, G.A., Cavazos, F.F., Schmok, J.C., et al. (2024). Integrated multi-omics analysis of zinc-finger proteins uncovers roles in RNA regulation. Mol Cell 84, 3826–3842.e3828. 10.1016/j.molcel.2024.08.010.

93. Brayer, K.J., and Segal, D.J. (2008). Keep your fingers off my DNA: protein-protein interactions mediated by C2H2 zinc finger domains. Cell Biochem Biophys 50, 111–131. 10.1007/s12013-008-9008-5.

94. Hart, T., Tong, A.H.Y., Chan, K., Van Leeuwen, J., Seetharaman, A., Aregger, M., Chandrashekhar, M., Hustedt, N., Seth, S., Noonan, A., et al. (2017). Evaluation and Design of Genome-Wide CRISPR/SpCas9 Knockout Screens. G3: Genes, Genomes, Genetics 7, g3.117.041277–g041273.041117.041277. 10.1534/g3.117.041277.

95. Chen, E.Y., Tan, C.M., Kou, Y., Duan, Q., Wang, Z., Meirelles, G.V., Clark, N.R., and Ma’ayan, A. (2013). Enrichr: interactive and collaborative HTML5 gene list enrichment analysis tool. BMC Bioinformatics 14, 128. 10.1186/1471-2105-14-128.

96. Jumper, J., Evans, R., Pritzel, A., Green, T., Figurnov, M., Ronneberger, O., Tunyasuvunakool, K., Bates, R., Žídek, A., Potapenko, A., et al. (2021). Highly accurate protein structure prediction with AlphaFold. Nature 596, 583–589. 10.1038/s41586-021-03819-2.

97. Varadi, M., Anyango, S., Deshpande, M., Nair, S., Natassia, C., Yordanova, G., Yuan, D., Stroe, O., Wood, G., Laydon, A., et al. (2022). AlphaFold Protein Structure Database: massively expanding the structural coverage of protein-sequence space with high-accuracy models. Nucleic Acids Res 50, D439–d444. 10.1093/nar/gkab1061.

98. Kolberg, L., Raudvere, U., Kuzmin, I., Adler, P., Vilo, J., and Peterson, H. (2023). g:Profiler-interoperable web service for functional enrichment analysis and gene identifier mapping (2023 update). Nucleic Acids Res 51, W207–w212. 10.1093/nar/gkad347.

99. Shannon, P., Markiel, A., Ozier, O., Baliga, N.S., Wang, J.T., Ramage, D., Amin, N., Schwikowski, B., and Ideker, T. (2003). Cytoscape: a software environment for integrated models of biomolecular interaction networks. Genome Res 13, 2498–2504. 10.1101/gr.1239303.

100. Doncheva, N.T., Morris, J.H., Gorodkin, J., and Jensen, L.J. (2019). Cytoscape StringApp: Network Analysis and Visualization of Proteomics Data. J Proteome Res 18, 623–632. 10.1021/acs.jproteome.8b00702.

101. Van Nostrand, E.L., Nguyen, T.B., Gelboin-Burkhart, C., Wang, R., Blue, S.M., Pratt, G.A., Louie, A.L., and Yeo, G.W. (2017). Robust, Cost-Effective Profiling of RNA Binding Protein Targets with Single-end Enhanced Crosslinking and Immunoprecipitation (seCLIP). Methods Mol Biol 1648, 177–200. 10.1007/978-1-4939-7204-3_14.

102. Ewels, P., Magnusson, M., Lundin, S., and Käller, M. (2016). MultiQC: summarize analysis results for multiple tools and samples in a single report. Bioinformatics 32, 3047–3048. 10.1093/bioinformatics/btw354.

103. Smith, T., Heger, A., and Sudbery, I. (2017). UMI-tools: modeling sequencing errors in Unique Molecular Identifiers to improve quantification accuracy. Genome Res 27, 491–499. 10.1101/gr.209601.116.

104. Martin, M. (2011). Cutadapt removes adapter sequences from high-throughput sequencing reads. EMBnet.journal 17, 3. 10.14806/ej.17.1.200.

105. Dobin, A., Davis, C.A., Schlesinger, F., Drenkow, J., Zaleski, C., Jha, S., Batut, P., Chaisson, M., and Gingeras, T.R. (2013). STAR: ultrafast universal RNA-seq aligner. Bioinformatics 29, 15–21. 10.1093/bioinformatics/bts635.

106. Li, H., Handsaker, B., Wysoker, A., Fennell, T., Ruan, J., Homer, N., Marth, G., Abecasis, G., and Durbin, R. (2009). The Sequence Alignment/Map format and SAMtools. Bioinformatics 25, 2078–2079. 10.1093/bioinformatics/btp352.

107. Quinlan, A.R., and Hall, I.M. (2010). BEDTools: a flexible suite of utilities for comparing genomic features. Bioinformatics 26, 841–842. 10.1093/bioinformatics/btq033.

108. Anders, S., Pyl, P.T., and Huber, W. (2015). HTSeq--a Python framework to work with high-throughput sequencing data. Bioinformatics 31, 166–169. 10.1093/bioinformatics/btu638.

109. Schwarzl, T., Sahadevan, S., Lang, B., Miladi, M., Backofen, R., Huber, W., Hentze, M.W., and Tartaglia, G.G. (2024). Improved discovery of RNA-binding protein binding sites in eCLIP data using DEWSeq. Nucleic Acids Res 52, e1. 10.1093/nar/gkad998.

110. Ramírez, F., Dündar, F., Diehl, S., Grüning, B.A., and Manke, T. (2014). deepTools: a flexible platform for exploring deep-sequencing data. Nucleic Acids Res 42, W187–191. 10.1093/nar/gku365.

111. Perez, A.R., Pritykin, Y., Vidigal, J.A., Chhangawala, S., Zamparo, L., Leslie, C.S., and Ventura, A. (2017). GuideScan software for improved single and paired CRISPR guide RNA design. Nature Biotechnology 35, 347–349. 10.1038/nbt.3804.

112. Perez, A.R., Sala, L., Perez, R.K., and Vidigal, J.A. (2021). CSC software corrects off-target mediated gRNA depletion in CRISPR-Cas9 essentiality screens. Nat Commun 12, 6461. 10.1038/s41467-021-26722-w.

113. Doench, J.G., Fusi, N., Sullender, M., Hegde, M., Vaimberg, E.W., Donovan, K.F., Smith, I., Tothova, Z., Wilen, C., Orchard, R., et al. (2016). Optimized sgRNA design to maximize activity and minimize off-target effects of CRISPR-Cas9. Nature Biotechnology 34, 184–191. 10.1038/nbt.3437.

114. Dede, M., McLaughlin, M., Kim, E., and Hart, T. (2020). Multiplex enCas12a screens detect functional buffering among paralogs otherwise masked in monogenic Cas9 knockout screens. Genome Biology 21, 262–262. 10.1186/s13059-020-02173-2.

115. Parrish, P.C.R., Thomas, J.D., Gabel, A.M., Kamlapurkar, S., Bradley, R.K., and Berger, A.H. (2021). Discovery of synthetic lethal and tumor suppressor paralog pairs in the human genome. Cell Reports 36, 109597–109597. 10.1016/J.CELREP.2021.109597.

116. Thompson, N.A., Ranzani, M., van der Weyden, L., Iyer, V., Offord, V., Droop, A., Behan, F., Gonçalves, E., Speak, A., Iorio, F., et al. (2021). Combinatorial CRISPR screen identifies fitness effects of gene paralogues. Nature Communications 12, 1302–1302. 10.1038/s41467-021-21478-9.

117. Dang, Y., Jia, G., Choi, J., Ma, H., Anaya, E., Ye, C., Shankar, P., and Wu, H. (2015). Optimizing sgRNA structure to improve CRISPR-Cas9 knockout efficiency. Genome Biology 16, 280–280. 10.1186/s13059-015-0846-3.

118. Love, M.I., Huber, W., and Anders, S. (2014). Moderated estimation of fold change and dispersion for RNA-seq data with DESeq2. Genome Biol 15, 550. 10.1186/s13059-014-0550-8.

119. Sterne-Weiler, T., Weatheritt, R.J., Best, A.J., Ha, K.C.H., and Blencowe, B.J. (2018). Efficient and Accurate Quantitative Profiling of Alternative Splicing Patterns of Any Complexity on a Laptop. Mol Cell 72, 187–200.e186. 10.1016/j.molcel.2018.08.018.

120. Merico, D., Isserlin, R., Stueker, O., Emili, A., and Bader, G.D. (2010). Enrichment map: a network-based method for gene-set enrichment visualization and interpretation. PLoS One 5, e13984. 10.1371/journal.pone.0013984.

121. Yu, G., Wang, L.G., Han, Y., and He, Q.Y. (2012). clusterProfiler: an R package for comparing biological themes among gene clusters. Omics 16, 284–287. 10.1089/omi.2011.0118.

